# MOFI reveals dynamic multi-omics coordination during cell fate progression

**DOI:** 10.64898/2026.10.09.757904

**Authors:** Jiantao Shen, Zhenyi Zhang, Hao Peng, Qiangwei Peng, Zihan Wang, Rui Liu, Peijie Zhou

**Affiliations:** Center for Machine Learning Research, Peking University, Beijing, 100871, China; AI for Science Institute, Beijing, 100080, China; School of Mathematics, South China University of Technology, Guangzhou, 510640, China; LMAM and School of Mathematical Sciences, Peking University, Beijing, 100871, China; School of Future Technology, South China University of Technology, Guangzhou, 511442, China; Center for Quantitative Biology, Peking University, Beijing, 100871, China; National Engineering Laboratory for Big Data Analysis and Applications, Beijing, 100871, China

## Abstract

Cell fate decisions are shaped by dynamic coordination across transcriptomic, chromatin, and protein layers. Time-series single-cell multi-omics assays now enable these molecular layers to be profiled across developmental stages, yet each cell is still observed only once, and the continuous multi-omics trajectories connecting discrete snapshots remain unresolved. A central challenge is therefore to jointly integrate paired omics measurements while reconstructing their shared developmental progression. Here we present Multi-Omics Fate Inference (MOFI), a dynamically constrained multi-omics integration framework designed to reconstruct continuous cell fate transitions from paired multi-omics snapshots sampled across multiple time points. MOFI jointly models omics-specific state dynamics, cross-omics correspondence, and shared population growth within a unified framework, enabling cross-omics perturbation response analysis and identification of candidate cross-omics transformation programs. Across simulated benchmarks and paired RNA–ATAC/RNA–protein time-series datasets, MOFI reconstructed continuous fate transitions, population dynamics and cross-omics changes consistent with observed measurements. In kidney organoids, MOFI captured coordinated RNA–ATAC progression of nephron lineage specification. In hematopoietic progenitors, MOFI predicted perturbation-responsive lineage redistribution and nominated multi-omics transformation programs linking transcriptional modules with protein-level responses. In neural developmental systems, MOFI resolved gradual brain organoid maturation and interneuron lineage allocation during human cortical development. Together, MOFI provides a dynamics-grounded method for converting discrete paired multi-omics snapshots into interpretable models of cell fate progression, perturbation response, and cross-omics transformation analysis.

## Introduction

Single-cell multi-omics technologies have transformed the study of cell fate decisions by enabling multiple molecular layers to be measured within the same biological system[1][2]. Joint RNA–chromatin profiling, such as single-cell RNA–ATAC sequencing[3][4], connects transcriptional output with the regulatory landscape that shapes it, whereas RNA–protein assays such as CITE-seq[5] link gene expression programs to cell-surface and functional protein states. When collected across ordered developmental stages, these paired multi-omics measurements provide a rich temporal view of how progenitors acquire lineage identity through coordinated changes in gene expression, chromatin accessibility, protein abundance and population composition.

Despite this progress, time-series single-cell multi-omics data consist of destructive snapshots[6]. Each cell is measured only once, and the continuous paths connecting early progenitors to later differentiated states are not directly observed. This limitation is especially important in developmental systems, where cell fate is not defined solely by a static molecular state, but by a dynamic process involving differentiation, proliferation, depletion, lineage allocation, and molecular response. Therefore, fully exploiting time-series multi-omics data requires more than aligning omics at individual time points. It requires a method that uses developmental progression itself as a biological constraint to integrate multi-omics and reconstruct continuous fate dynamics from discrete paired observations.

Existing multi-omics integration methods have substantially advanced single-cell analysis by aligning molecular profiles, constructing shared representations and transferring information across transcriptomic, epigenomic and proteomic assays[7][8]. These approaches have enabled joint cell-state annotation, omics imputation, data harmonization and cross-omics comparison across diverse biological systems[9][10]. However, most integration frameworks still view cross-omics integration essentially as a static process[2]. Although they can align molecular states across omics, the absence of temporal constraints limits both the integration itself and the biological questions it can address. Consequently, under such methods, it is challenging to determine how integrated multi-omics states are connected across time, nor can they reconstruct the continuous developmental dynamics through which progenitors progress toward differentiated lineages. This leaves a central gap for cell fate inference: what is needed is not only cross-omics alignment, but dynamic multi-omics integration in which molecular correspondence is constrained by developmental progression.

To address the lack of temporal information, a variety of dynamic inference methods have been developed to reconstruct cellular progression from snapshot measurements[11]. Pseudotime[12], RNA velocity[13], neural ordinary differential equation models[14], dynamic optimal transport[15] and flow-based approaches[16][17] have provided tools for ordering cells, estimating directions of change and modeling population transport over time. Unbalanced optimal transport further allows cell abundance to vary, which is essential for modeling proliferation, apoptosis and lineage expansion during development[17][18]. However, these approaches are largely restricted to a single omics layer[11]. Although they can recover developmental dynamics within one molecular space, such trajectories provide only a partial view of a developmental process that is jointly manifested across transcriptional, chromatin and protein states. Therefore, a unified framework is needed to treat cross-omics correspondence and developmental progression as a coupled dynamic inference problem.

Recent work has begun to connect multi-omics integration with dynamic inference tasks. MultiVelo[19] models coordinated RNA and chromatin changes through velocity-related assumptions, providing an important step toward coupled RNA–chromatin dynamics, but it is primarily designed for snapshot RNA–ATAC settings. scMultiNODE[20] learns dynamics in a shared latent representation, which is useful for connecting different omics datasets; however, reconstructing developmental dynamics after compressing diverse omics into a common space can make it difficult to preserve the biological changes observed in each omics and to recover the underlying multi-omics developmental process. Moscot[21] provides a flexible optimal transport framework for temporal, spatial, and cross-omics alignment tasks. While it focuses on establishing alignment between observed snapshots, the continuous dynamics are not explicitly generated. Thus, paired time-series multi-omics fate inference still requires a framework that jointly models cross-omics correspondence, population changes, and cellular dynamics within the same developmental process.

Here we introduce MOFI (Multi-Omics Fate Inference), a dynamically constrained integration framework for paired time-series single-cell multi-omics data. Resolving cell-fate transitions across molecular layers requires multi-omics integration and dynamic inference to be modeled jointly. MOFI therefore provides an interpretable view for understanding cell-fate decisions from complementary molecular perspectives. Inspired by synchronized unbalanced optimal transport theory [22], MOFI reconstructs continuous fate trajectories and relative population changes while preserving cross-omics correspondence and omics-specific variation. This design enables cell trajectory reconstruction, lineage allocation analysis, in silico perturbation, and candidate cross-omics transformation programs within one unified developmental model.

We demonstrate MOFI across simulated systems with known ground truth and diverse single-cell multi-omics datasets spanning RNA–chromatin and RNA–protein measurements. In simulations, MOFI recovers branching trajectories and growth patterns from discrete multi-omics snapshots. In kidney organoid differentiation[23], MOFI reconstructs stepwise nephron lineage progression and coordinated RNA–ATAC marker dynamics, including *HNF1B, EYA1* and *PAX8*. In hematopoietic stem and progenitor cells[24], MOFI predicts perturbation-responsive fate redistribution toward or away from the MkP (megakaryocyte progenitor) lineage and identifies mast-cell-associated cross-omics transformation programs. In human brain organoids[25], MOFI resolves gradual maturation from embryoid-body-like states toward neuronal populations and reconstructs coordinated RNA–ATAC dynamics. In the developing human cerebral cortex[26], MOFI constructs lineage progression and predicts *ERBB4*-associated changes in interneuron lineage allocation. Collectively, these analyses show that dynamically constrained multi-omics integration can convert discrete paired omics snapshots into interpretable models of cell fate progression, perturbation response, and cross-omics transformation analysis.

## Results

### MOFI reconstructs continuous multi-omics dynamics from discrete paired snapshots

MOFI is designed for paired multi-omics measurements collected at ordered experimental time points, such as matched RNA expression and chromatin accessibility or protein abundance measured in the same cells. Given these discrete snapshots, MOFI reconstructs continuous cell-fate trajectories together with changes in molecular states across omics and in population composition (Figure 1A). This formulation treats paired molecular measurements as complementary views of the same developmental process while retaining omics-specific features (Figure 1B).

**Figure 1:**
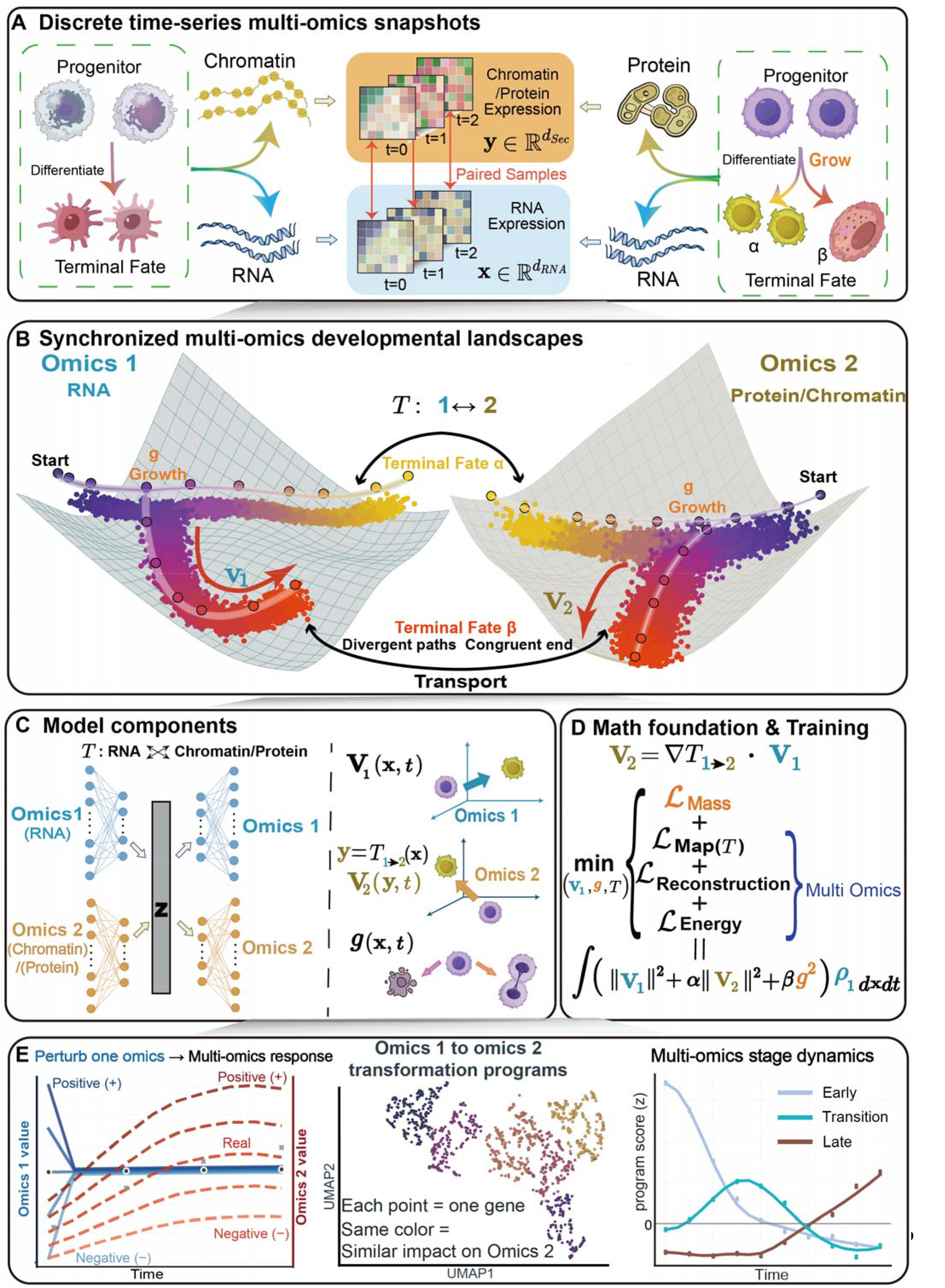
MOFI reconstructs continuous multi-omics cell-fate trajectories from discrete paired snapshots. (A) Input discrete multi-omics snapshots. Paired multi-omics data (e.g., RNA expression and chromatin accessibility/protein levels) collected at separate time points serve as input. MOFI infers the continuous evolution of cell populations across omics between these sampled time points. (B) Synchronized multi-omics developmental landscapes. Illustration of progenitor cell differentiation and proliferation toward distinct terminal fates across different omics spaces, capturing both shared fate transition programs and omics-specific dynamics. (C) Model framework. MOFI decomposes developmental processes into biologically interpretable components: bidirectional cross-omics map ***T***^**1→1**^, ***T***^1→2^, ***T***^**2→1**^, ***T***^**2→2**^,,that align cell states between two omics; velocity fields **v**_1_and **v**_2_ that describe cell-state transitions; and a growth rate ***g*** that models cell proliferation and death. Integration of multi-omics information generates omics-specific cell-state dynamics. (D) Training strategy. The model is optimized by matching predicted cell distributions to observed multi-omics populations at each time point. The objective integrates trajectory **ℒ**_reconstruction_. **ℒ**_Reconstruction_, cell-abundance matching (**ℒ**_Mass_), cross-omics map (**ℒ**_Map_), and energy regularization. **ℒ**_Energy_. (E) Biological downstream applications. The continuous model enables reconstruction of expression trajectories under native and perturbed conditions; identification of genes with similar cross-omics transformation profiles; and characterization of stage-specific dynamics of key developmental markers.

To achieve this objective, MOFI decomposes the developmental process into three interpretable components: (i) a cross-omics map **T** relates paired molecular states across multi-omics; (ii) velocity fields **v**_1_(**x**,*t*) and **v**_2_(**x**,*t*)describe time-dependent changes in multi-omics state; and (iii) a growth rate ***g***(**x**,*t*) captures the relative expansion or depletion of cell states during differentiation (Figure 1C). MOFI learns these components jointly by matching cell distributions observed at successive time points, while maintaining temporal and cross-omics coherence (Figure 1D). MOFI therefore provides an integrated representation of molecular dynamics, developmental progression, and population change during cell-fate transitions.

MOFI supports multiple downstream biological applications. First, MOFI interpolates continuous cell trajectories from discrete snapshots and simulates the predicted consequences of in silico perturbations applied to genes, proteins, or both omics (Figure 1E, left). These analyses trace how a modeled perturbation is associated with coordinated responses across multi-omics. Second, MOFI derives cross-omics transformation programs from the learned cross-omics map and velocity field. These programs quantify how changes in one omics are associated with predicted responses in the other omics and can be used to group genes with similar cross-omics transformation profiles (Figure 1E, middle). Third, the reconstructed continuous trajectories facilitate comprehensive profiling of the dynamic patterns of core lineage markers across multi-omics (e.g., transcriptome and chromatin accessibility) throughout early, transitional, and late cell states, uncovering coordinated changes in these molecular features during cell fate transitions (Figure 1E, right).

### MOFI recovers known dynamics in a simulation benchmark

Before applying MOFI to real-world data, we first assessed whether the framework could recover known dynamical structure from a controlled simulation in which the ground truth is fully specified. The simulation dataset was generated from a regulatory network defining causal relationships among variables **X, Y**, and **I**, governed by a state-dependent potential landscape **Z** that contains both attracting and repulsive regions (Figure 2A). The system was designed to produce two distinct lineages, with the repulsive potential field (dark blue circle in the landscape) acting as a factor shaping lineage transition trajectories. Multi-omics snapshots were sampled at discrete time points, with omics-1 capturing the **X**, **Y** plane and omics-2 capturing **X**, **Y**, **Z**, thereby mimicking the scenario in which omics-2 carries additional molecular dimensions that constrain the underlying dynamics (Figure 2B–C). Importantly, the simulation employed different numbers of sampling points across the two omics, with five time points for omics-1 and three for omics-2, to mirror realistic experimental constraints in which transcriptomic profiling is relatively inexpensive and can be performed at many time points, while proteomic or chromatin accessibility measurements are more costly and typically fewer. MOFI reconstructed the continuous dynamics from these discrete snapshots (Figure 2D), recovering both the velocity field structure in the omics-1 space and the pushforward-constrained trajectories in the omics-2 space. The predicted growth rates matched the ground-truth proliferation and depletion patterns across the state manifold (Figure 2E), confirming that MOFI distinguishes proliferative and non-proliferative regions. We next performed hold-one-out experiments to determine whether MOFI could recover observations excluded from training (Figure 2F). Individual time points were withheld in turn from both omics, and selected omics observations were additionally removed at specific time points. MOFI consistently achieved lower Wasserstein-1 distances than Moscot[21] and generally lower errors than the direct concatenation baseline adapted from stVCR[27] when reconstructing the hold-out distributions. Notably, the trajectories reconstructed by the direct concatenation baseline and Moscot deviate from the ground-truth dynamics (Figure 2G). The method for generating simulation data and detailed hold-one-out comparison are provided in Supplementary Note 1. Overall, these results demonstrate that MOFI can reconstruct hold-out multi-omics states while preserving the underlying velocity, growth, and trajectory structure when the cross-omics map is known.

**Figure 2:**
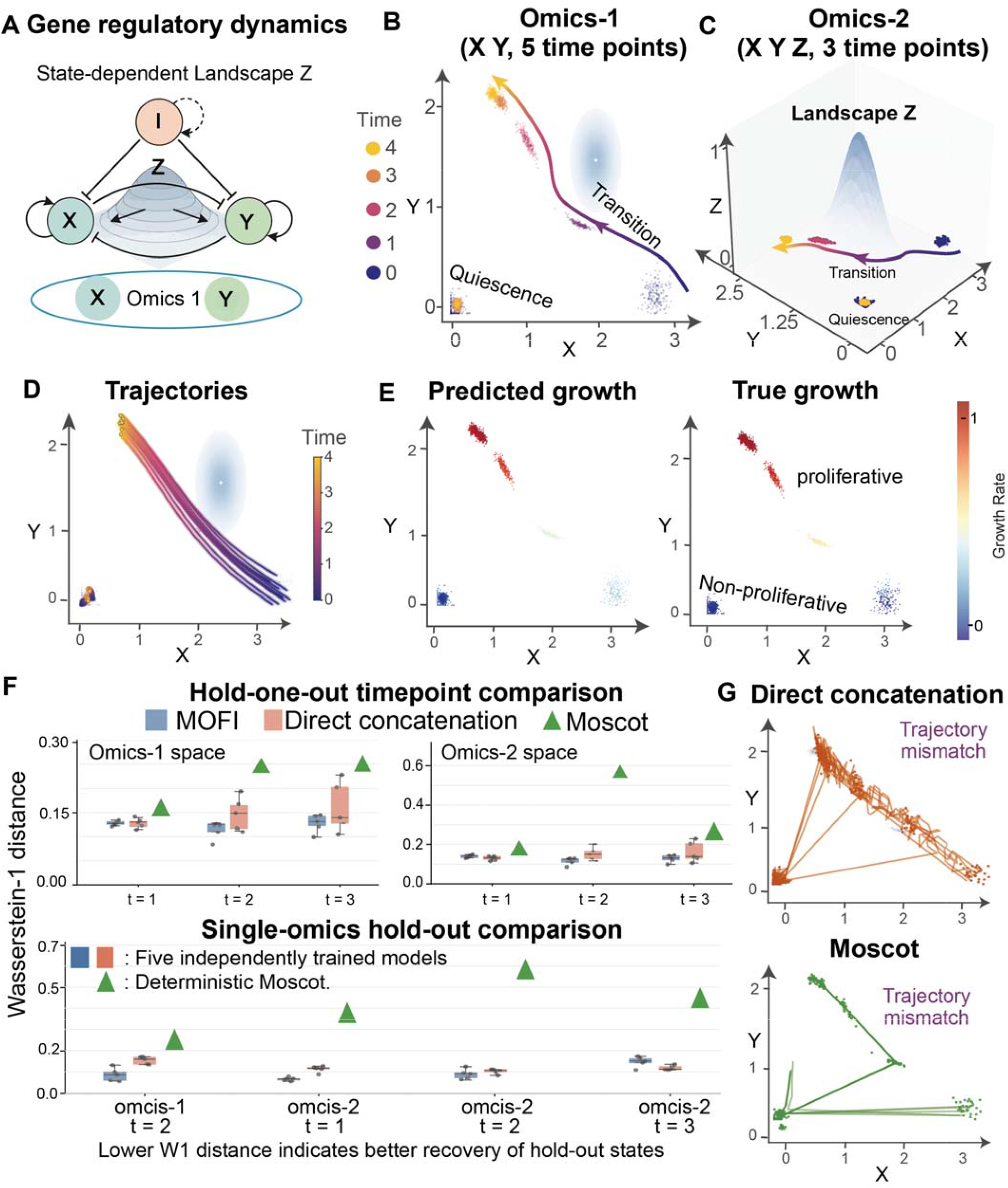
Validation of MOFI on a simulation benchmark. (A) Schematic of the regulatory interactions used to generate simulation datasets. The network defines the causal relationships among variables X, Y, and I, governed by a state-dependent potential landscape Z. (B–C) Multi-omics snapshots with divergent dynamics. Temporal snapshots of cell evolution in (B) omics-1 (X, Y) and (C) omics-2 (X, Y, Z). The system captures two distinct lineages. In the landscape, the dark blue circle represents a repulsive potential field. (D) Dynamics learned by MOFI. (E) Evaluation of growth rate estimation. Comparison of the growth rates inferred by MOFI (left) and the ground truth (right). (F) Hold-one-out evaluation of trajectory reconstruction under two missing-data paradigms. MOFI (blue) and direct concatenation baseline (orange) are compared via Wasserstein-1 (W1) distance across five independently trained models (seeds 22–26); green triangles mark the deterministic Moscot pairwise optimal transport reference. Lower W1 indicates better distribution recovery. The top two panels share the same dual-omics hold-one-out design: one intermediate time point (t = 1, 2, 3) is removed from both omics during training. Top left. Evaluated in omics-1 (2D) space. MOFI outperforms direct concatenation baseline at t = 2 and t = 3, with near-equivalent performance at t = 1. Top right. Evaluated in omics-2 space. Bottom. Time-point hold-one-out restricted to a single omics, with the other omics fully observed across all time points. Two scenarios are tested: t = 2 evaluated in omics-1, t = 1–3 evaluated in omics-2. (G) Trajectories reconstructed by direct concatenation baseline and Moscot.

### MOFI resolves multi-lineage fate commitment in kidney organoid differentiation

To assess whether MOFI could reconstruct branched developmental progression while preserving coordinated RNA and chromatin dynamics in a real biological system, we analyzed a paired RNA–ATAC time-series dataset of kidney organoid differentiation[23]. Nephron progenitor cells (NPCs) derived from posterior intermediate mesoderm give rise to developing (Dev) intermediate states and subsequently to epithelial lineages including proximal tubule (PT), loop of Henle (LOH), distal nephron (DN), and differentiated podocytes (POD).

The analysis revealed coordinated dynamical patterns between RNA expression and chromatin accessibility. In the RNA space, the inferred velocity fields trace a developmental trajectory progressing from day 7 cell states toward the four terminal cell states at day 26, consistent with measurements collected at five experimental time points (Figure 3A, top). Similarly, the ATAC space recapitulates a comparable developmental progression, exhibiting complementary yet distinct dynamics relative to RNA. Notably, the synchronized transport lines bridging the two omics illustrate how MOFI preserves temporal coherence across omics while retaining omics-specific biological information.

**Figure 3:**
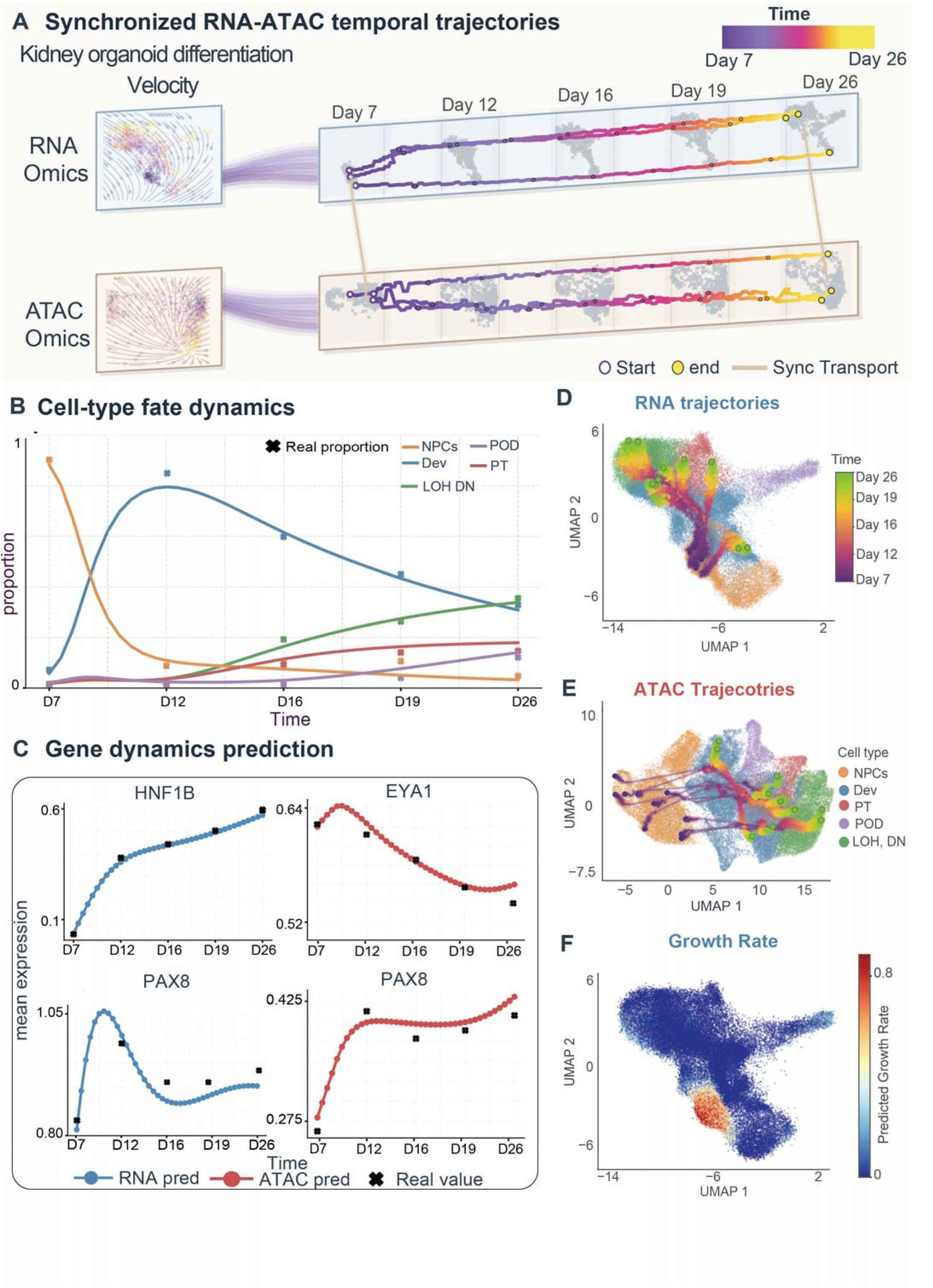
MOFI reconstructs the dynamic fate of kidney organoid differentiation. (A) Synchronized RNA–ATAC temporal trajectories. Multi-omics dynamic modeling of kidney organoid differentiation from Day 7 to Day 26. Left: Inferred multi-omics velocity fields for RNA and ATAC omics. Right: Continuous-time state trajectories simulated by MOFI, overlaid on discrete experimental snapshots (gray background). The lines highlight the temporal coupling between the two omics. (B) Cell-type fate dynamics. Predicted continuous cell-type proportions (lines) vs. observed discrete data (crosses), capturing the transition from NPCs to Dev to specialized lineages (PT, LOH, DN, and POD). (C) Gene and chromatin dynamics. Predicted profiles (lines) vs. observed experimental values (crosses) for marker genes (*HNF1B, EYA1*, and *PAX8*), showing high temporal fidelity across both RNA and ATAC omics. (D–E) UMAP state trajectories. UMAP visualizations of generated trajectories (colored by time) projected onto the observed data manifold for (D) RNA and (E) ATAC. (F) Predicted growth rates visualized on the RNA UMAP, representing the shared growth rate across omics.

Quantitative evaluation of cell-type compositional dynamics shows that MOFI recapitulates the temporal progression of lineage commitment during kidney organoid differentiation (Figure 3B). The predicted continuous trajectories match discrete experimental measurements across all time points: the NPC population progressively declines from nearly 100% at day 7 to below 10% by day 26 via lineage commitment into Dev. In turn, the Dev compartment is gradually depleted as it differentiates into the four terminal nephron lineages, which steadily accumulate over the course of development. Collectively, these dynamics reproduce the unidirectional, stepwise differentiation hierarchy observed in the original experiments[23]. Furthermore, quantitative reconstruction and hold⍰one⍰out comparison were performed across all real-world datasets to assess the recovery of experimentally unobserved intermediate states. MOFI was compared with scMultiNODE and a direct concatenation baseline, and the detailed results are presented in Supplementary Note 2 and Supplementary Table 3. At the individual gene level, MOFI demonstrates temporal fidelity for core kidney lineage markers (Figure 3C). MOFI reconstructed the temporal RNA expression of *HNF1B* and chromatin accessibility of *EYA1*, while the temporal profiles of *PAX8* were recovered in both RNA and ATAC. These results indicate that our framework not only reproduces global cell-type proportion shifts but also reconstructs temporal changes in molecular features during cell fate commitment.

An overlap was observed between the inferred trajectories and the experimentally profiled cells at matched developmental stages (Figure 3D–E). The inferred growth rate was highest in progenitor-enriched regions and decreased along differentiation paths (Figure 3F), consistent with the expected loss of growth rate as cells mature. Collectively, the kidney organoid analysis supports the use of synchronized RNA–ATAC dynamics for resolving branched developmental programs.

### In silico perturbation modeling and cross-omics transformation programs in hematopoietic stem and progenitor cells

To examine whether the learned multi-omics dynamics could support cross-omics analysis, we next analyzed a time-series RNA–protein dataset of human hematopoietic stem and progenitor cells (HSPCs) pairing RNA expression with protein-level measurements across days 2, 4, 7, and 10 (Figure 4). This system contains a well-characterized branch point at which hematopoietic stem cells (HSCs) commit to distinct fates, providing an opportunity to evaluate whether the learned dynamics can support perturbation-responsive fate analyses. MOFI generated intermediate snapshots for unobserved days 3, 5, 6, 8, and 9 (Figure 4A).

**Figure 4:**
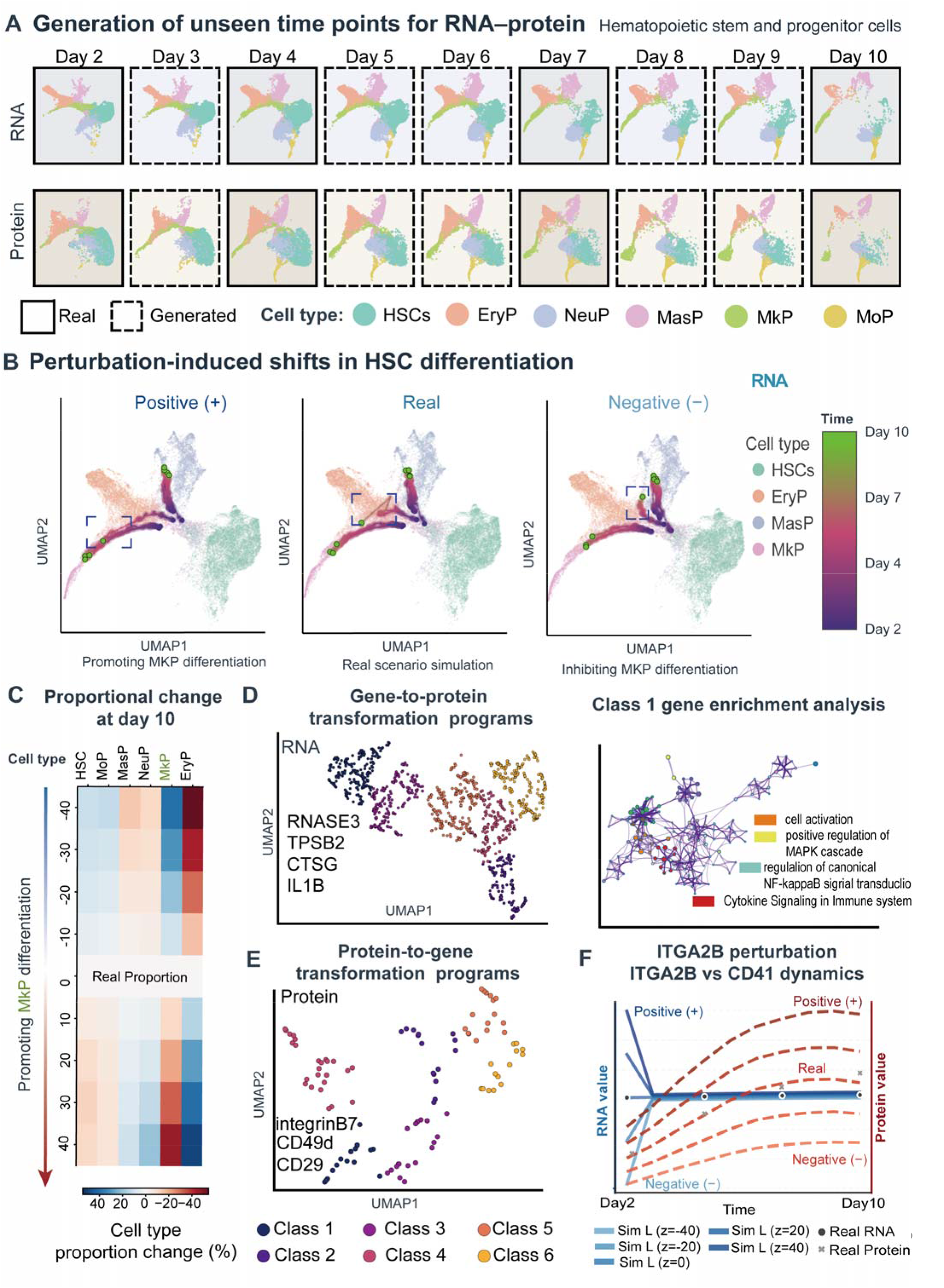
In silico perturbation modeling and cross-omics transformation analysis of HSPC differentiation. (A) Generation of unseen time points. MOFI generates continuous snapshots for Days 3, 5, 6, 8, and 9 (dashed boxes) by interpolating between experimental RNA–protein snapshots at Days 2, 4, 7, and 10 (solid boxes), capturing the fluid transition of hematopoietic stem and progenitor cells (HSPCs). Abbreviations: HSCs, hematopoietic stem cells; EryP, erythroid progenitor; NeuP, neutrophil progenitor; MasP, mast cell progenitor; MkP, megakaryocyte progenitor; MoP, monocyte progenitor. (B) Perturbation-induced shifts in HSCs fate trajectories. Simulation of HSCs differentiation under perturbations of MkP-associated genes. Positive (+) perturbation increased the predicted allocation of simulated HSC trajectories toward the MkP lineage, while negative (–) perturbation shifted the simulated HSCs trajectories away from MkP-associated states. (C) Proportion changes at Day 10. Heatmap illustrates cell-type composition changes under varying perturbation intensities of MkP-associated genes. (D) Gene-to-protein transformation programs. Left: UMAP embedding of the gene-to-protein transformation matrix. Genes are clustered into six classes (Classes 1–6) via k-nearest-neighbor (KNN) analysis based on their cross-omics transformation profiles, highlighting high-influence genes in class 1. Right: Functional enrichment network for class 1 genes. (E) Protein-to-gene transformation programs. Clustering of proteins based on protein-to-gene transformation profiles across transcriptomic states. (F) *ITGA2B* vs CD41 dynamics under *ITGA2B* perturbation. Predicted expression/abundance levels for *ITGA2B* (RNA) and CD41 (protein) under a gradient of RNA-level perturbations (z).

To test whether the learned dynamics captured the sensitivity of this branch point to targeted gene perturbation, we performed in silico perturbations of key MkP-associated genes including *ITGA2B, GP1BA, PF4*, and *PPBP*[28] and re-simulated the trajectories across a signed range of perturbation strengths. Under positive perturbation, the simulated HSC trajectories shifted more strongly toward the MkP lineage, whereas under negative perturbation the same initial cells showed an increased tendency toward alternative fates (Figure 4B). The fitted dynamics were therefore sensitive to the modeled perturbations around this branch point. A perturbation-intensity sweep further quantified the predicted day-10 redistribution of cell-type occupancy (Figure 4C). Consistent with the trajectory shifts observed in Figure 4B, the MkP fate expanded under the positive perturbation of MkP-associated genes, while the alternative erythroid progenitors (EryP) contracted. An opposite pattern was observed under inhibitory (negative) perturbation, demonstrating that MOFI captures systematic changes in modeled cell-fate allocation across transcriptomic perturbation strengths.

MOFI further characterizes cross-omics coordination by summarizing how individual molecular features are associated with predicted changes in the other omics along the learned dynamics. For each gene, we defined a gene-to-protein transformation profile in mast cell progenitors (MasP) as the pattern of predicted responses across measured proteins associated with local variation in that gene. Conversely, each protein was represented by a protein-to-gene profile describing its predicted associations with transcriptomic changes. We then compared genes and proteins according to the similarity of their respective transformation profiles. Projection of these profiles onto two-dimensional maps revealed distinct groups of molecular features with shared cross-omics transformation patterns (Figure 4D–E). These classes provide an entry point for exploring potential cross-omics coordination.

In the gene-to-protein transformation programs (Figure 4D), the highest-ranked class 1 included mast-cell and myeloid-associated genes such as *RNASE3, TPSB2, CTSG*, and *IL1B*. Functional enrichment analysis of these class 1 genes revealed an overrepresentation of pathways broadly implicated in mast-cell activation and function, including cell activation, cytokine signaling, and the canonical NF-κB and MAPK cascades (Figure 4D, right), consistent with established mast-cell biology[29]. Mirroring the transcriptomic findings, the protein-to-gene transformation programs (Figure 4E) highlighted surface receptors and adhesion molecules in class 1, including integrin β_7_ (ITGB7), CD49d (ITGA4), and CD29 (ITGB1). These integrins are documented to form functional heterodimers such as α_4_β_7_, that are critical for the homing, migration, and tissue-specific maturation of mast cell progenitors[30].

Finally, we evaluated whether MOFI could simulate the continuous, coordinated dynamics of a matched RNA–protein pair. We focused on the *ITGA2B*–CD41 axis, in which *ITGA2B* encodes integrin αIIb (CD41), a cell-surface receptor subunit expressed in megakaryocytes and platelets and upregulated during megakaryocytic differentiation[31]. The simulated *ITGA2B* perturbation produced graded and coordinated responses in both *ITGA2B* RNA and CD41 protein trajectories from day 2 to day 10 (Figure 4F), illustrating how MOFI can propagate perturbations across omics.

Taken together, the HSPCs analysis demonstrates that MOFI can interpolate unobserved RNA–protein states, simulate perturbation-dependent fate redistribution, and nominate putative cross-omics transformation programs for downstream validation.

### Human brain organoid maturation exhibits gradual compositional redistribution

Human brain organoid development proceeds as a continuous, temporally resolved maturation process, marked by gradual shifts in the relative proportions of embryoid body (EB) and post⍰mitotic neuronal populations, alongside the emergence of the neuroectoderm (nect) and its derivative neuroepithelium (nepi), over a two⍰month time course[25]. To assess whether MOFI can reconstruct such progressive compositional reweighting, we applied the framework to a time⍰series multi⍰omics dataset spanning days 4 to 61 (Figure 5).

**Figure 5:**
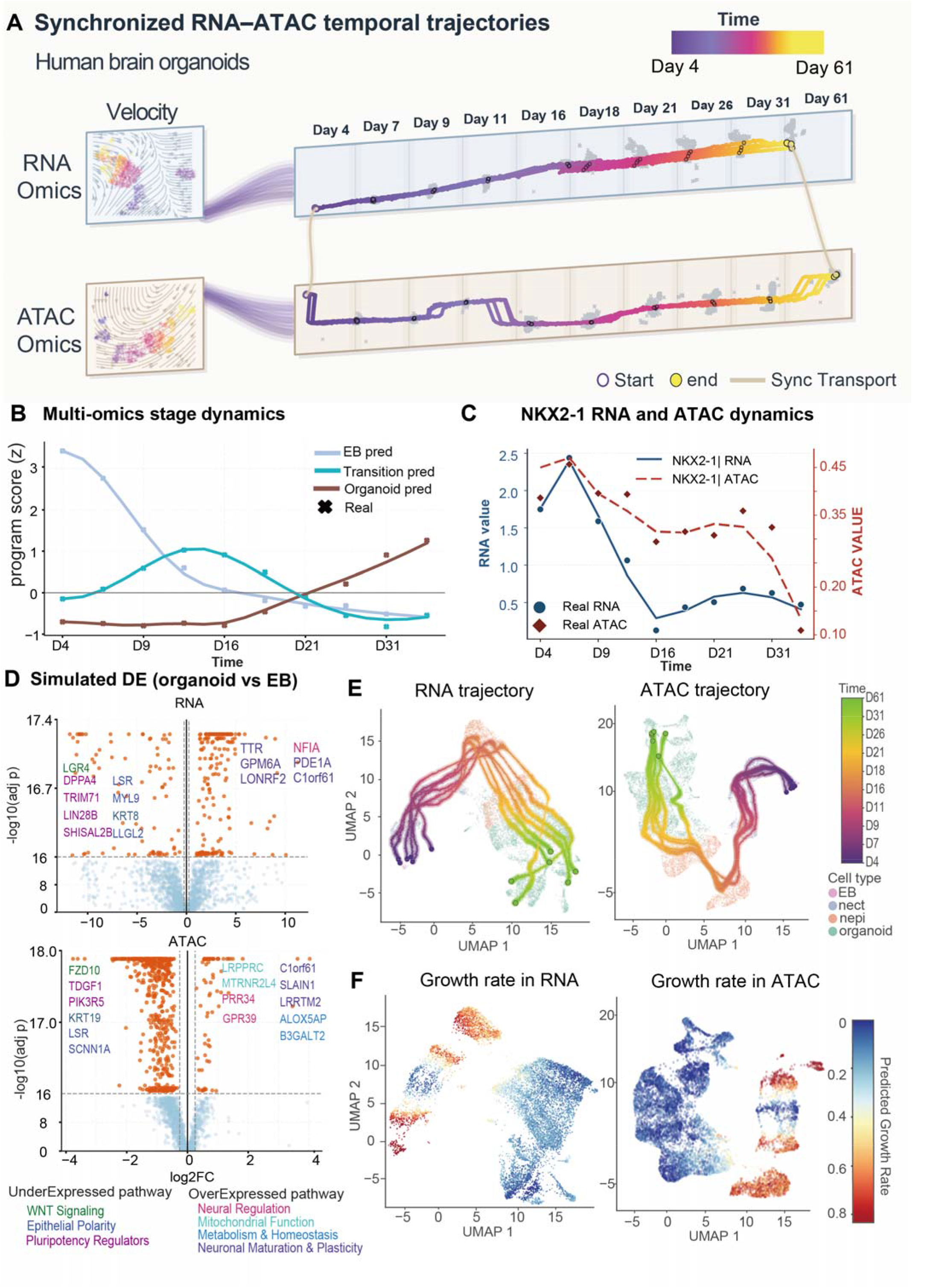
MOFI reconstructs the dynamic fate of human brain organoids. (A) Synchronized RNA–ATAC temporal trajectories. Multi-omics dynamic modeling of human brain organoid development from Day 4 to Day 61. Left: Inferred multi-omics velocity fields for RNA and ATAC omics. Right: Continuous-time state trajectories simulated by MOFI, overlaid on discrete experimental snapshots (gray background). (B) Multi-omics stage dynamics. Comparison of temporal program scores (z-scores) between real data (crosses) and MOFI reconstruction (lines). (C) Temporal dynamics for *NKX2-1* in RNA and ATAC. MOFI-predicted trajectories (lines) follow the experimental values for both RNA expression (blue circles) and chromatin accessibility (red diamonds). (D) Differential expression analysis: Organoid vs. EB. Volcano plots show simulated differential analysis in RNA (top) and ATAC (bottom) between the organoid and EB stages. Key genes are highlighted, with down-regulated (left) and up-regulated (right) markers corresponding to the functional pathways listed below. (E) Reconstructed state trajectories. Reconstructed continuous trajectories (lines) projected onto the experimental cell-type manifold for RNA (left) and ATAC (right). (F) Inferred multi-omics growth fields. Spatial distribution of predicted growth rates visualized on RNA (left) and ATAC (right) UMAPs.

MOFI-reconstructed temporal trajectories reveal a smooth progression across the full developmental axis (Figure 5A). Starting from EB states at day 4, cells sequentially transition through intermediate progenitor compartments before attaining later neuronal identity by day 61.

To assess whether the reconstructed molecular dynamics recapitulated observed cell-fate transitions during brain organoid development, we characterized temporal changes in core developmental programs (Figure 5B). Temporal program scores demonstrate gradual attenuation of early developmental signatures alongside concomitant activation of late-stage programs. This trend agreed with the experimental snapshot scores in both the transcriptomic and chromatin accessibility datasets. Furthermore, MOFI reconstructed the temporal dynamics of *NKX2-1* RNA expression and ATAC gene activity, which agreed with the experimentally observed values, thereby validating the model’s precision in reconstructing gene-specific dynamics (Figure 5C).

We next examined molecular features associated with this developmental progression through differential-feature analysis between generated organoid and EB stages (Figure 5D). Genes significantly upregulated in mature organoids are enriched in functional pathways related to Neuronal Maturation & Plasticity and Neural Regulation, including *NFIA*[32], a transcription factor implicated in neural progenitor differentiation and cortical development. In particular, *C1orf61*, which has been associated with early human cortical progenitor populations[33], was detected in both RNA and ATAC. In contrast, downregulated genes are predominantly implicated in Wnt signaling pathways, epithelial cell polarity maintenance, and pluripotency regulation. This pattern is consistent with progressive lineage restriction toward neural fates during organoid maturation. Projected trajectories showed continuous movement from progenitor-enriched to neuron-dominant regions in both RNA and ATAC embedding spaces (Figure 5E). Growth-rate inference localized higher growth rates to early progenitor regions and lower growth rates to late-stage states (Figure 5F). Collectively, these results demonstrate that MOFI enables joint modeling of temporal developmental progression and dynamic molecular coordination across multi-omics layers.

### Continuous corticogenesis with perturbation-responsive lineage allocation in the developing human cerebral cortex

The developing human cerebral cortex[26] provides a long developmental time series with paired RNA–ATAC measurements and multiple progenitor and neuronal populations, allowing us to assess whether MOFI can reconstruct lineage progression over an extended temporal interval while preserving omics-specific structure. We applied MOFI to a single-cell multi-omics atlas spanning gestational weeks 18 to 52 to test whether the framework could resolve both shared developmental progression and omics-specific structure.

The trajectories reconstructed by MOFI capture the temporal dynamics of corticogenesis (Figure 6A). Using experimental snapshots collected at gestational weeks 18, 19, 23, 24 and 52, the framework generates continuous trajectories that reveal the gradual transition from progenitor-dominated to neuron-dominated states.

**Figure 6:**
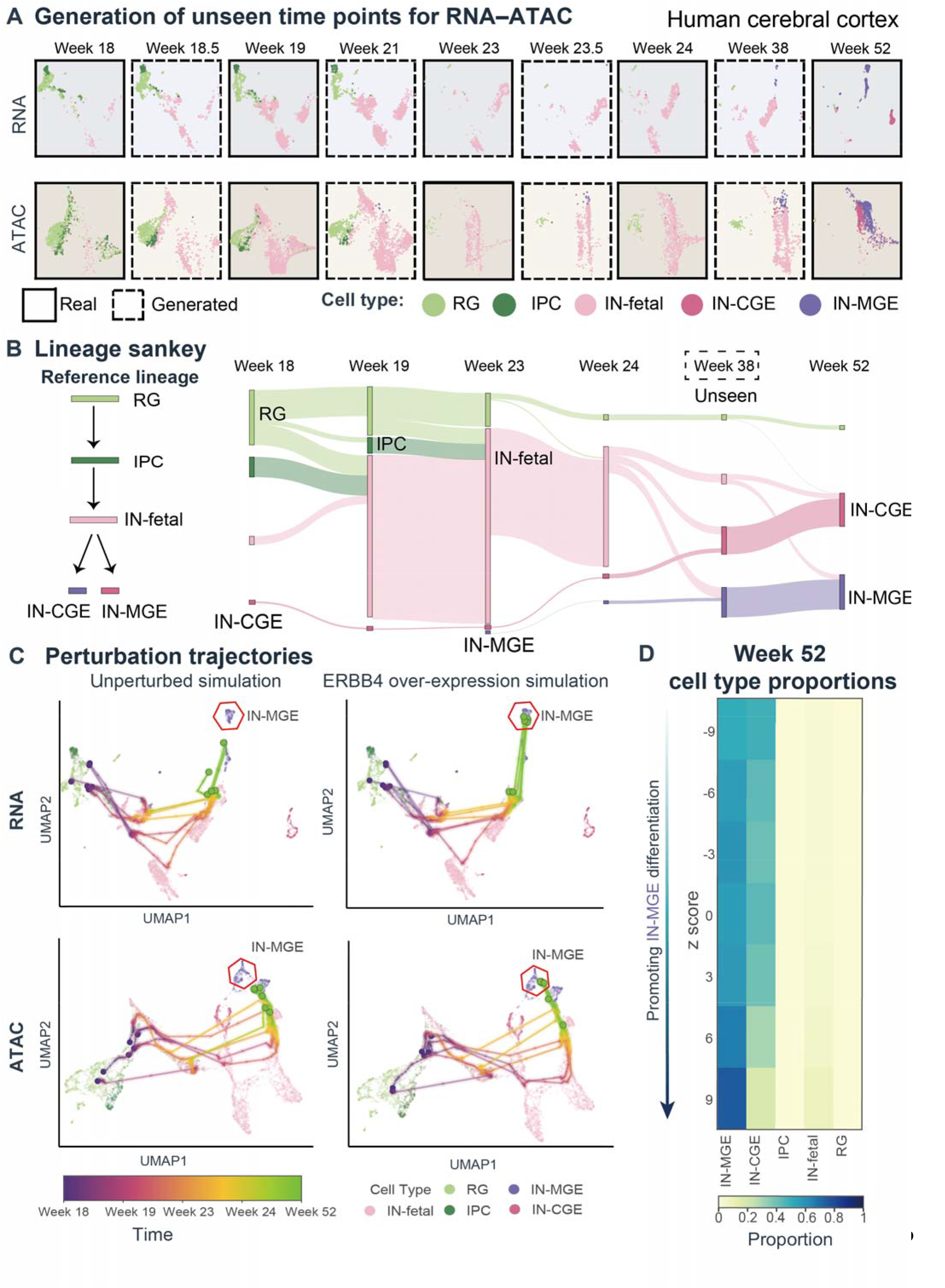
In silico lineage reconstruction and perturbation modeling in the human cerebral cortex. (A) Generation of unseen time points. MOFI generates continuous snapshots for Weeks 18.5, 21, 23.5, and 38 (dashed boxes) by interpolating between experimental RNA– ATAC snapshots at Weeks 18, 19, 23, 24, and 52 (solid boxes). (B) Multi-stage lineage Sankey diagram. Left: The reference lineage hierarchy. Right: Sankey visualization illustrating the dynamic shift in cell-type proportions and their predicted ancestral origins across developmental stages. The flow width represents the transport intensity between consecutive time points. (C) Perturbation-induced fate shifts. Comparison of simulated trajectories between the unperturbed scenario (left) and *ERBB4* overexpression (right) across RNA and ATAC omics. (D) Lineage composition under perturbation intensity. Heatmap showing cell-type proportions at Week 52 under a gradient of *ERBB4* perturbation strengths.

A core question in cortical development is how progenitors commit to distinct terminal neuronal fates. To address this, we constructed a multi-stage lineage Sankey diagram demonstrating experimentally validated hierarchies[26] (left panel) with MOFI-reconstructed proportional cell flow dynamics (right panel, Figure 6B)[26]. The overall trend aligns with the canonical hierarchy: RG dominates at week 18, generating IPCs by week 19; IPCs differentiate into IN-fetal cells, which emerge as the dominant cell population by week 23. IN-fetal subsequently branches into IN-CGE and IN-MGE by week 52, with RG and IPC populations nearly depleted. Overall, the agreement between experimental lineages and MOFI-predicted flows supports the ability of the framework to reconstruct the reference lineage progression in human brain development.

To test whether the learned dynamics capture the sensitivity of cortical lineage allocation to molecular perturbation, we performed in silico overexpression of *ERBB4*, a receptor tyrosine kinase implicated in cortical interneuron migration and maturation[34][35] (Figure 6C). The perturbation analysis reveals a pattern: *ERBB4* overexpression predominantly affects late-stage cell-type composition. In the RNA space, the perturbed trajectories show increased representation of IN-MGE populations at the expense of other fates, while the ATAC space exhibits corresponding changes in chromatin accessibility. The perturbation intensity sweep provides a quantitative assessment of this dose-response relationship (Figure 6D). Across a signed range of *ERBB4* perturbation strengths, interneuron-associated fates respond monotonically: positive perturbation shifts the predicted allocation toward IN-MGE-associated states while decreasing other populations, and negative perturbation produces the opposite effect. These results identify a model-predicted association between *ERBB4* perturbation and interneuron fate allocation that is consistent with existing biological knowledge[34][35].

## Discussion

By jointly modeling paired omics in a dynamic manner, MOFI improves the reconstruction of continuous cell-state dynamics when the two omics provide complementary information. For instance, in the controlled simulation, omics-1 captured only part of the information governing the underlying transitions, whereas omics-2 contained additional state information that influenced the trajectory. Combining the two omics improved the recovery of held-out intermediate states (Figure 2F) and the underlying trajectory structure, and a similar ability was supported by the hold-one-out comparison across real-world datasets (Supplementary Note 2). These results suggest that paired omics can reduce ambiguity in trajectory reconstruction by requiring inferred dynamics to remain consistent with both molecular layers.

A practical advantage of MOFI in experimental design arises when the two omics are sampled at different temporal resolutions, that is, one omics is sampled less densely in time than the other. Our simulation used five sampling times for omics-1 and three for omics-2, and MOFI recovered intermediate states of the more sparsely observed layer by combining its measurements with information from the more densely sampled omics. Together with the real-data single omics hold-out analyses, these results suggest that asymmetric temporal sampling is feasible when one molecular layer is substantially more difficult or costly to profile densely.

The ability to retain complementary information across omics also suggests a characteristic between MOFI and other approaches that first merge molecular measurements into a shared representation. Direct concatenation places paired features in a common state space before dynamical inference, whereas scMultiNODE[20] learns dynamics in a shared latent representation. MOFI instead retains omics-specific representations while explicitly modeling how the two omics change over developmental time. The kidney organoid analysis illustrates the practical consequence of this design: RNA expression and chromatin accessibility followed the same overall developmental progression while retaining distinct molecular patterns, and their joint modeling recovered intermediate cell states and temporal marker dynamics (Figure 3A). These results suggest that synchronized modeling may be particularly useful when different omics provide related but non-redundant views of the same developmental process.

Biologically, by preserving this cross-omics correspondence throughout dynamical inference, MOFI enables downstream analyses that extend beyond trajectory reconstruction. In the HSPC RNA–protein data, the learned relationship associated transcriptomic changes with model-predicted protein changes. The *ITGA2B*–CD41 analysis provided an example of coordinated RNA–protein responses (Figure 4F). The in silico perturbation analyses further quantified how the fitted dynamical model responded to changes in selected molecular features, including predicted shifts in HSC-derived fate allocation and ERBB4-associated changes in cortical interneuron composition (Figure 6C, D). By embedding the cross-omics relationship within the dynamical model, MOFI allows coordinated omics changes to be examined along cell-fate progression.

From an algorithmic perspective, building on the theoretical framework of synchronized UOT[22], MOFI introduces a deep learning based dynamical modeling methodology to improve the computational scalability of the formulation, enabling processing of large single-cell multi-omics datasets and downstream biological analyses.

Several limitations remain. First, the inferred population-change field represents relative expansion or depletion of modeled cell states. Changes in observed population mass can also reflect sampling and other sources of compositional variation. Second, continuous trajectories reconstructed from destructive snapshots are not uniquely determined by the observations. MOFI uses temporal distribution matching and paired multi-omics information to reduce this ambiguity, but the inferred dynamics remain dependent on the assumptions of the model. Third, the in silico perturbation responses and cross-omics transformation programs should be interpreted as model-based hypotheses for cross-omics responses and fate redistribution and therefore require experimental validation. Finally, the current implementation is limited to paired two-omics datasets and relies on numerical integration of neural dynamical models, which can become computationally demanding for very large datasets. Extending the framework to additional molecular or spatial layers and incorporating more efficient flow-based or simulation-free strategies[16][17][36] may broaden its applicability.

Overall, MOFI provides a dynamically constrained framework for integrating paired time-series multi-omics measurements and converting discrete snapshots into interpretable models of cell-fate progression. By linking coordinated molecular changes across omics with population dynamics, perturbation responses, and candidate cross-omics transformation programs, the framework provides a basis for formulating experimentally testable hypotheses about how molecular coordination shapes developmental fate decisions.

## Methods

### Data and problem formulation

We consider time-series multi-omics data measured at ordered experimental times 0=*t*_0_ < *t*_1_ <… <*t*_*M*_ = 1. At each time point *t*_*m*_, we observe a collection of cells indexed by *i*= 1,…*N*_*m*_, each with an omics-1 coordinate 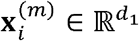 (e.g. the high-dimensional transcriptomic state after PCA), and another paired omics state 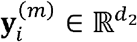 (e.g. protein abundance or chromatin accessibility after PCA). This formulation can be extended to multi-omics observations that are paired at only a subset of time points. In practical applications, MOFI can combine the lower-resolution measurements obtained under the more costly omics assays with the higher-resolution measurements obtained from RNA (Supplementary Notes 1,2). The two state spaces are

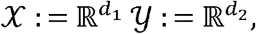

with generic elements denoted as **x**∈*X*, **y** ∈*Y*. The two empirical distributions of cells at time *t*_*m*_ are

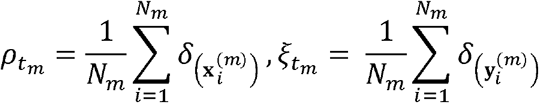

which we regard as a finite-sample approximation of an underlying population distribution *ρ*(·, *t*) on *ξ*(·, *t*) *X,Y*. Our goal is to infer the underlying single-cell dynamics that interpolate the observed snapshots and jointly capture multi-omics velocity, cell proliferation, and cell death across both state spaces, while maintaining consistency between the dynamical behavior across omics.

### Overview of optimal transport

A natural way to couple the time-indexed distributions 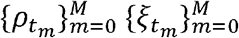 is to view them as marginals of a continuous flow of measures {*ρ*(·*t*)}_*t*∈[0,1]_,{ξ(·,*t*)}_*t*∈[0,1]_driven by a dynamical system on *X and Y*. Optimal transport (OT) theory provides complementary perspectives on such flows[11].

#### Dynamical optimal transport

In single-omics dynamical OT [37], the evolution of *ρ* is described by a velocity field 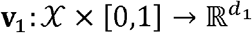 through the continuity equation

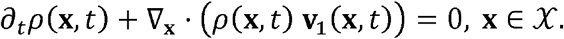

Among all pairs (*ρ*, **v**_1_) that satisfy this equation and match the observed endpoint distributions, dynamical OT selects those with minimal kinetic energy, measured by the action functional

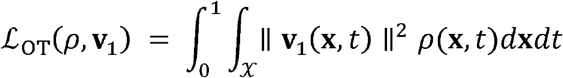

This action quantifies the minimal effort required to morph one population distribution into another under a mass-preserving flow, and the optimal trajectories provide a continuous interpolation of cell states in the omics-1 expression space.

#### Unbalanced transport and growth

However, in time-series single-cell data, the total number of cells typically changes across time due to proliferation, apoptosis, and sampling bias. To allow for such mass variations, unbalanced dynamical OT[38][39] introduces a scalar growth rate *g* :*X*×[0,1] → ℝ and relaxes the continuity equation to

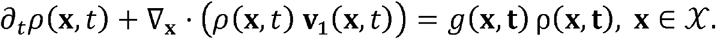

The energy is augmented by a penalty on deviations from mass conservation, yielding

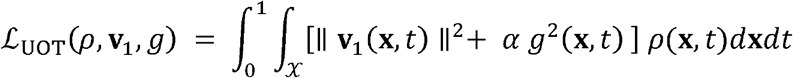

where *α* > 0 controls the strength of the growth regularization. In this formulation, **v**_1_ encodes where cells move through the omics-1 expression space, while *g*(·,*t*)captures where cells tend to be amplified or depleted over time.

### Synchronized Unbalanced Optimal Transport for multi-omics dynamics

When multi-omics characterize the same dynamically evolving biological system, it is insufficient to model biological dynamics solely within a single-omics space. The synchronized unbalanced optimal transport formulation introduces a differentiable cross-omics map and enforces growth consistency for matched cells across omics.

To address this requirement, the synchronized unbalanced dynamical OT (Sync-UOT) framework introduces a cross-omics map *T*^1→2^:*X→Y* together with its inverse counterpart *T*^1→2^:*Y→X*. We further define the omics-2 velocity field 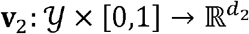, the omics-2 growth rate function 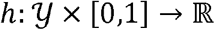, and impose a synchronized growth consistency constraint: *h*(*T* ^1→2^(**x**),*t*) = *g*(**x**,*t*). This constraint reflects a feasible biological assumption that paired measurements of the same cells should exhibit the same growth rate across the two omics.

The Sync-UOT problem extends the single-space UOT formulation to a multi-space system (*X* ×*Y*). The full continuous optimization problem is:

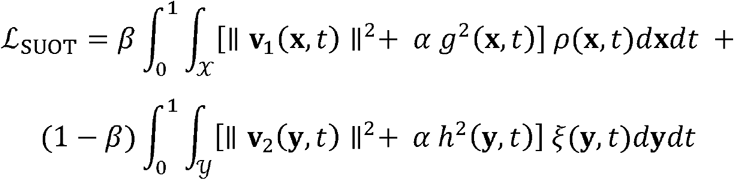

subject to the multi-omics continuity equation

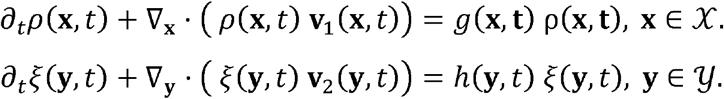

Here, 0 < *β* < 1 balances the contribution of the two omics. In this setting, v_2_ encodes where cells move in omics-2 space.

A central theoretical result is that this multi-omics problem admits an equivalent reduced formulation in a single space[22]. Specifically, by substituting the omics-2 space constraints and applying the pushforward relation 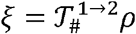 from *T*^1→ 2^(**x**) =***y*** together with the above growth consistency condition, the problem reduces to

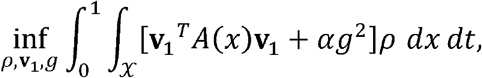

where the effective metric tensor is 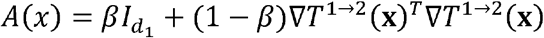 (Supplementary Note 3). This result has a clear geometric interpretation: synchronizing the two omics is equivalent to performing unbalanced optimal transport in the omics-1 under a modified Riemannian metric that incorporates the omics-2 geometry. This term penalizes velocity components more heavily in directions to which omics-2 is sensitive, thereby enforcing that inferred cellular trajectories remain biologically consistent across omics simultaneously.

### Weighted particle approximation

To numerically approximate *ρ* and *ξ* on the state spaces *X* and *Y*, respectively, we represent them by a finite system of *N* paired weighted particles 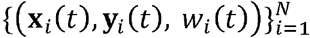, where **x**_*i*_(*t*) in *X* and **y**_*i*_(*t*) in *Y* denote the states of the *i*-th particle in the two omics spaces, and *w*_*i*_(*t*) denotes its associated population mass. At each time t, the corresponding empirical measures are defined as

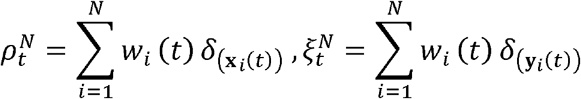

which provide discrete approximations of *ρ*_*t*_ and *ξ*_*t*_, respectively. The use of shared particle weights ensures that the paired states in the two omics spaces describe the same evolving cell population.

The particle locations and weights evolve according to

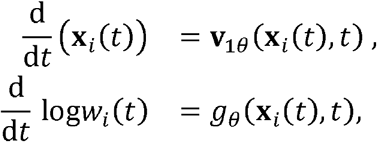

where **v**_1*θ*_ is the velocity field in the first omics space and *g*_*θ*_ is the growth rate, both parameterized by neural networks. States in the omics-2 space are coupled to those in the omics-1 space via the learned cross-omics map operator 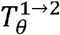, such that 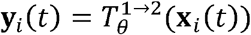. This particle-based formulation offers a finite-dimensional approximation for the underlying growth-augmented continuity equations, bridging the continuum dynamical model and its numerical realization. It permits the reconstruction of continuous, population-aware single-cell multi-omics trajectories from snapshot measurements collected at discrete time points.

### Numerical integration of particle flows across multi-omics

For each mini batch of initial cells sampled from the earliest time point *t*_0_, we initialize particle states by setting (**x**(*t*_0_)) equal to the observed cell states and set *w*_*i*_(*t*_0_)1/*N*. We then integrate the interacting ODE system forward in time from *t*_0_ to *t*_*M*_ using a standard explicit ODE solver with a uniform time grid. At each solver step, we: (i) compute the velocities **v**_1*θ*_, **v**_2*θ*_ and growth rate *g*_*θ*_ for all particles; (ii) update particle positions in omics-1 and weights according to ODEs; and (iii) derive particle positions in omics-2 by mapping the updated omics-1 positions via the learned cross-omics map 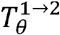, which is parameterized by neural networks.

### Training objective and optimization

MOFI is trained by optimizing the cross-omics map, the velocity and growth fields, and their agreement with the observed time-series snapshots. Let *v*_1*θ*_ denote the velocity field in the omics-1 space, *g*_*θ*_(*x, t*)the shared growth-rate field, and *T*_*θ*_ the learned cross-omics map. The overall objective is

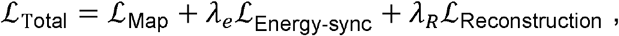

Each term serves a distinct and complementary purpose: *ℒ*_map_ is a bidirectional cross-omics mapping loss; *ℒ*_Energy-sync_ regularizes the velocity and growth fields according to the metric A(x); and the reconstruction loss *ℒ*_Reconstruction_ aligns simulated particle distributions with experimental snapshots across all time points while incorporating population mass matching through the mass loss term *ℒ*_Mass_. All *λ* and *α* coefficients are non-negative weighting parameters. Full details regarding the map training procedure, default coefficient values, and corresponding sensitivity analyses can be found in Supplementary Notes 4 and 5.

#### Cross-omics mapping loss

As illustrated in Figure 1C, the map consists of two omics-specific encoders, *E*_1*θ*_ and *E*_2*θ*_, and two omics-specific decoders, *D*_1*θ*_ and *D*_2*θ*_. Their compositions define four mappings,

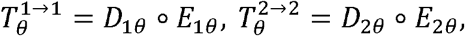

for within-omics reconstruction, and

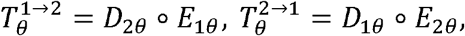

For each paired observation (**x**, **y**), the two encoders map the corresponding measurements into a shared latent space. We therefore additionally impose a latent consistency loss between *E*_2_(**x**) and *E*_2_(**y**), encouraging paired measurements to have concordant latent representations.

The overall mapping objective is

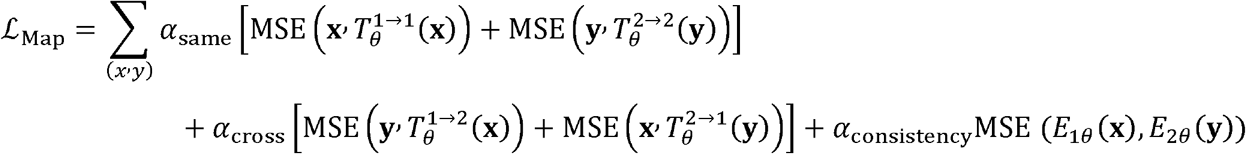

#### Synchronized energy

The synchronized energy regularization term is formulated as:

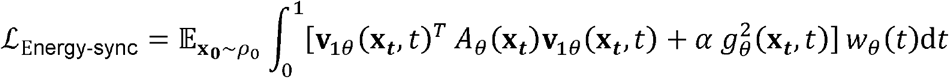

Where ***x***_*t*_ denotes the omics-1 state, and *wθ(t)* is the time-evolving particle weight. This expression is approximated using the weighted-particle system introduced above. Let *0=τ*_*0*_ < …_*1*_ <*τ* _*K*_ =1 denote the numerical integration grid. *W*_*i*_*(τ*_*k*_)is the population mass carried by particle *I* at time *τ*_*k*_, then

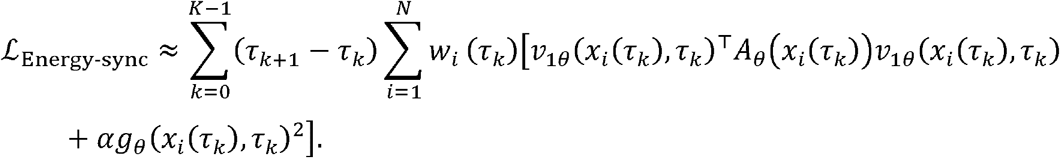

Thus, the same particle states and weights that evolve according to *v*_*1θ*_*(x,t)*,_and *g*_*θ*_also define the discrete integration for energy. Minimizing this term ensures plausible dynamical trajectories, prevents overfitting to discrete snapshot data, and enforces that omics-1 dynamics respect the dynamical structure of the omics-2 layer.

#### Snapshot reconstruction and population-mass matching

*ℒ*_Reconstruction_ aligns the simulated particle system with observed data through a combination of optimal transport loss *ℒ*_OT_ and a mass matching loss *ℒ*_Mass_:

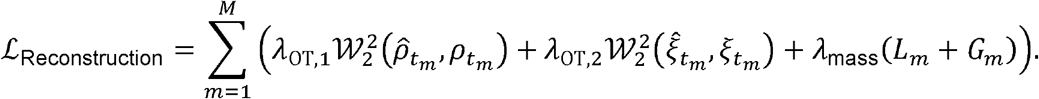

where 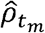 and 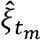 denote the reconstructed weighted particle distributions at time *t*_*m*_. They are obtained by numerically integrating the particle states and weights in the omics-1 space and mapping the resulting states to the omics-2 space through the learned cross-omics map 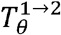. The coefficients *λ*_OT,1_ and *λ*_OT,z_ control the contributions of distribution matching in the two omics spaces, respectively.

For two normalized empirical distributions 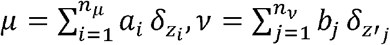, we use the discrete quadratic Wasserstein distance to measure the distance between the two distributions:

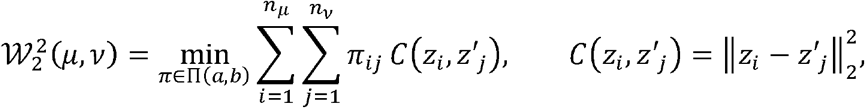

where *Π* (*a,b*) = {*π*≥ *0: π***1** *= a, π*^⊤^ *⊤* **1***= b*} is the set of transport couplings with marginals *a* and *b*.

In addition to distributional alignment, *ℒ*_Mass_ is formulated using two complementary terms to infer cell proliferation and death rates. The local mass error m ensures that the weight distribution of simulated particles aligns with the local density of the real data:

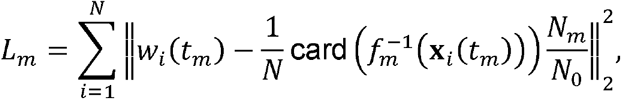

where *f*_*m*_ maps real data points to their closest simulated particles, thus card 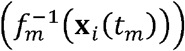 providing the number of observed cells assigned to particle *i*. Here *N*_*m*_and *N*_*0*_ denote he numbers of observed paired cells at times *t*_*m*_ and *t*_*0*_, respectively, so that *N*_*m*_*/N*_*0*_ provides the observed relative population-size ratio used to supervise the particle mass. Simultaneously, to ensure the model captures macroscopic population dynamics, we enforce the global consistency term 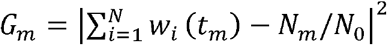, which constrains the total accumulated weight to match the observed relative expansion or contraction of the cell population size.

#### Optimization strategy

To improve convergence, we employ a three-stage training strategy. First, the map operator *T*_*θ*_ is pretrained using the four-direction reconstruction objective, providing a reliable differentiable coupling between the omics-1 and omics-2 spaces. In the second stage, the velocity network *v*_*1θ*_ and growth network *g*_*θ*_ are initialized in the omics-1 space alone, minimizing a simplified objective dominated by the unbalanced OT reconstruction loss (*set β= 1* in *A(x)* and *λ* = 0 in *ℒ*_Reconstruction_). In the final stage, the full Sync-UOT objective is activated. This phased strategy prevents the optimization from diverging due to poor initial estimates of the transport map or the velocity field. A detailed algorithm is provided in Supplementary Note 5.

### Model architecture and implementation

All experiments were performed using a single NVIDIA GeForce RTX 4090 GPU with CUDA acceleration. We parameterized the omics-1 velocity field *ν*_1θ_, the growth *g*_θ_ rate, and the map operator *T*_θ_ using neural networks in PyTorch. The map comprises four functional subnetworks: 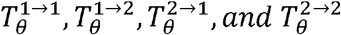. Following the Sync-UOT framework, the omics-2 velocity field is defined as 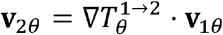. To capture the distinct dynamics of omics transitions, we employed a multilayer perceptron (MLP) for the velocity network. This architecture accepts concatenated state and time vectors *(x,t)* to compute high-dimensional velocities. Specifically, the network comprises two hidden layers with 400 units each and LeakyReLU activation functions. The growth rate *g*_*θ*_ utilizes a similar architecture. The map operator *T*_*θ*_ comprises two sub-encoders that map 50-dimensional principal components of the expression matrices to a shared latent space *z*. Two symmetric sub-decoders subsequently project these latent representations back to their respective 50-dimensional omics spaces.

### Data preprocessing

All real datasets were processed as paired or cross-modally matched time-series multi-omics objects. For datasets generated using same-cell multi-omics assays, the two molecular measurements corresponded to the same cells. For the human brain organoid dataset, we used the processed RNA–ATAC multi-omic metacell data[25] constructed from the developmental scRNA-seq (E-MTAB-12001) and scATAC-seq (E-MTAB-11998) datasets and retained the established cross-modal correspondence. Omics-specific feature matrices, sampling times, and cell identifiers were obtained from the processed data released by the original studies. Quality-control filtering was performed using the original annotations and dataset-specific marker information. Cells were retained only when they were present in both omics-specific objects, after which cell identifiers and ordering were synchronized to preserve the pairing between the two omics measurements.

For the transcriptomic layer, raw count matrices, when available, were normalized by library size to 10,000 counts per cell and log transformed. Highly variable genes were selected from the normalized expression matrix before dimensionality reduction. For the chromatin omics, ATAC measurements were represented as gene-activity matrices. When the input consisted of peak counts, peaks overlapping a gene body were assigned to that gene; peaks outside gene bodies were assigned to the nearest gene within 100 kb, and peak counts assigned to the same gene were summed. Gene-activity matrices were subsequently processed independently from the RNA matrices. For RNA–protein data, the antibody-derived protein abundance matrix was used as the omics-2 representation after the quality-control procedures applied to the paired object.

Each omics layer was processed independently to retain its omics-specific variation. Where the number of measured features permitted, 2,000 highly variable features were selected separately for each omics. Principal-component analysis was then applied to each omics, and the first 50 principal components were used as the input representation for MOFI. The fitted PCA means and loading matrices were retained so that inferred latent trajectories could subsequently be projected back to the original molecular-feature space.

Experimental sampling times were mapped to monotonically increasing numerical coordinates used by the dynamical model, while the original biological times were retained in the cell metadata for interpretation and visualization. The resulting analysis objects therefore contained synchronized cell identities, paired 50-dimensional omics representations, experimental time labels and the original cell-state annotations. These processing steps were applied consistently across the kidney organoid RNA–ATAC, hematopoietic RNA–protein, brain organoid RNA– ATAC and developing cerebral cortex RNA–ATAC datasets.

### Downstream analyses

#### Continuous trajectory reconstruction

Continuous multi-omics trajectories were generated from the trained MOFI model by initializing weighted particles from cells collected at the earliest experimental time point. Initial cells were sampled without replacement using a fixed random seed of 42. Particle states were propagated through the learned velocity and growth fields by numerical integration with an ODE step size of 0.05. Particle positions, velocities, weights, and inferred growth rates were recorded at the experimental times observed and at intermediate evaluation times used for continuous trajectory reconstruction. Trajectories were first integrated in the omics-1 space and were subsequently mapped to the omics-2 space through the learned cross-omics map. The reconstructed states in both omics were stored together with their inferred time, trajectory identifier, source-cell identifier, and particle weight. Additional numerical experiments, hold-one-out analyses comparing diverse interpolation strategies, and supplementary results from all biological datasets are provided in Supplementary Notes 2 and 6–9.

#### Cell-state assignment and population dynamics

Cell type labels for experimentally unobserved states were assigned using a class-balanced multinomial logistic-regression classifier. The classifier was trained on the concatenated 50-dimensional representations of the paired omics from experimentally measured cells, using the L-BFGS optimizer and a maximum of 1,200 iterations. The original cell-type annotations provided with each dataset were used as training labels.

For every reconstructed particle, the classifier returned a probability distribution over the annotated cell states. Continuous cell-type composition was calculated by weighting these probabilities by the corresponding particle masses and normalizing across the simulated population at each time point. This procedure accounted for both state transitions and changes in the relative population mass inferred by the growth field.

For lineage-flow visualization, particle-weighted state transitions were aggregated between consecutive evaluation times. The resulting transition masses were used to construct Sankey diagrams describing the inferred redistribution of cells across developmental states. For the cortical lineage analysis, edges accounting for less than 0.08 of the corresponding source-state population were omitted from the visualization. This threshold was applied only to improve graphical readability and did not affect trajectory integration or cell-state prediction.

#### Reconstruction of molecular dynamics

RNA expression, ATAC gene activity, and protein-abundance trajectories were recovered from the inferred low-dimensional particle states using the stored PCA means and loading matrices. For a molecular feature or predefined feature program, its value at each evaluation time was summarized as a particle-weighted mean. Experimental reference values were calculated by aggregating the corresponding feature measurements from observed cells at each sampling time.

Temporal marker and program trajectories were compared between reconstructed and experimentally observed populations at matched time points. When standardized profiles were reported, values were z-score normalized within each molecular feature or program across the analyzed time course. This procedure was used for both individual markers and multi-feature developmental programs.

#### Visualization of inferred trajectories and growth

For visualization, experimentally measured cells were embedded in two dimensions using UMAP fitted separately to the processed representation of each omics. Reconstructed states were projected into the corresponding embedding using the same preprocessing and transformation procedure. Continuous particle paths were then overlaid on the experimentally observed cell-state manifold. The inferred growth rate *g* (**x**.*t*) was evaluated along reconstructed trajectories and visualized on the corresponding low-dimensional embedding. Positive and negative values of *g* indicate relative expansion and depletion of the modeled population mass, respectively.

#### In silico perturbation analysis

In silico perturbations were introduced by modifying selected molecular features in the original feature space. For each perturbed gene or gene set, signed perturbation values were defined as multiples of the empirical standard deviation of the corresponding feature. The modified molecular profiles were projected into the PCA space using the fitted preprocessing transformation and were then propagated through the same trained MOFI dynamics.

Both positive and negative perturbation strengths were evaluated to characterize the predicted direction and magnitude of the response. Perturbed and unperturbed particles were initialized from the same source population, enabling direct comparison of changes in trajectories and terminal cell-state composition. Cell-fate redistribution was quantified from the particle-weighted classifier probabilities of the full simulated population at the terminal time.

#### Cross-omics transformation programs

Cross-omics transformation profiles were derived from local sensitivities of the learned dynamical and cross-omics mapping functions. At each analyzed cell state, Jacobians of the velocity field and the forward or reverse cross-omics map were calculated in the latent representations. These transformation profiles were projected through the PCA reconstruction matrices to express the predicted responses in the original gene and protein feature spaces.

For the gene-to-protein analysis, each gene was represented by a vector describing its predicted association with changes across the measured protein features. Conversely, each protein was represented by its predicted association with changes across the transcriptomic features in the protein-to-gene analysis. Pairwise similarity between these vectors was used to identify molecular features with similar cross-omics transformation patterns. The resulting gene-to-protein and protein-to-gene profiles were embedded into two dimensions and grouped into nearest-neighbor-based classes for visualization.

#### Differential and functional analyses

Differential-feature analysis was performed between reconstructed populations assigned to the compared developmental stages. For each molecular feature, a two-sided Wilcoxon rank-sum test was used, and the resulting P values were adjusted for multiple testing with the Benjamini– Hochberg procedure.

## Data availability

All the datasets used in this paper are publicly available. The kidney organoid RNA–ATAC data were obtained from GEO under accession GSE213152. The hematopoietic stem and progenitor cell RNA–protein time-series data were obtained from GEO under accession GSE305370. Specifically, we used the CITE-seq data from donor 31800 at Days 2, 4, 7, and 10. The human brain organoid developmental time-course scRNA-seq and scATAC-seq data were obtained from ArrayExpress under accessions E-MTAB-12001 and E-MTAB-11998. The developing human cerebral cortex multi-omics data were obtained from GEO under accession GSE204684.

## Code availability

The simulated benchmark dataset was generated using the procedures described in Supplementary Note 1. The MOFI source code and scripts for preprocessing, model training and reproduction of the main analyses are released at https://github.com/jiantaoShen12/MOFI. The configuration files and the notebooks used to reproduce the results are made available through the same repository.

## Acknowledgements

This work was supported by the National Natural Science Foundation of China (NSFC Nos. 12288101, 8206100646 and T2321001 to P.Z.), the Fundamental and Interdisciplinary Disciplines Breakthrough Plan of the Ministry of Education of China (grant no. JYB2025XDXM502), and the Guangdong Major Project of Basic Research (2026B0303000003 to R.L.). We also acknowledge support from the High-performance Computing Platform of Peking University for computational support.

## Author information

Authors and Affiliations

**Center for Machine Learning Research, Peking University, Beijing, China**.

Jiantao Shen & Peijie Zhou

**AI for Science Institute, Beijing, China**.

Jiantao Shen & Peijie Zhou

**School of Mathematics, South China University of Technology, Guangzhou, China**.

Jiantao Shen, Hao Peng & Rui Liu

**LMAM and School of Mathematical Sciences, Peking University, Beijing, China**.

Zhenyi Zhang & Qiangwei Peng

**School of Future Technology, South China University of Technology, Guangzhou, China**.

Hao Peng

**Center for Quantitative Biology, Peking University, Beijing, China**

Zihan Wang, Peijie Zhou

**National Engineering Laboratory for Big Data Analysis and Applications, Beijing, China**

Peijie Zhou

## Contributions

P.Z. and R.L. conceived the study. J.S., Z.Z., H.P., Q.P., and Z.W. designed the algorithm. J.S. implemented the algorithm and performed data analysis. J.S., H.P. and P.Z. interpreted the results and wrote the paper. J.S. wrote the supplementary materials. P.Z. and R.L. supervised the research.

## References

[1] Stuart, T. & Satija, R. Integrative single-cell analysis. Nat. Rev. Genet. 20, 257–272 (2019).

[2] Argelaguet, R., Cuomo, A. S. E., Stegle, O. & Marioni, J. C. Computational principles and challenges in single-cell data integration. Nat. Biotechnol. 39, 1202–1215 (2021).

[3] Buenrostro, J. D., Giresi, P. G., Zaba, L. C., Chang, H. Y. & Greenleaf, W. J. Transposition of native chromatin for fast and sensitive epigenomic profiling of open chromatin, DNA-binding proteins and nucleosome position. Nat. Methods 10, 1213–1218 (2013).

[4] Cao, J. et al. Joint profiling of chromatin accessibility and gene expression in thousands of single cells. Science 361, 1380–1385 (2018).

[5] Stoeckius, M. et al. Simultaneous epitope and transcriptome measurement in single cells. Nat. Methods 14, 865–868 (2017).

[6] Schiebinger, G. et al. Optimal-Transport Analysis of Single-Cell Gene Expression Identifies Developmental Trajectories in Reprogramming. Cell 176, 928–943 (2019).

[7] Baião, A. R. et al. A technical review of multi-omics data integration methods: from classical statistical to deep generative approaches. Brief. Bioinform. 26, bbaf355 (2025).

[8] Vandereyken, K., Sifrim, A., Thienpont, B. & Voet, T. Methods and applications for single-cell and spatial multi-omics. Nat. Rev. Genet. 24, 494–515 (2023).

[9] Argelaguet, R. et al. MOFA+: a statistical framework for comprehensive integration of multi-modal single-cell data. Genome Biol. 21, 111 (2020).

[10] Hao, Y. et al. Integrated analysis of multimodal single-cell data. Cell 184, 3573–3587 (2021).

[11] Zhang, Z. et al. Deciphering cell-fate trajectories using spatiotemporal single-cell transcriptomic data. NPJ Syst. Biol. Appl. 12, 2 (2026).

[12] Trapnell, C. et al. The dynamics and regulators of cell fate decisions are revealed by pseudotemporal ordering of single cells. Nat. Biotechnol. 32, 381–386 (2014).

[13] La Manno, G. et al. RNA velocity of single cells. Nature 560, 494–498 (2018).

[14] Tong, A., Huang, J., Wolf, G., Van Dijk, D. & Krishnaswamy, S. TrajectoryNet: a dynamic optimal transport network for modeling cellular dynamics. Proc. Mach. Learn. Res. 119, 9526–9536 (2020).

[15] Zhang, Z., Li, T. & Zhou, P. Learning stochastic dynamics from snapshots through regularized unbalanced optimal transport. In Proc. 13th International Conference on Learning Representations (ICLR, 2025).

[16] Wang, D. et al. Joint Velocity-Growth Flow Matching for Single-Cell Dynamics Modeling. Adv. Neural Inf. Process. Syst. 38, 178180–178216 (2025).

[17] Peng, Q. et al. WFR-FM: simulation-free dynamic unbalanced optimal transport. In Proc. 14th International Conference on Learning Representations (2026).

[18] Sha, Y., Qiu, Y., Zhou, P. & Nie, Q. Reconstructing growth and dynamic trajectories from single-cell transcriptomics data. Nat. Mach. Intell. 6, 25–39 (2024).

[19] Li, C., Virgilio, M. C., Collins, K. L. & Welch, J. D. Multi-omic single-cell velocity models epigenome–transcriptome interactions and improves cell fate prediction. Nat. Biotechnol. 41, 387–398 (2023).

[20] Zhang, J., Chakravarthy, M. & Singh, R. scMultiNODE: Integrative and Scalable Framework for Multi-Modal Temporal Single-Cell Data. In Proc. 16th ACM International Conference on Bioinformatics, Computational Biology and Health Informatics, Article 69 (ACM, 2025).

[21] Klein, D. et al. Mapping cells through time and space with moscot. Nature 638, 1065–1075 (2025).

[22] Cang, Z., Wang, J., Wei, X. & Zhao, Y. Synchronization of unbalanced dynamical optimal transport across multiple spaces. Preprint at arXiv:2602.18725 (2026).

[23] Yoshimura, Y. et al. A single-cell multiomic analysis of kidney organoid differentiation. Proc. Natl. Acad. Sci. USA 120, e2219699120 (2023).

[24] DeMeo, B. et al. Active learning framework leveraging transcriptomics identifies modulators of disease phenotypes. Science 390, eadi8577 (2025).

[25] Fleck, J. S. et al. Inferring and perturbing cell fate regulomes in human brain organoids. Nature 621, 365–372 (2023).

[26] Zhu, K. et al. Multi-omic profiling of the developing human cerebral cortex at the single-cell level. Sci. Adv. 9, eadg3754 (2023).

[27] Peng, Q., Zhou, P. & Li, T. stVCR: spatiotemporal dynamics of single cells. Nat. Methods 23, 542–553 (2026).

[28] Novershtern, N. et al. Densely interconnected transcriptional circuits control cell states in human hematopoiesis. Cell 144, 296–309 (2011).

[29] Cildir, G. et al. Understanding mast cell heterogeneity at single cell resolution. Trends Immunol. 42, 523–535 (2021).

[30] Gurish, M. F. et al. Intestinal mast cell progenitors require CD49dβ7 (α4β7 integrin) for tissue-specific homing. J. Exp. Med. 194, 1243–1252 (2001).

[31] Dumon, S. et al. Itga2b regulation at the onset of definitive hematopoiesis and commitment to differentiation. PLoS ONE 7, e43300 (2012).

[32] Mason, S., Piper, M., Gronostajski, R. M. & Richards, L. J. Nuclear factor one transcription factors in CNS development. Mol. Neurobiol. 39, 10–23 (2009).

[33] Eze, U. C. et al. Single-cell atlas of early human brain development highlights heterogeneity of human neuroepithelial cells and early radial glia. Nat. Neurosci. 24, 584–594 (2021).

[34] Flames, N. et al. Short- and long-range attraction of cortical GABAergic interneurons by neuregulin-1. Neuron 44, 251–261 (2004).

[35] Anton, E. S. et al. Receptor tyrosine kinase ErbB4 modulates neuroblast migration and placement in the adult forebrain. Nat. Neurosci. 7, 1319–1328 (2004).

[36] Tong, A. et al. Improving and generalizing flow-based generative models with minibatch optimal transport. Trans. Mach. Learn. Res. (2024).

[37] Benamou, J.-D. & Brenier, Y. A computational fluid mechanics solution to the Monge– Kantorovich mass transfer problem. Numer. Math. 84, 375–393 (2000).

[38] Chizat, L., Peyré, G., Schmitzer, B. & Vialard, F.-X. An interpolating distance between optimal transport and Fisher–Rao metrics. Found. Comput. Math. 18, 1–44 (2018).

[39] Chizat, L., Peyré, G., Schmitzer, B. & Vialard, F.-X. Unbalanced optimal transport: dynamic and Kantorovich formulations. J. Funct. Anal. 274, 3090–3123 (2018).

