## Supplementary Information for "MOFI reveals dynamic multi-omics coordination during cell fate progression"

**Contents**

**Supplementary Notes**

1. Simulation data generation and benchmark comparison

2. Benchmark comparison across real paired multi-omics datasets

3. Mathematical formulation and derivation

4. Transport map pretraining and convergence

5. Training pseudocode, implementation settings and hyperparameter sensitivity

6. HSPCs RNA–protein analyses

7. Human brain organoid RNA–ATAC analyses

8. Developing human cerebral cortex RNA–ATAC analyses

9. GSE213152 kidney nephron RNA–ATAC analyses

**Supplementary Figures**

Figs. 1–4. Transport map convergence and hyperparameter sensitivity

Figs. 5–9. HSPCs RNA–protein analyses

Figs. 10–13. Human brain organoid RNA–ATAC analyses

Figs. 14–17. Developing human cerebral cortex RNA–ATAC analyses

Figs. 18–19. GSE213152 kidney nephron RNA–ATAC trajectories

**Supplementary Tables**

Tables 1–4. Benchmark design and summary statistics

Tables 5–12. Pointwise W1 benchmarks on real datasets (HSPCs, brain organoid, cortex and kidney)

Table 13. Intermediate time point holdout reconstruction across real datasets

Tables 14–15. Recommended starting configuration and training settings for each dataset

Table 16. Predefined marker sets used for HSPCs perturbation analyses

Supplementary Note 1. Simulation data generation and benchmark comparison

Simulation data generation

To evaluate whether MOFI can recover continuous multi-omics dynamics when the underlying process and correspondence between omics are known, we constructed a synthetic paired longitudinal dataset from a stochastic dynamical system. Following the notation used in the main text, the regulatory state is represented by $X$, $Y$, and $I$, while $Z\left( X,Y \right)$ denotes a nonlinear landscape coordinate. The primary omics space consists of $\left( X,Y \right)$, whereas the paired secondary omics space consists of $\left( X,Y,Z \right)$. Thus, the secondary omics contain an additional nonlinear landscape feature while sharing the same underlying developmental process.

The dynamical variables $X$, $Y$, and $I$ were evolved using an Euler–Maruyama discretization with time step $dt=0.1$. The drift terms combine self activation, inhibition between variables and linear decay, with an additional localized repulsive force in the $X$–$Y$ plane:

$$dX=\left( \frac{C_{X}X^{2}+S}{1+C_{X}X^{2}+H_{Y}Y^{2}+H_{I}I^{2}+S}-d_{X}X+F_{X}\left( X,Y \right) \right)dt+\sigma_{X}dW_{X},$$

$$dY=\left( \frac{C_{Y}Y^{2}+S}{1+H_{X}X^{2}+C_{Y}Y^{2}+H_{I}I^{2}+S}-d_{Y}Y+F_{Y}\left( X,Y \right) \right)dt+\sigma_{Y}dW_{Y},$$

$$dI=\left( \frac{C_{I}I^{2}}{1+C_{I}I^{2}}-d_{I}I \right)dt+\sigma_{I}dW_{I}.$$

The parameters were $C_{X}=0.5$, $C_{Y}=1.5$, $C_{I}=1$, $H_{X}=0.3$, $H_{Y}=1$, $H_{I}=10$, $d_{X}=d_{Y}=d_{I}=0.4$, $S=1.5$, $\sigma_{X}=\sigma_{Y}=0.025$, and $\sigma_{I}=0.01$. A localized hill centered at $\left( X,Y \right)=\left( 2,1.5 \right)$ defines the additional coordinate in the secondary omics and the repulsive force:

$$Z\left( X,Y \right)=exp\left[ -\frac{\left( X-2 \right)^{2}}{2\times0.05}-\frac{\left( Y-1.5 \right)^{2}}{2\times0.05} \right],$$

$$F_{X}\left( X,Y \right)=-\nabla_{X}Z\left( X,Y \right)=\frac{Z\left( X,Y \right)\left( X-2 \right)}{0.05},$$

$$F_{Y}\left( X,Y \right)=-\nabla_{Y}Z\left( X,Y \right)=\frac{Z\left( X,Y \right)\left( Y-1.5 \right)}{0.05}.$$

The landscape coordinate $Z$ is a deterministic nonlinear function of $X$ and $Y$ and therefore has no independent stochastic evolution. This construction makes the secondary omics informative about the geometry that shapes the underlying regulatory dynamics.

The initial population contained 400 cells from two equally sized Gaussian components. Two hundred cells were sampled around $\left( X,Y,I \right)=\left( 2.8,0.2,0 \right)$ and 200 around $\left( 0,0,2 \right)$, with standard deviation 0.1 in each regulatory coordinate; negative coordinates were truncated to zero. Population change was introduced through a division probability that depends on state $P_{\mathrm{div}}=0.05\frac{Y^{2}}{1+Y^{2}}dt$. Therefore, the two branches exhibit distinct behaviors: one undergoes progressive branching (upper‑right), while the other shows no obvious branching events.

The simulator generated ground truth states for both omics at simulation times 0, 6, 12, 18 and 24 (normalized indices t = 0, 1, 2, 3 and 4). To reproduce the asymmetric temporal sampling design described in the main text, the primary omics was supplied at all five time points, whereas the secondary omics was supplied at only three time points (t = 0, 2 and 4). The simulator nevertheless retained ground truth secondary states at all five times for evaluation. The holdout experiments described below remove additional observations from this already asymmetric design and therefore use fewer training observations than the base setting.

Paired and single omics holdout evaluation on simulated data

Starting from the asymmetric observation design above, with five observed time points in the primary omics and three observed time points in the secondary omics, we evaluated reconstruction at intermediate target times $t=1,2$and $3$. Throughout this section, a holdout target denotes a target-time distribution that is unavailable to the model during fitting, either because the corresponding snapshot is explicitly masked or because it is absent from the original asymmetric sampling design.

In the paired target-time holdout setting, neither omics provides an observation at the target time during model fitting. For $t=1$ and $t=3$, the corresponding primary snapshot was explicitly withheld, whereas the secondary snapshot was already unobserved under the base sampling design. For $t=2$, the observed snapshots in both omics were explicitly withheld. Ground-truth states generated by the simulator at the target time were used only for evaluation.

We additionally evaluated modality-specific target holdouts while retaining all available observations from the other omics. The primary-omics holdout at $t=2$ removes only the primary snapshot at $t=2$, while retaining the secondary observations at $t=0,2$ and $4$. For secondary-omics evaluation, $t=1$and $t=3$are naturally unobserved target times under the asymmetric sampling design, whereas the $t=2$experiment additionally removes the observed interior secondary snapshot, leaving only $t=0$and $4$. These settings therefore assess reconstruction under progressively different patterns of temporal information availability.

All original results are saved in Supplementary Table 1. The first three rows of the table correspond to the three numerical results in the upper left of Fig. 2F, the fourth to sixth rows correspond to the three numerical results in the upper right, and the seventh to tenth rows correspond to the single omics holdout results shown in the bottom panel.

MOFI and direct concatenation baseline adapted from stVCR[1] were evaluated using five independent final training seeds (22–26), with MOFI validation seeds 27–29 used for model selection. MOSCOT[2] is deterministic and is reported as a single value. For paired snapshot holdout experiments, MOSCOT predictions were obtained by pairwise optimal transport displacement interpolation between the available flanking snapshots.

Supplementary Note 2. Benchmark comparison across real paired multi-omics datasets

Datasets and evaluation design

Using the models trained as described in the main text, we evaluated MOFI on four paired real-world datasets: HSPCs RNA–protein, human brain organoid RNA–ATAC, the developing human cerebral cortex branch (HC) RNA–ATAC, and GSE213152 kidney nephron RNA–ATAC. For each benchmark dataset and omics, generated cells at each time point after the initial state were compared with the corresponding observed cells using the Wasserstein 1 (W1) distance; lower values indicate closer distributional reconstruction.

We compared MOFI with direct concatenation and scMultiNODE[3]. Direct concatenation was implemented as a natural extension of the stVCR dynamical backbone: paired features from each omics were concatenated into a single joint input before dynamical inference, and principal component analysis (PCA) was applied to the concatenated input to obtain a 50D state representation for the real dataset analyses so that the dynamical model retained the same state dimension used elsewhere in the benchmark. No explicit map between omics was used in this baseline. scMultiNODE provided a neural temporal baseline for multi-omics data. Results are reported for each dataset and omics and summarized across the 40 evaluation points after the initial time.

Dataset characteristics are summarized in Supplementary Table 2. Time averaged W1 distances are reported in Supplementary Table 3, and Supplementary Table 4 summarizes pairwise comparisons across the 40 evaluation points after the initial time. Complete pointwise reconstruction results are provided in Supplementary Tables 5–12. The dedicated intermediate time point holdout results are reported in Supplementary Table 13.

Intermediate time point holdout reconstruction on real datasets

We extended the holdout experiments to all four real paired multi-omics datasets and to every intermediate sampled time point. The earliest and latest snapshots were retained in every experiment. At each remaining time, two masks were evaluated. In the paired holdout, both modalities at the target time were excluded from fitting and reconstruction was evaluated separately in the primary and secondary spaces. In the omics holdout, only the secondary observation was excluded while the paired primary observation remained available, and reconstruction was evaluated in the secondary space. The evaluated intermediate times were t = 1, 2 and 3 for GSE213152 and the developing human cerebral cortex dataset, t = 1 through 8 for the human brain organoid dataset, and t = 1 and 2 for HSPCs. This design produced 48 withheld distributions: 16 primary and 16 secondary evaluations under paired holdout, plus 16 secondary evaluations under secondary omics holdout. (Supplementary Table 13)

The dedicated holdout benchmark compared MOFI with scMultiNODE and direct concatenation baseline. Every model and scenario used seed 42. W1 was evaluated only at the omitted time point in the canonical stored PCA space for the corresponding dataset, with lower values indicating closer reconstruction of the withheld distribution. Because the omitted cells were not used for model fitting, these experiments test recovery of missing observations.

MOFI achieved the lowest W1 distance in 46 of the 48 withheld distributions (95.8%). It ranked first in all 32 secondary space evaluations, including every secondary reconstruction under paired holdout and every secondary omics holdout. For primary space reconstruction under paired holdout, MOFI ranked first in 14 of 16 cases. Across GSE213152 and HSPCs, MOFI yielded the lowest W1 at every omitted time and for every evaluated modality. In the human brain organoid dataset, MOFI also ranked first in all 24 evaluations across t = 1 through 8. In the developing human cerebral cortex dataset, MOFI ranked first in all six secondary space evaluations and in one of the three primary space evaluations. Together, these results show consistent reconstruction of withheld intermediate time points across the four paired multi-omics datasets.

Supplementary Note 3. Mathematical formulation and derivation

Synchronized unbalanced optimal transport formulation

$$C_{M}\left( \mu,\nu;\alpha,\beta\right):=\inf_{\left( \rho,\mathbf{v}_{1};\xi,\mathbf{v}_{2} \right)}\int_{0}^{1} \left( \beta\int_{X} \rho\left( \parallel\mathbf{v}_{1}\parallel^{2}+\alpha g^{2} \right)d\mathbf{x}+\left( 1-\beta\right)\int_{Y} \xi\left( \parallel\mathbf{v}_{2}\parallel^{2}+\alpha h^{2} \right)d\mathbf{y} \right)dt$$

subject to

$$\left\{ \begin{matrix} \partial_{t}\rho+\nabla\cdot\left( \rho\mathbf{v}_{1} \right)-g\rho=0, \\ \rho\left( \cdot,0 \right)=\mu, \\ \rho\left( \cdot,1 \right)=\nu, \\ \partial_{t}\xi+\nabla\cdot\left( \xi\mathbf{v}_{2} \right)-h\xi=0, \\ \xi\left( \cdot,t \right)=T_{♯}(\rho\left( \cdot,t \right)d\mathbf{x}), \\ g\left( \mathbf{x},t \right)=h\left( T\left( \mathbf{x} \right),t \right), \rho\left( \cdot, t \right)-a.e. \end{matrix} \right.$$

Equivalence to generalized dynamic optimal transport

Let $X\subset\mathbb{R}^{d_{1}}$and $Y\subset\mathbb{R}^{d_{2}}$, and let $T:X\to Y$be a time-independent $C^{1}$injective immersion. Let $\rho\left( \cdot, t \right)$denote the density of the primary-space population with respect to Lebesgue measure on $X$, and define the secondary-space population as the push-forward measure

$$\xi_{t}=T_{\#}(\rho\left( \cdot, t \right)\text{ }dx).$$

For the differentiable, injective map T used to couple the two spaces, which does not depend on time, the synchronized formulation reduces to the following generalized dynamic optimal transport problem[4]:

$$\min_{\left( \rho,\mathbf{v}_{1},g \right)}\int_{0}^{1} \int_{X} \rho\left\{ \mathbf{v}_{1}^{\top}\left[ \beta I+\left( 1-\beta\right)\left( \nabla T \right)^{\top}\left( \nabla T \right) \right]\mathbf{v}_{1}+\alpha g^{2} \right\}d\mathbf{x}dt$$

subject to

$$\partial_{t}\rho+\nabla\cdot\left( \rho\mathbf{v}_{1} \right)-g\rho=0, \rho\left( \cdot,0 \right)=\mu, \rho\left( \cdot,1 \right)=\nu.$$

*Proof.* Fix $t\in[0,1]$ and let $\xi( \cdot,t)=T_{♯}\rho( \cdot,t)$. For any test function $\phi\in C_{c}^{\infty}(Y)$ we have

$$\int_{Y} \left( \partial_{t}\xi-h\xi\right) \phi\left( y \right) dy=-\int_{Y} \nabla\cdot\left( \xi v_{2} \right) \phi dy=\int_{Y} \xi v_{2}\cdot\nabla\phi dy.$$

Substituting $\xi=T_{♯}\rho$ on the left side gives

$$\int_{Y} \left( \partial_{t}\xi-h\xi\right) \phi dy=\frac{d}{dt}\int_{Y} \phi\left( y \right)\xi\left( y,t \right) dy-\int_{Y} \phi\left( y \right)h\left( y,t \right)\xi\left( y,t \right) dy.$$

$$\frac{d}{dt}\int_{Y} \phi\left( y \right)\xi\left( y,t \right) dy-\int_{Y} \phi\left( y \right)h\left( y,t \right)\xi\left( y,t \right) dy=\frac{d}{dt}\int_{X} \phi\left( T\left( x \right) \right)\rho\left( x,t \right) dx-\int_{X} \phi\left( T\left( x \right) \right)g\left( x,t \right)\rho\left( x,t \right) dx.$$

$$\frac{d}{dt}\int_{X} \phi\left( T\left( x \right) \right)\rho\left( x,t \right) dx-\int_{X} \phi\left( T\left( x \right) \right)g\left( x,t \right)\rho\left( x,t \right) dx=\int_{X} \left( \partial_{t}\rho-g\rho\right)\left( \phi\circ T \right) dx.$$

Using the continuity equation $\partial_{t}\rho+\nabla\cdot(\rho\mathbf{v}_{\boldsymbol{1}})-g\rho=0,$ on the right side yields

$$\int_{X} \left( \partial_{t}\rho-g\rho\right)\left( \phi\circ T \right) dx=-\int_{X} \nabla\cdot\left( \rho v_{1} \right)\left( \phi\circ T \right) dx.$$

$$-\int_{X} \nabla\cdot\left( \rho v_{1} \right)\left( \phi\circ T \right) dx=\int_{X} \rho v_{1}\cdot\nabla\left( \phi\circ T \right) dx.$$

$$\int_{X} \rho v_{1}\cdot\nabla\left( \phi\circ T \right) dx=\int_{X} \rho\left[ \nabla\phi\left( T\left( x \right) \right) \right]^{\top}\left[ \nabla T\left( x \right) \right]v_{1} dx.$$

Pulling the secondary space velocity term back to X gives

$$\int_{Y} \xi v_{2}\cdot\nabla\phi dy=\int_{X} \rho v_{2}\left( T\left( x \right),t \right)\cdot\nabla\phi\left( T\left( x \right) \right) dx.$$

$$\int_{X} \rho v_{2}\left( T\left( x \right),t \right)\cdot\nabla\phi\left( T\left( x \right) \right) dx=\int_{X} \rho\left[ \nabla\phi\left( T\left( x \right) \right) \right]^{\top}v_{2}\left( T\left( x \right),t \right) dx.$$

So

$$\int_{X} \rho\left( \nabla\phi\left( T\left( x \right) \right) \right)^{\top}\left( \nabla T\left( x \right) \right)v_{1} dx=\int_{X} \rho\left( \nabla\phi\left( T\left( x \right) \right) \right)^{\top}v_{2}\left( T\left( x \right),t \right) dx.$$

Since the test function $\phi$ is arbitrary, the two weak forms give the push-forward velocity relation

$$v_{2}\left( T\left( x \right),t \right)=\left( \nabla T\left( x \right) \right)v_{1}\left( x,t \right) \rho\text{-a.e.}$$

The secondary space energy can therefore be written in primary space coordinates as

$$\int_{Y} \xi\left( \parallel v_{2}\parallel^{2}+\alpha h^{2} \right)dy=\int_{X} \rho\left( \parallel v_{2}\left( T\left( x \right),t \right)\parallel^{2}+\alpha g^{2} \right)dx.$$

$$\int_{X} \rho\left( \parallel v_{2}\left( T\left( x \right),t \right)\parallel^{2}+\alpha g^{2} \right)dx=\int_{X} \rho\left( v_{1}^{\top}\left( \nabla T \right)^{\top}\left( \nabla T \right)v_{1}+\alpha g^{2} \right)dx.$$

Substituting this identity into the original cost functional yields the equivalent primary space expression

$$\int_{0}^{1} \left[ \beta\int_{X} \rho\left( \parallel v_{1}\parallel^{2}+\alpha g^{2} \right)dx+\left( 1-\beta\right)\int_{Y} \xi\left( \parallel v_{2}\parallel^{2}+\alpha h^{2} \right)dy \right]dt$$

$$\int_{0}^{1} \int_{X} \rho\left( v_{1}^{\top}\left[ \beta I+\left( 1-\beta\right)\left( \nabla T \right)^{\top}\left( \nabla T \right) \right]v_{1}+\alpha g^{2} \right) dx dt.$$

This completes the proof.

Supplementary Note 4. Transport map pretraining and convergence

MOFI uses a differentiable transport map between omics before and during multi-omics dynamics fitting. The implemented mapper follows a spliced autoencoder with omics specific encoders and decoders. Training and validation losses were monitored for the HSPCs RNA–protein and human brain organoid RNA–ATAC datasets, including reconstruction within each omics, reconstruction across omics, and when enabled, latent space consistency, cross reconstruction terms.

For HSPCs, the transport map objective decreased rapidly during early epochs and then reached a plateau (Supplementary Fig. 1A). RNA to RNA, protein to protein, RNA to protein and protein to RNA reconstruction losses decreased and stabilized, with similar training and validation trends across the major loss components.

For the human brain organoid dataset, total loss decreased during early training and plateaued as the learning rate was reduced (Supplementary Fig. 1B). RNA to RNA, ATAC to ATAC, RNA to ATAC and ATAC to RNA losses stabilized with similar training and validation trends.

Supplementary Note 5. Training pseudocode, implementation settings and hyperparameter sensitivity

MOFI training follows three implementation stages: pretraining the paired transport map between omics, initializing the velocity and growth dynamics in the primary latent space, and refining the dynamics with the paired secondary omics. Let $\mathcal{K}_{1}$ and$\mathcal{K}_{2}$ denote the sets of time indices with observed primary and secondary snapshots, respectively. In the dynamics stages, each epoch begins by sampling particles from the earliest observed primary snapshot; the particles are then propagated sequentially across adjacent time intervals, optimized interval by interval, and detached before entering the next interval. The primary space OT, mass and energy terms are evaluated along the primary observation sequence, whereas the secondary space OT term is added only when the target index satisfies k + 1 ∈ $\mathcal{K}_{2}$. This conditional secondary loss is what allows the same training procedure to accommodate asymmetric temporal sampling such as the five versus three simulation design. Algorithm 1 summarizes the procedure, while Supplementary Tables 14 and 15 provide the recommended starting settings and the schedules used for the reported analyses in each dataset.

Algorithm 1. MOFI training

**Notation.** The notation is aligned with the main text. $X_{k}\mathcal{\subset X}$ and $Y_{k}\mathcal{\subset Y}$ are the observed cell states in the primary and secondary omics at time $t_{k}$. $\mathcal{K}_{1}$ and $\mathcal{K}_{2}$ are the corresponding observed time index sets, allowing $\mathcal{K}_{2}\subseteq\mathcal{K}_{1}$. $T_{\theta}^{a\to b}$ denotes a map from omics $a$ to omics $b$; $T_{\theta}^{1\to2}$ is the forward map between omics used during dynamics fitting. $v_{1\theta}\left( x,t \right)$ is the primary omics velocity field, $g_{\theta}\left( x,t \right)$ is the growth rate, and $w>0$ is the particle mass. The implementation evolves $\log w$ for numerical stability, with $w=exp\left( \log w \right)$. $e$ denotes the accumulated dynamical energy state, $J_{\theta}\left( x \right)=\nabla_{x}T_{\theta}^{1\to2}\left( x \right)$ is the Jacobian of the forward map, $B$ is the batch size, and $r_{k+1}=N_{k+1}/N_{0}$ is the observed relative population mass.

**Input:** primary snapshots $\{X_{k}{\}}_{k\in\mathcal{K}_{1}}$; secondary snapshots $\{Y_{k}{\}}_{k\in\mathcal{K}_{2}}$; ordered times $\{t_{k}\}$; batch size $B$; loss coefficients for each stage.

**Output:** trained map between omics $T_{\theta}$ and fitted dynamics $v_{1\theta},g_{\theta}$.

**Pretrain: pretrain the paired map between omics** $T_{\theta}$ **using MSE.**

Initialize the map subnetworks $T_{\theta}^{1\to1}$, $T_{\theta}^{1\to2}$, $T_{\theta}^{2\to1}$ and $T_{\theta}^{2\to2}$.

**for** each map training epoch **do**

    Sample paired cells $\left( X,Y \right)$ observed in both omics.

    Compute the four reconstructions

$$\hat{X}_{11}=T_{\theta}^{1\to1}\left( X \right), \hat{Y}_{12}=T_{\theta}^{1\to2}\left( X \right), \hat{X}_{21}=T_{\theta}^{2\to1}\left( Y \right), \hat{Y}_{22}=T_{\theta}^{2\to2}\left( Y \right).$$

$$X_{\mathrm{latent}}= {Encoder}_{1\theta}(X), Y_{\mathrm{latent}} ={Encoder}_{2\theta}(Y)$$

    Evaluate the map objective

$$\mathcal{L}_{\mathrm{Map}}=\alpha_{\mathrm{same}}\mathcal{L}_{\mathrm{same}}+\alpha_{\mathrm{cross}}\mathcal{L}_{\mathrm{cross}}+\alpha_{c\mathrm{onsistency}}\mathcal{L}_{c\mathrm{onsistency}}.$$

$$\mathcal{L}_{\mathrm{same}}=MSE\left( \hat{X}_{11},X \right)+MSE\left( \hat{Y}_{22},Y \right),$$

$$\mathcal{L}_{\mathrm{cross}}=MSE\left( \hat{Y}_{12},Y \right)+MSE\left( \hat{X}_{21},X \right).$$

$$\mathcal{L}_{c\mathrm{onsistency}}=MSE\left( X_{\mathrm{latent}},Y_{\mathrm{latent}} \right).$$

    Update the parameters of $T_{\theta}$ by Adam; monitor validation MSE and retain the selected checkpoint.

**end for**

Retain the checkpoint for $T_{\theta}$. *No particle mass or particle weight is used in Pretrain.*

**Stage 1: initialize** $v_{1\theta}$ **and** $g_{\theta}$ **in the primary omics space.**

Initialize the velocity network $v_{1\theta}$ and growth network $g_{\theta}$.

**for** each primary space training stage $s$ and each epoch **do**

    Sample the initial particle batch and initialize uniform mass:

$$x_{0}\leftarrow Sample\left( X_{0},B \right), \log w_{0}\leftarrow log\left( 1/B \right) \mathbf{1}_{B}.$$

**for** $k=0,\ldots,K-1$ **do**

        Sample the next observed target in the primary omics and its relative population mass:

$$X_{k+1}^{\mathrm{tar}}\leftarrow Sample\left( X_{k+1},B \right), r_{k+1}=\frac{N_{k+1}}{N_{0}}.$$

        Propagate the current particles over $\left[ t_{k},t_{k+1} \right]$:

$$\left( x_{k+1},logw_{k+1},e_{k+1} \right)\leftarrow NeuralODE\left( x_{k},logw_{k},0;t_{k},t_{k+1} \right).$$

        During integration,

$$\frac{dx}{dt}=v_{1\theta}\left( x,t \right), \frac{d\log w}{dt}=g_{\theta}\left( x,t \right), \frac{de}{dt}=\left[ \frac{1}{2}\parallel v_{1\theta}\left( x,t \right)\parallel^{2}+g_{\theta}\left( x,t \right)^{2} \right]w, w=e^{\log w}.$$

        Let $w_{k+1}=exp\left( \log w_{k+1} \right)$ and $e_{k+1}=\sum_{i=1}^{B} e_{i,k+1}$. Compute

$$\mathcal{L}_{k}=\lambda_{OT,1}\mathcal{L}_{\mathrm{OT}}\left( x_{k+1},X_{k+1}^{\mathrm{tar}};w_{k+1} \right)+\lambda_{\mathrm{mass}}\mathcal{L}_{\mathrm{Mass}}\left( x_{k+1},w_{k+1};X_{k+1}^{\mathrm{tar}},r_{k+1} \right)+\lambda_{Energy}e_{k+1}.$$

        Zero gradients; backpropagate $\mathcal{L}_{k}$; update $v_{1\theta}$ and $g_{\theta}$.

        Detach $x_{k+1}$ and $\log w_{k+1}$ and use them as the initial particle state for the next interval.

**end for**

**end for**

**Stage 2: refine the dynamics using the secondary omics wherever it is observed.**

Load the map for the selected direction using the checkpoint chosen by validation $T_{\theta}$. In standard fitting the map is held fixed unless joint updating is explicitly enabled.

**for** each paired multi-omics training stage $s$ and each epoch **do**

    Initialize $x_{0}$ and $\log w_{0}$ as in Stage 1; the interval energy state is reset to zero for each Neural ODE step.

**for** $k=0,\ldots,K-1$ **do**

        Sample $X_{k+1}^{\mathrm{tar}}$ and compute $r_{k+1}=N_{k+1}/N_{0}$.

        Propagate $\left( x_{k},logw_{k},0 \right)$ over $\left[ t_{k},t_{k+1} \right]$. During integration,

$$\frac{dx}{dt}=v_{1\theta}\left( x,t \right), \frac{d\log w}{dt}=g_{\theta}\left( x,t \right),$$

$$\frac{de}{dt}=\left\{ \frac{1}{2}\left[ \parallel v_{1\theta}\left( x,t \right)\parallel^{2}+\alpha_{\mathrm{multi}}\parallel J_{\theta}\left( x \right)v_{1\theta}\left( x,t \right)\parallel^{2} \right]+g_{\theta}\left( x,t \right)^{2} \right\}w, J_{\theta}\left( x \right)=\nabla_{x}T_{\theta}^{1\to2}\left( x \right).$$

        The induced velocity in the secondary omics is

$$v_{2\theta}\left( y,t \right)=J_{\theta}\left( x \right)v_{1\theta}\left( x,t \right), y=T_{\theta}^{1\to2}\left( x \right).$$

        Initialize the interval loss with the OT, mass and energy terms in the primary omics:

$$\mathcal{L}_{k}=\lambda_{OT,1}\mathcal{L}_{\mathrm{OT}}\left( x_{k+1},X_{k+1}^{\mathrm{tar}};w_{k+1} \right)+\lambda_{\mathrm{mass}}\mathcal{L}_{\mathrm{Mass}}\left( x_{k+1},w_{k+1};X_{k+1}^{\mathrm{tar}},r_{k+1} \right)+\lambda_{Energy}\bar{e}_{k+1}.$$

**if** $k+1\in\mathcal{K}_{2}$ **then**

$$Y_{k+1}^{\mathrm{tar}}\leftarrow Sample\left( Y_{k+1},B \right), y_{k+1}=T_{\theta}^{1\to2}\left( x_{k+1} \right),$$

$$\mathcal{L}_{k}\leftarrow\mathcal{L}_{k}+\lambda_{OT,2}\mathcal{L}_{\mathrm{OT}}\left( y_{k+1},Y_{k+1}^{\mathrm{tar}};w_{k+1} \right).$$

**end if**

        Zero gradients; backpropagate $\mathcal{L}_{k}$; update the trainable dynamics parameters.

        Detach $x_{k+1}$ and $\log w_{k+1}$ before the next interval.

**end for**

**end for**

**return** $T_{\theta}$, $v_{1\theta}$ and $g_{\theta}$.

**Symbol meanings used in the loss.** $\lambda_{OT,1}$ weights distribution matching in the primary omics; $\lambda_{OT,2}$ weights distribution matching in the secondary omics when that snapshot is observed; $\lambda_{\mathrm{mass}}$ weights local/global population mass matching; $\lambda_{Energy}$ weights the dynamical energy; and $\alpha_{\mathrm{multi}}$ controls the contribution of the mapped velocity in the secondary omics to the secondary space dynamical energy.

In the implementation, the mapper and dynamics have distinct loss parameters. The transport map is a spliced autoencoder trained with the four direct reconstruction terms. When enabled, it also includes latent space consistency and cross reconstruction terms. During dynamics fitting, $\alpha_{multi}$ controls the mapped velocity contribution to the dynamical energy through ODEFunc, whereas $\lambda_{OT,2}$ weights the secondary space distribution matching loss. The transport map is fixed during standard dynamics fitting unless joint updating is explicitly enabled.

The pseudocode follows the implementation paths used for transport map training and staged neural ODE optimization[5]. Recommended starting values are separated from the final configurations used for each dataset: Supplementary Table 14 provides a reproducible starting regime, whereas Supplementary Table 15 records the saved settings associated with the reported analyses.

Hyperparameter sensitivity analysis

Finally, we examined the sensitivity of MOFI to several hyperparameters controlling optimal transport matching, dynamical energy and alignment between omics. We focused on five hyperparameters that directly affect the inferred dynamics: the primary space OT weight, the relative contributions of the dynamical energies in the primary and secondary spaces, the latent space consistency loss, and the secondary space OT weight. Each parameter was varied over a broad range of 0.1×, 0.5×, 1×, 2× and 10× relative to its original pipeline value, and MOFI was retrained with three independent random seeds at each setting. The configuration used in the reported analysis served as the 1× reference.

We first evaluated the three dynamical hyperparameters using the simulation benchmark in Fig. 2. The primary space OT weight controls the contribution of distribution matching in the primary state space. MOFI showed relatively small changes in W1 reconstruction error over the tested range, with the reference setting (Supplementary Table 15) lying within the region of low error in both the primary space and the analytically mapped secondary space (Supplementary Fig. 2A, B). Very small or large OT weights modestly increased the reconstruction error, indicating that sufficient distribution matching is required while excessive weighting provides no additional benefit.

The relative balance between the dynamical energies in the primary and secondary spaces showed a stronger effect (Supplementary Fig. 2C, D). Reducing or moderately increasing the primary space dynamical energy contribution produced similar reconstructed distributions, whereas excessive weighting of the primary space dynamical energy progressively increased W1 error, particularly at 10×. Consistently, deterministic trajectories initialized from the same simulated cells remained similar around the reference setting but diverged more substantially under the largest perturbation (Supplementary Fig. 3). These results indicate that MOFI tolerates moderate changes in the dynamical energy balance but that strongly favoring one omics can distort the jointly inferred dynamics.

The secondary space dynamical energy coefficient showed a similar but weaker pattern. Reconstruction remained largely stable from 0.1× to 2×, whereas the 10× setting increased W1 distance in both state spaces (Supplementary Fig. 2E,F). The corresponding trajectory analysis likewise showed limited changes around the reference configuration (Supplementary Fig. 3). Together, the simulation experiments indicate that MOFI is relatively insensitive to moderate perturbations of the OT and dynamical energy coefficients, while extreme imbalance between dynamical constraints can degrade reconstruction.

We next examined two parameters governing coordination between omics using the developing human cerebral cortex RNA–ATAC dataset. The latent space consistency loss controls how strongly the RNA and ATAC latent representations are aligned during learning of the Map between omics. W1 reconstruction changed only modestly across the tested range in both the primary RNA space and the mapped ATAC space (Supplementary Fig. 4A, B), suggesting that the inferred population dynamics are not strongly dependent on the exact alignment loss weight within this range.

The secondary space OT weight determines how strongly the mapped secondary distribution contributes to fitting the joint dynamics. Moderate changes in this coefficient produced comparable reconstruction errors, whereas a large 10× weight substantially increased W1 distance in the primary RNA space (Supplementary Fig. 4C, D). This result suggests that excessive emphasis on secondary space distribution matching can perturb the primary dynamics.

Supplementary Note 6. HSPCs RNA–protein analyses

We analyzed the HSPCs RNA–protein dataset to characterize continuous hematopoietic differentiation across the paired RNA and protein spaces and to examine perturbation responses associated with lineage. The dataset contains paired transcriptomic and surface protein measurements at four experimental time points; normalized model time 0–3 follows the ordered experimental progression used in the main text. The supplementary analyses cover trajectories, cell type fate probabilities, lineage programs, velocity and potential landscapes, perturbation responses and marker dynamics.

Downstream HSPCs trajectories were simulated with time step 0.05 and evaluated on a normalized model time grid from 0.25 to 3.00 in increments of 0.25 using seed 42. Generated states were recovered by PCA projection. Cell type probabilities were obtained with a linear classifier trained in the joint RNA–protein space on observed annotations. MkP was the target lineage for path analyses, and fate redistribution summaries included EryP, MasP, MkP, MoP and NeuP.

Continuous RNA–protein trajectory reconstruction and lineage program dynamics

MOFI reconstructed a continuous developmental process from the discrete HSPCs RNA–protein snapshots (Supplementary Fig. 5A). In RNA PCA space, generated trajectories progressed from early progenitor states toward differentiated branches while preserving the fan shaped geometry of the observed hematopoietic manifold. Mapping the same generated cells into protein space produced a concordant branching structure and preserved the temporal organization of paired RNA–protein states.

Applying the joint space cell type classifier to cells generated by MOFI across continuous time yielded a progressive redistribution among erythroid, neutrophil, megakaryocyte, mast cell and monocyte progenitor states (Supplementary Fig. 5B).

Lineage molecular programs were summarized over the reconstructed time course using predefined marker sets informed by established hematopoietic lineage markers: megakaryocyte (ITGA2B, GP1BA, NFE2, GATA1, PPBP and PF4), erythroid (KLF1, GATA1, TFRC, ALAS2, HBA1 and HBB), and myeloid (MPO, LYZ, ELANE, S100A8, S100A9 and CSF3R). The reconstructed program heatmap showed coordinated temporal remodeling across lineages (Supplementary Fig. 5C), with corresponding RNA and paired multi-omics summaries (Supplementary Fig. 5D, E).

Perturbation response, velocity fields and potential landscapes

Coordinated molecular responses were examined across the paired state space. Perturbation of ITGA2B, a megakaryocyte-associated gene, produced a graded response along the ITGA2B–GP1BA axis across the reconstructed time course (Supplementary Fig. 6A).

RNA space velocity streamlines pointed from early regions enriched for progenitors toward differentiated branches, and the mapped protein space velocity field showed the same global directionality (Supplementary Fig. 6B, C). Potential landscapes in RNA and protein spaces contained basins along terminal regions (Supplementary Fig. 6D,E).

Potential landscape construction. The potential landscapes in Supplementary Fig. 6D,E and Supplementary Fig. 13A, B were obtained by fitting a scalar neural network $U_{\varphi}\left( \mathbf{x},t \right)$ to the learned velocity field in the displayed 2D coordinates. The potential network was optimized so that the negative spatial gradient of its scalar output matched the projected velocity field, $v\left( \mathbf{x},t \right)$ ≈ - ${-\nabla}_{X}U_{\varphi}\left( \mathbf{x} \right)$, over the sampled states used for the corresponding visualization. The fitted scalar field was then evaluated over the displayed 2D domain to generate the potential landscape; regions of low potential were interpreted as dynamical basins.

Marker set perturbation sweeps

Using the in silico perturbation procedure described in the main Methods, we evaluated seven predefined HSPCs marker sets (Supplementary Table 16). Mk_union is the megakaryocyte marker set used for the perturbation analysis focused on MkP in the main text. Each group was evaluated at z scores of -8, -6, -4, -2, 0, 2, 4, 6 and 8. Endpoint occupancy changes were quantified for EryP, MasP, MkP, MoP and NeuP; temporal fate curves were examined at z = -8, 0 and 8.

Perturbations of mast cell marker sets produced model predictions that depended on perturbation sign. Positive perturbation of Mas_core_2 and Mas_union increased mast cell occupancy, and negative perturbation reduced it (Supplementary Fig. 7A,B). Mas_core_1 showed the same overall direction, with the strongest reciprocal change involving monocyte occupancy (Supplementary Fig. 7D).

Perturbations of megakaryocyte marker sets shifted predicted occupancy toward the MkP lineage. Positive perturbation of Mk_union and Mk_core_1 increased MkP occupancy, and negative perturbation reduced it (Supplementary Fig. 7C, E). Mk_core_2 showed the same dependence on perturbation sign (Supplementary Fig. 8A). Ery_core_1 produced an analogous shift in erythroid occupancy, as predicted by the model (Supplementary Fig. 8B).

Molecular trajectory reconstruction

Marker dynamics were evaluated in RNA and protein. The RNA panel contained ITGA2B, GP1BA, TFRC, KLF1, GATA1, MPO, LYZ and HMGCR, and the protein panel contained CD42b and CD71. Supplementary Fig. 8C shows reconstructed CD71 and CD42b trajectories with observed measurements at sampled time points.

RNA trajectories for MPO, LYZ, TFRC, ITGA2B, GATA1 and KLF1 varied smoothly across the reconstructed differentiation process and remained aligned with observed measurements at sampled times (Supplementary Fig. 8D). The RNA and protein panels represent myeloid, erythroid and megakaryocyte molecular programs.

Temporal fate probability curves were evaluated under Mas_union perturbation at z = -8, 0 and 8. Negative and positive perturbations produced opposite temporal shifts in lineage probabilities relative to the neutral trajectory (Supplementary Fig. 9A–C), showing how the endpoint redistribution emerged over model time. The temporal curves complement the endpoint analysis by showing the sequence of predicted fate redistribution along the HSPCs differentiation trajectory.

Supplementary Note 7. Human brain organoid RNA–ATAC analyses

We analyzed the human brain organoid RNA–ATAC dataset across its developmental time course, spanning embryoid body states through neuroectodermal and neuroepithelial intermediates to later organoid states. In the original study, scRNA-seq and scATAC-seq were generated using separate assays and subsequently matched to construct multi-omic metacells.[6] The established RNA–ATAC correspondence was retained and used directly as input to MOFI. The supplementary analyses cover continuous trajectories, cell-type fate dynamics, developmental programs, marker reconstruction, RNA–ATAC temporal shift analysis and reverse-direction training.

Downstream inference used seed 42 and simulation step 0.05. Generated states were sampled on a normalized time grid from 0.5 to 9.0 in increments of 0.5, with at most 2,000 cells per sampling pass, and recovered by PCA projection. Cell type probabilities were obtained with a linear classifier trained in the joint RNA–ATAC space. Fate summaries used EB, nect, nepi and organoid annotations.

Latent reconstruction and gene decoding

The dynamics model used 9,894 paired cells across the ordered organoid time course. The 50D RNA and ATAC latent coordinates used for dynamics fitting were retained throughout downstream trajectory generation, and gene summaries were recovered from the corresponding PCA representations.

Continuous trajectory reconstruction and cell type fate dynamics

MOFI reconstructed continuous developmental trajectories in both RNA and ATAC spaces (Supplementary Fig. 10A, B). The RNA trajectory followed a broad arc from early EB cells through neuroepithelial and neuroectodermal intermediates toward late organoid states. The ATAC gene activity trajectory retained the same temporal ordering across model time.

Cell type fate probabilities showed sequential redistribution from states enriched for EB cells to neuroepithelial and neuroectodermal intermediates, followed by increasing organoid occupancy at late model times (Supplementary Fig. 10C,E). The heatmap representation shows the same ordered compositional change across continuous time.

Program dynamics and marker reconstruction

Three predefined RNA programs, selected to represent established pluripotency, neuroectodermal/neuroepithelial transition and neural maturation states, were summarized over the reconstructed time course: early pluripotency (POU5F1, TDGF1, LIN28A, EPCAM, FOXD3-AS1 and L1TD1), transition neuroectoderm/neuroepithelium (PRTG, IGFBP5, PRSS23, PTN, CLDN6, BST2 and GPC3), and neural maturation (STMN2, GPM6A, GAP43, ELAVL4, MAPT, NR2F1 and HES6). The early pluripotency program decreased over developmental time, the transition program was highest at intermediate stages, and the neural maturation program increased later in the trajectory (Supplementary Fig. 10D,F).

Marker temporal profiles were summarized from the reconstructed trajectories. The RNA panel included L1TD1, POU5F1, GPC3, ID1, LIN28A, IGFBP5 and NR2F1 (Supplementary Fig. 11A), and the ATAC gene activity panel included ASIC2, ROR1, TSPEAR, CUBN, GPM6A, PRTG and GCNT2 (Supplementary Fig. 12A).

An illustrative temporal-shift analysis of selected RNA–ATAC marker pairs

Temporal shifts were evaluated for three predefined matched RNA–ATAC pairs: ABCA4, NEUROG2 and PAX8 (Supplementary Fig. 12B). For each pair, similarity between the RNA and ATAC temporal profiles was evaluated across candidate temporal offsets, and the offset yielding the highest similarity was taken as the estimated shift. Pairs with a maximal similarity greater than 0.6 were retained for interpretation. Across the retained pairs, the optimal shifts consistently indicated that changes in RNA expression occurred later than the corresponding changes in ATAC gene activity.

Reverse training with ATAC as the primary omics

For reverse fitting, the omics roles were exchanged and the transport map was retrained for the reversed direction using the same mapper architecture and nominal map settings. After exchanging the primary and secondary spaces, the model was trained using the dataset-specific settings reported in Supplementary Table 15.

The organoid model was also trained in the reverse direction, using ATAC as the primary omics and RNA as the mapped secondary omics. The reverse model produced structured ATAC and RNA landscapes, an ATAC velocity field and a joint trajectory aligned with the observed developmental manifold (Supplementary Fig. 13A–D). The predicted growth rate is consistent with the fact that cells undergo division in both the ATAC and RNA spaces (Supplementary Fig. 13E, F).

Supplementary Note 8. Developing human cerebral cortex RNA–ATAC analyses

We analyzed the two branches of the developing human cerebral cortex RNA–ATAC dataset, which is also used in the benchmark in Supplementary Note 2. The paired data span radial glial progenitors (RG), intermediate progenitor cells (IPC), fetal interneurons (IN-fetal), and CGE- and MGE interneuron states (IN-CGE and IN-MGE). The supplementary analyses cover forward RNA to ATAC reconstruction, temporal progression, reverse ATAC to RNA training and marker reconstruction.

Downstream inference used seed 42, a simulation step of 0.05 and normalized model times 1, 2, 3 and 4, with at most 2,000 generated cells per sampling pass. Generated states were recovered by PCA projection. Cell type probabilities were obtained using a linear classifier trained in the joint RNA–ATAC space across RG, IPC, IN-fetal, IN-CGE and IN-MGE states; IN-MGE was used as the target lineage for path analyses.

RNA and ATAC reconstruction

Using RNA as the primary omics and ATAC as the secondary omics, MOFI generated continuous trajectories in both latent spaces (Supplementary Fig. 14A, B). Three predefined cortical lineage programs, based on established markers associated with progenitor and interneuron states, summarized the reconstructed process: RG/IPC (HES5, VIM, EOMES, HES1, YAP1 and WWTR1), CGE interneuron (PROX1, CALB2, VIP, NR2F2, LEF1 and ALDH1A2), and MGE interneuron (SOX6, ERBB4, GAD1, GAD2, RELN and GLI3). The progenitor program was strongest early, and the interneuron programs increased later (Supplementary Fig. 14C).

Temporal visualization of cortical progression

Temporal PCA panels for RNA and ATAC show generated and observed states across the reconstructed progression (Supplementary Fig. 15). Generated cells moved from early regions dominated by RG and IPC toward later states enriched for interneurons, with intermediate model times occupying corresponding regions of the observed manifold.

Reverse training with ATAC as the primary omics

We repeated the analysis with ATAC as the primary omics and RNA as the secondary omics. The reverse model recovered trajectories in both ATAC and RNA spaces (Supplementary Fig. 16A, B). Predicted fate dynamics showed depletion of RG and IPC populations, expansion of IN-fetal states at intermediate times, and later emergence of IN-CGE and IN-MGE populations (Supplementary Fig. 16C, D).

For reverse fitting, the omics roles were exchanged and the transport map was retrained for the reversed direction using the same mapper architecture and nominal map settings. The velocity and growth architectures were unchanged. OT, mass and energy weights were initialized at the same order of magnitude as in the forward fit and were adjusted when needed to maintain stable optimization after exchanging the primary and secondary spaces.

Marker gene reconstruction

Marker reconstruction over time was evaluated in ATAC gene activity and RNA expression (Supplementary Fig. 17). The ATAC panel included ERBB4, RELN, GAD2, GAD1, LEF1 and HES5; the RNA panel included ERBB4, SOX6, GAD1, RELN, GAD2, VIM, PROX1 and EOMES. These markers represent progenitor and interneuron features in the predefined cortical programs.

The HC analyses show cortical progression in both RNA to ATAC and ATAC to RNA training directions, including fate redistribution from progenitors to interneurons and marker dynamics for cortical lineage genes.

Supplementary Note 9. GSE213152 kidney nephron RNA–ATAC analyses

We analyzed the GSE213152 kidney nephron RNA–ATAC time series as the fourth application to real data, together with the HSPCs, human brain organoid and developing human cortex analyses in Supplementary Notes 6–8. The analyzed branch contains 60,845 paired cells after construction of RNA and ATAC gene activity matrices, quality control and lineage filtering. Each omics was represented in a 50D PCA space for dynamics modeling.

Downstream trajectory generation used seed 42, simulation step 0.05, a model time grid from 0 to 4 in increments of 0.1, and 10,000 initial cells. Cell type summaries follow the six kidney states used in the main text: nephron progenitor cells (NPC), developing intermediate states (Dev), proximal tubule (PT), loop of Henle (LOH), distal nephron (DN), and podocytes (POD). RNA was treated as the primary space and ATAC gene activity as the mapped secondary space.

Quantitative reconstruction benchmark

GSE213152 is included in the reconstruction benchmark across four datasets in Supplementary Note 2, with complete pointwise RNA and ATAC values provided in Supplementary Tables 11 and 12. In the dedicated intermediate time point holdout analysis, MOFI had the lowest W1 for RNA and ATAC under paired holdout and for ATAC under the secondary omics holdout at each evaluated intermediate time (Supplementary Table 13).

Continuous RNA and ATAC trajectory visualization

The generated RNA states followed the ordered temporal observations across the RNA space (Supplementary Fig. 18). Mapping the same modeled progression into the ATAC gene space produced an ordered secondary space trajectory across the observed time points (Supplementary Fig. 19). These panels provide qualitative trajectory visualization complementary to the W1 reconstruction tests.

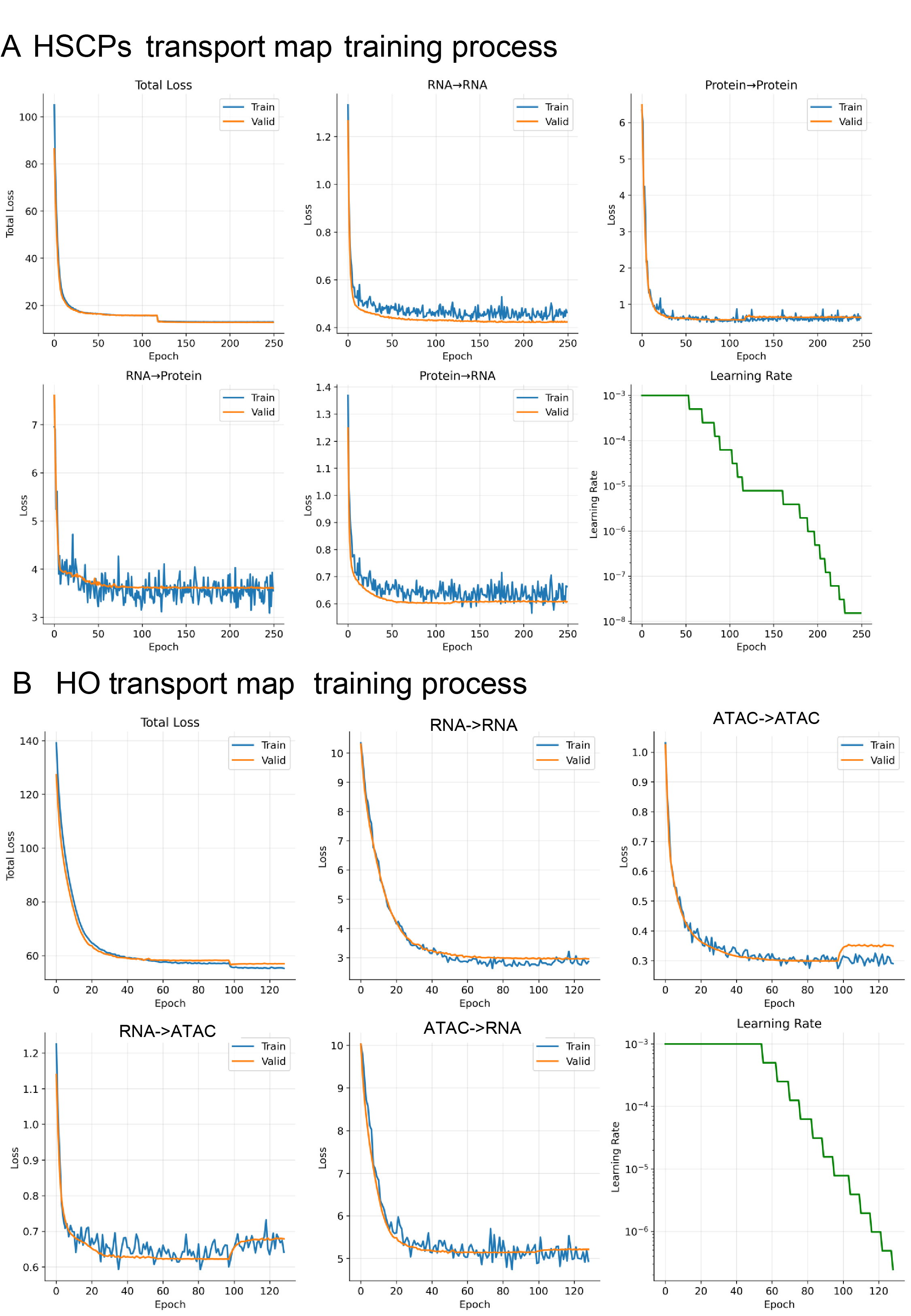

**Supplementary Fig. 1. Transport map training convergence for HSPCs and human brain organoid datasets.** A, HSPCs RNA–protein transport map training process, including total loss, reconstruction losses within each omics, reconstruction losses across omics and the learning rate schedule. B, Human brain organoid RNA–ATAC transport map training process with the same monitored loss components. Training and validation curves show rapid early loss reduction followed by stable plateaus.

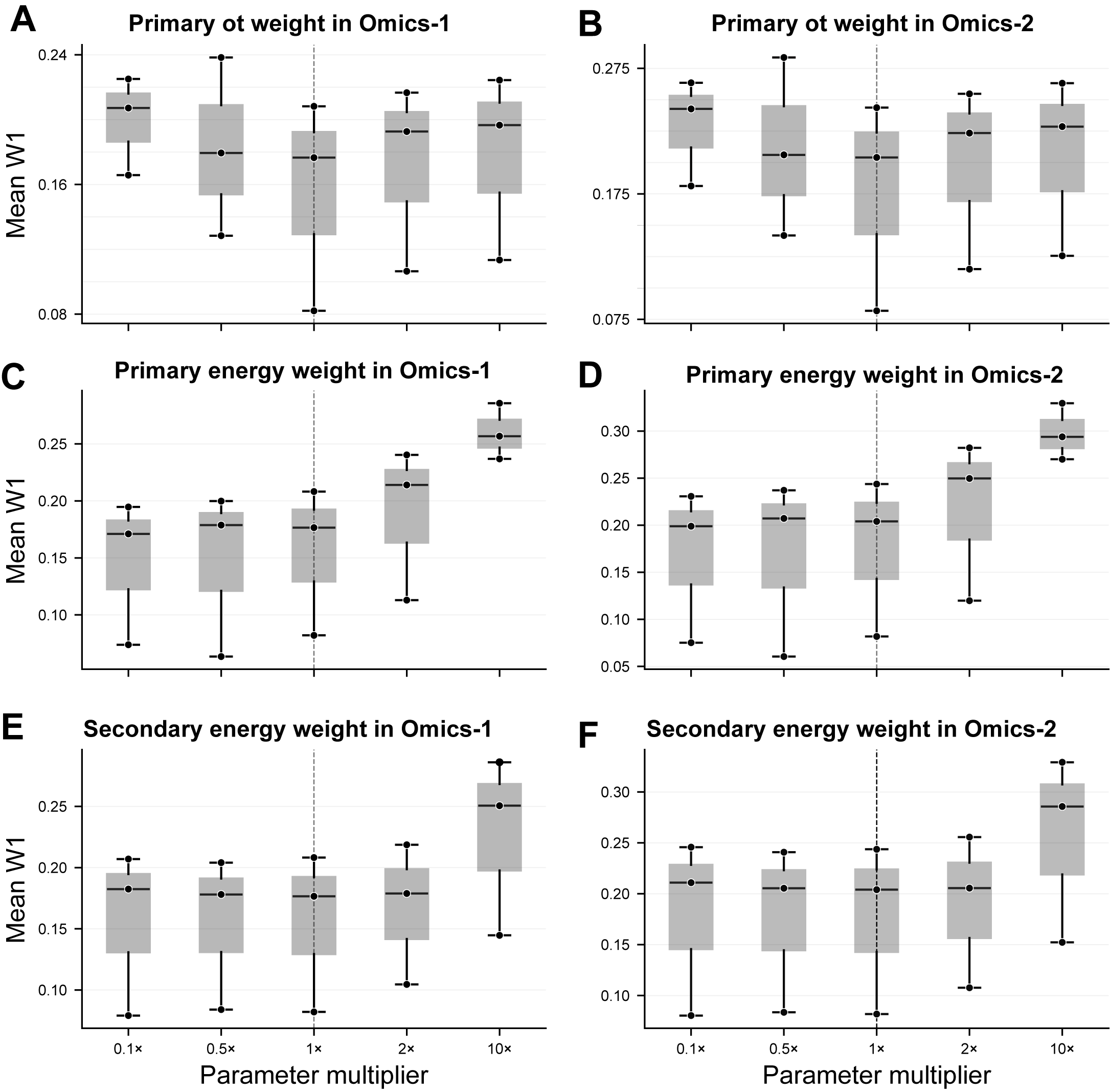

**Supplementary Fig. 2. Simulation sensitivity of W1 reconstruction to primary space OT and dynamical energy coefficients**. A, B, Primary space OT weight scan evaluated in the simulated primary 2D space (A) and analytically mapped secondary 3D space (B). C, D, Primary space dynamical energy scan in the primary (C) and mapped secondary (D) spaces. E, F, Secondary space dynamical energy scan in the primary (E) and mapped secondary (F) spaces. Each coefficient was evaluated at 0.1×, 0.5×, 1×, 2× and 10× of the original pipeline value. Boxes summarize three retrained time averaged W1 values from three retrained seeds and points show the individual seeds; the dashed line marks the original 1× setting.

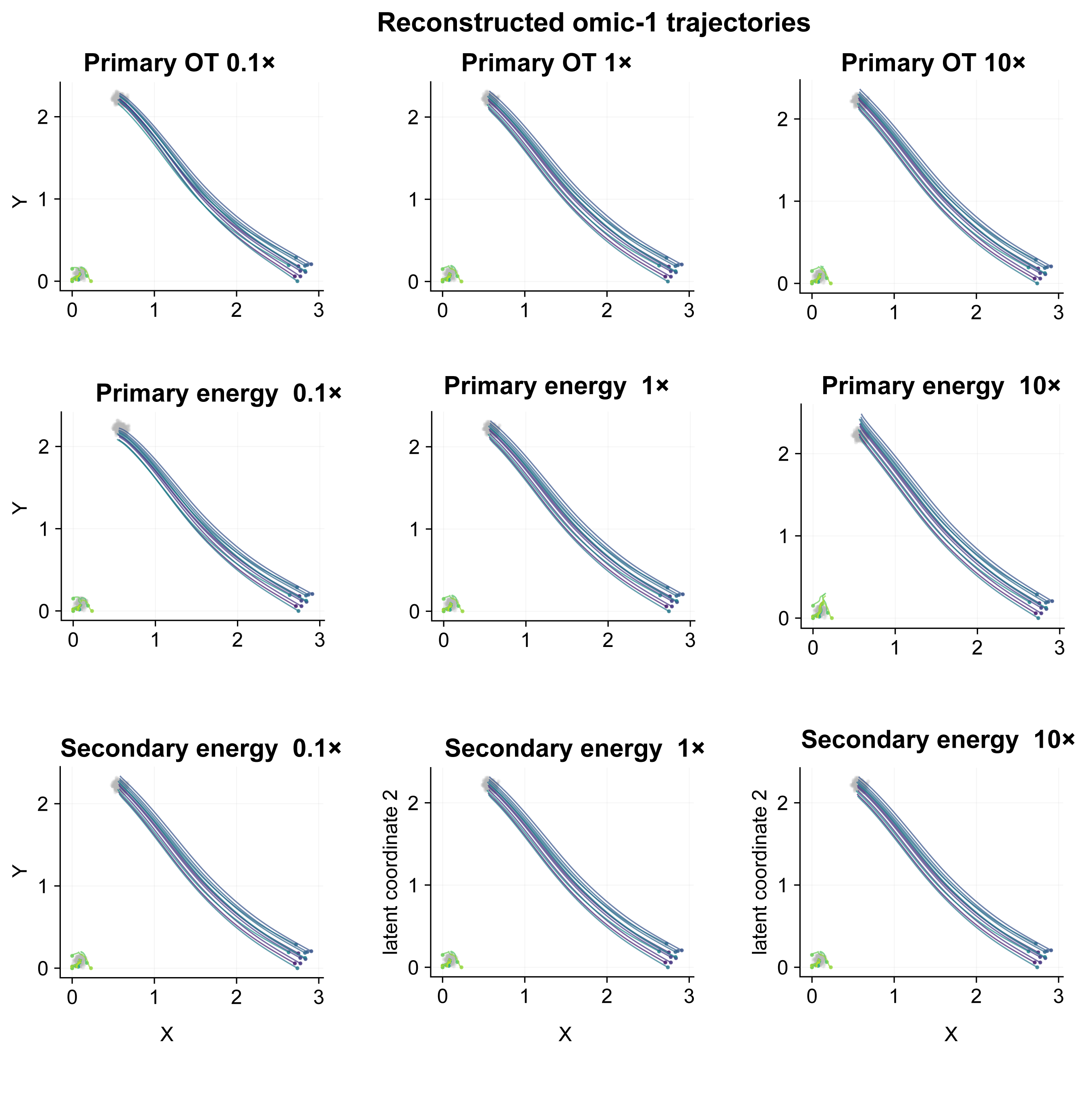

**Supplementary Fig. 3. Simulation trajectory sensitivity under OT and dynamical energy perturbations.** Rows show the primary space OT weight, primary space dynamical energy balance and secondary space dynamical energy weight; columns show 0.1×, 1× and 10× settings. Curves are deterministic primary space rollouts from the same 24 initial cells. The panels visualize changes in the fitted dynamics relative to the original 1× setting.

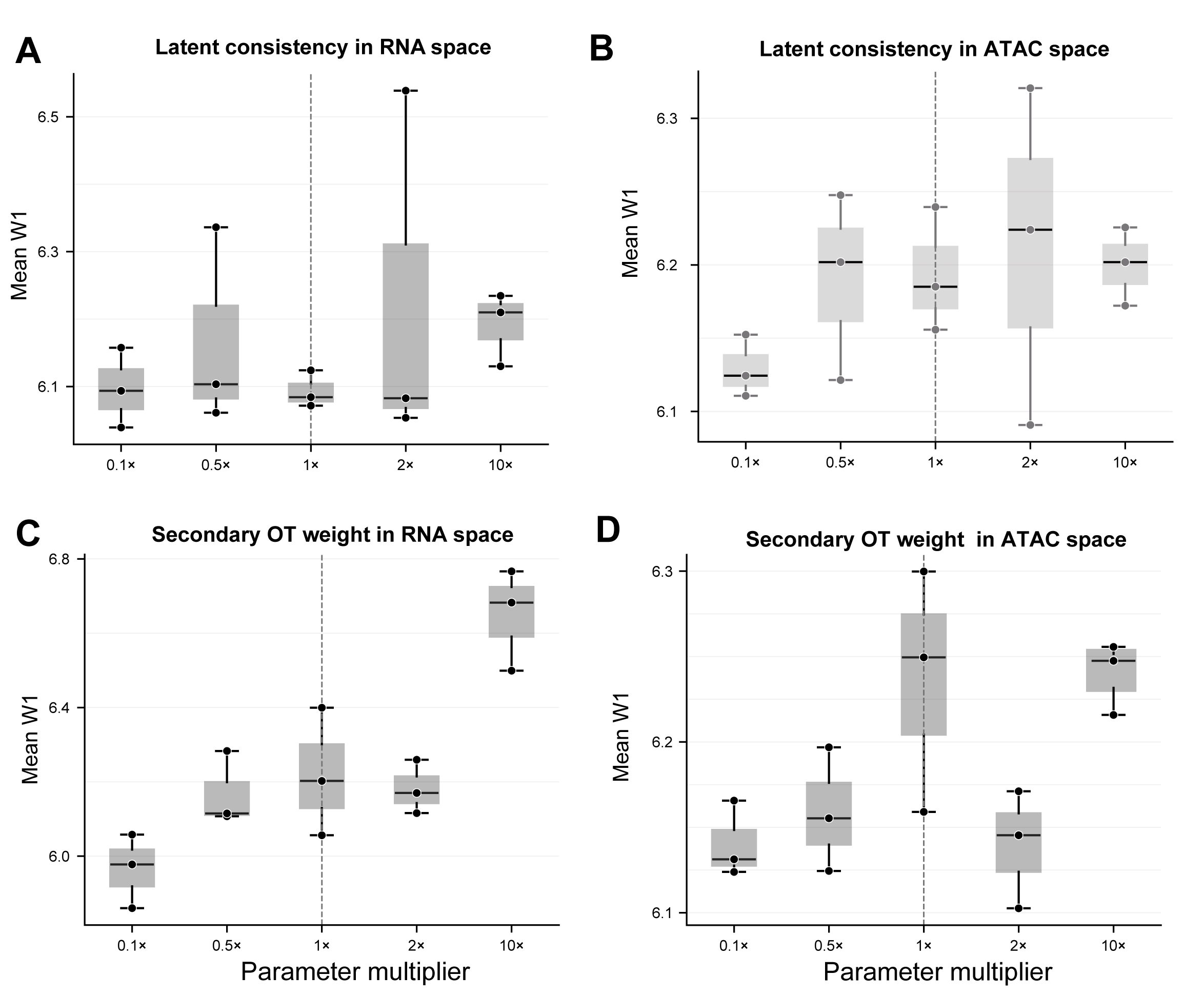

**Supplementary Fig. 4. W1 sensitivity on real data in the developing human cerebral cortex RNA–ATAC analysis.** A, B, latent space consistency loss scanned in the primary RNA PCA50 space (A) and the ATAC PCA50 space transformed by the Map trained for each factor (B). C, D, Secondary space OT weight scanned in the primary RNA space (C) and mapped ATAC space (D). Each point is the mean W1 value from one retrained seed over t = 1–4 (n = 3 seeds per multiplier); boxes summarize the three seeds and the dashed line marks the original 1× setting. Map alignment variants used their RNA→ATAC Maps retrained for each factor, whereas secondary space OT variants used the 1× Map retrained with the same seed to isolate the secondary space OT coefficient from Map variability.

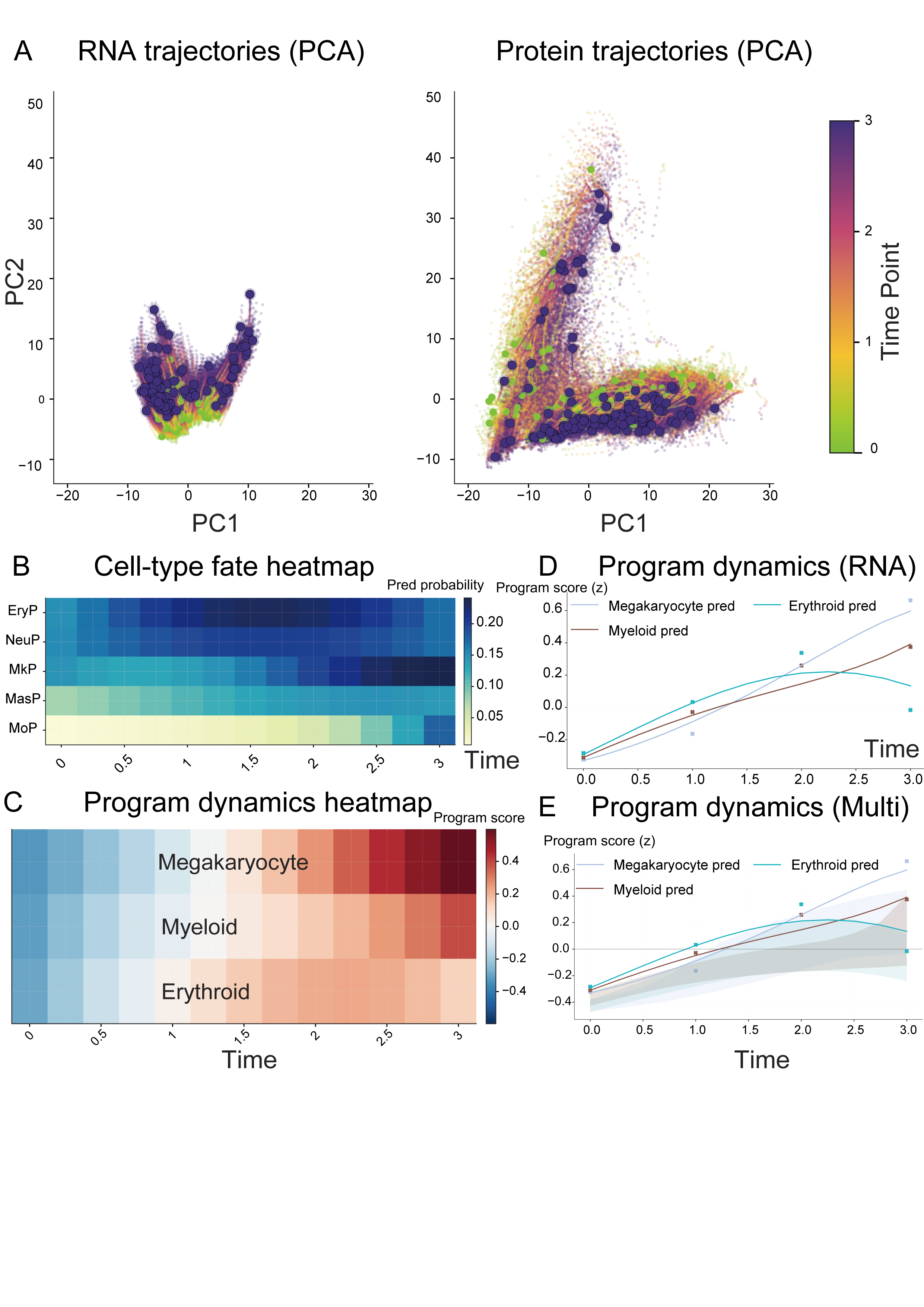

**Supplementary Fig. 5. Downstream analysis on HSPCs data.** A, continuous trajectories generated by MOFI projected into RNA PCA space and mapped into protein space, colored by normalized model time. B, Cell type fate probability heatmap across continuous time, computed by applying the joint space classifier to cells generated by MOFI. C, Lineage program dynamics across time for hematopoietic programs defined by marker sets. D, E, RNA (D) and paired multi-omics (E) program summaries across model time.

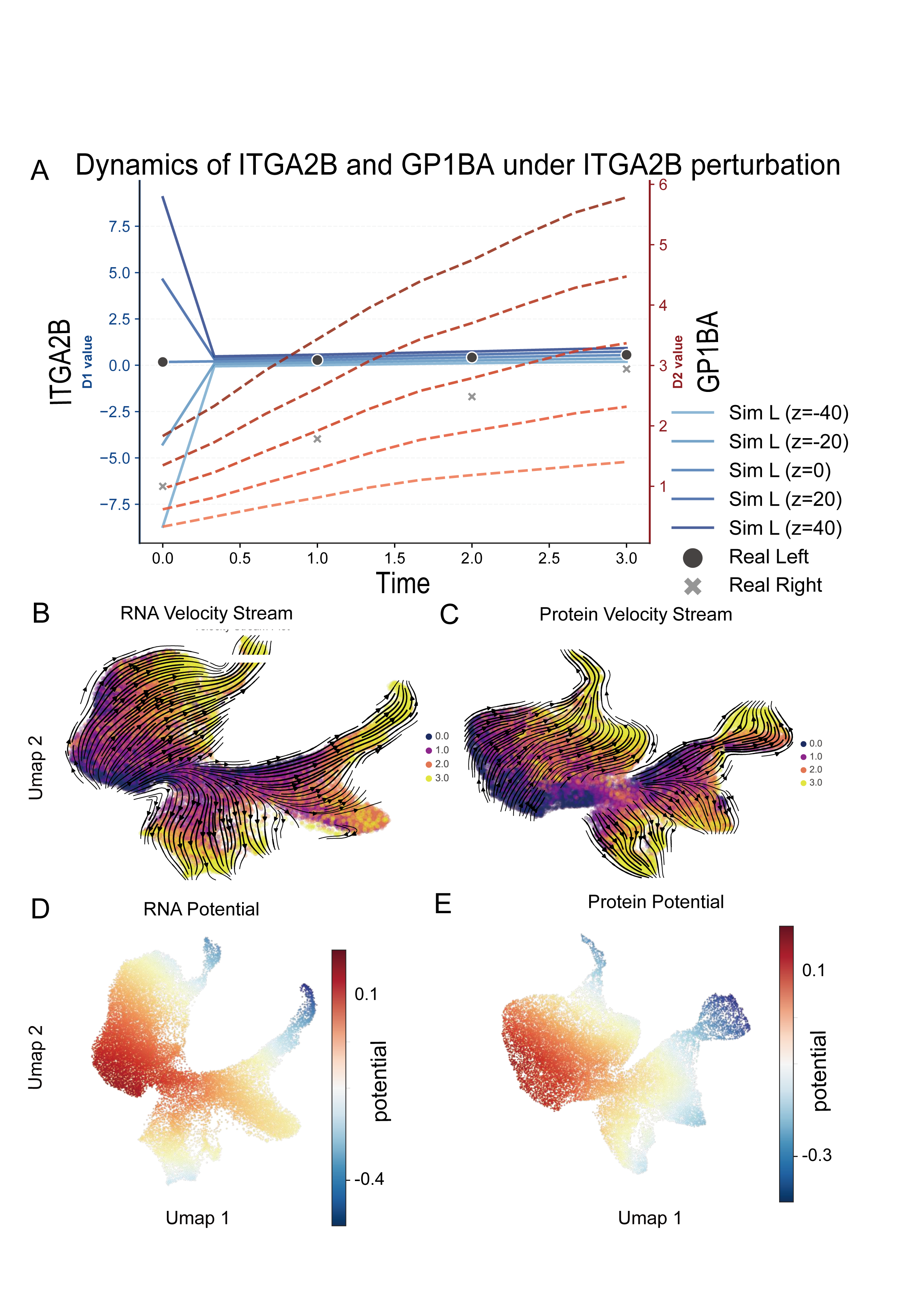

**Supplementary Fig. 6. RNA perturbation response, velocity fields and potential landscapes, related to Fig. 4.** A, Predicted ITGA2B–GP1BA temporal response under signed perturbation of ITGA2B. B, C, RNA space (B) and protein space (C) velocity stream plots inferred by MOFI. D, E, RNA space (D) and protein space (E) potential landscapes, showing structured terminal basins along the reconstructed differentiation manifold.

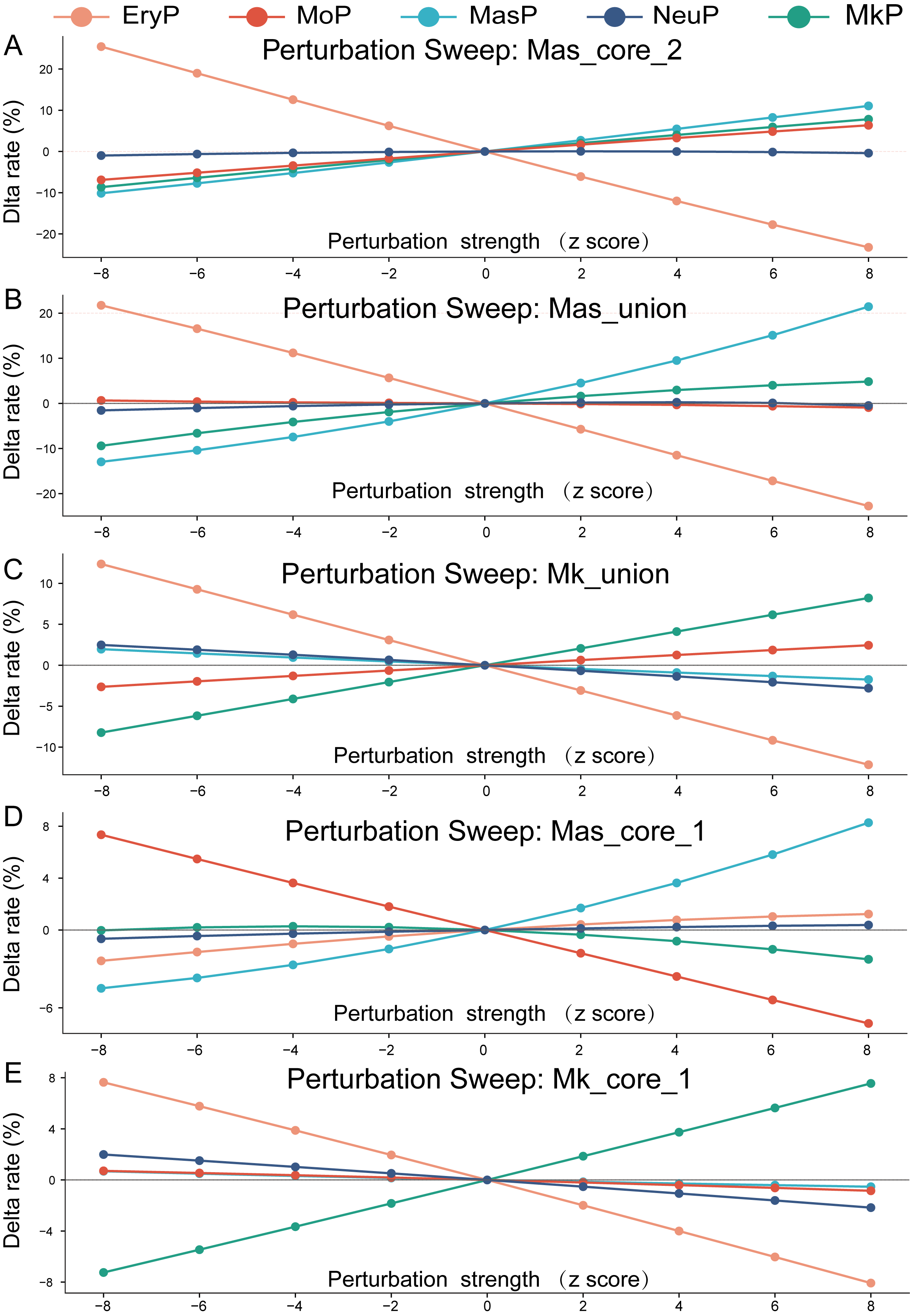

**Supplementary Fig. 7. Endpoint fate redistribution under mast cell and megakaryocyte marker set perturbations.** A–E, Change in final cell type occupancy under signed perturbation of Mas_core_2 (MS4A2, FCER1A, IL3RA) (A), Mas_union (TPSAB1, TPSB2, CPA3, MS4A2, FCER1A, IL3RA) (B), Mk_union (ITGA2B, GP1BA, PF4, PPBP) (C), Mas_core_1 (TPSAB1, TPSB2, CPA3) (D) and Mk_core_1 (ITGA2B, GP1BA) (E). Perturbations were evaluated from z = -8 to 8 in steps of 2; the zero point corresponds to the unperturbed simulation.

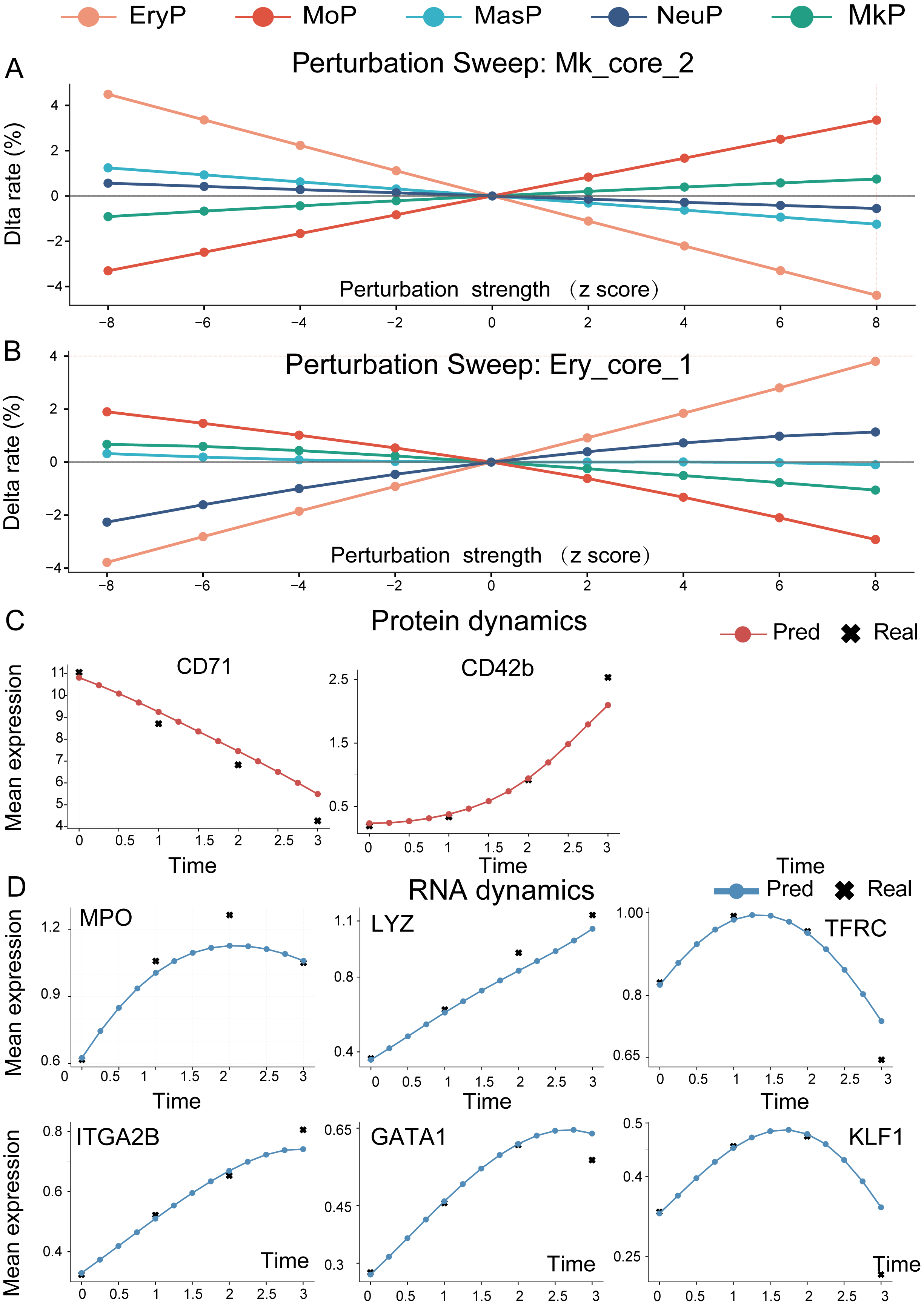

**Supplementary Fig. 8. Additional marker set perturbation sweeps and temporal dynamics of individual markers.** A, B, Final cell type occupancy changes under signed perturbation of Mk_core_2 (PF4, PPBP) (A) and Ery_core_1 (KLF1, GATA1, TFRC) (B). C, protein dynamics reconstructed by MOFI for CD71 and CD42b, with observed measurements shown at sampled time points. D, RNA dynamics reconstructed by MOFI for MPO, LYZ, TFRC, ITGA2B, GATA1 and KLF1.

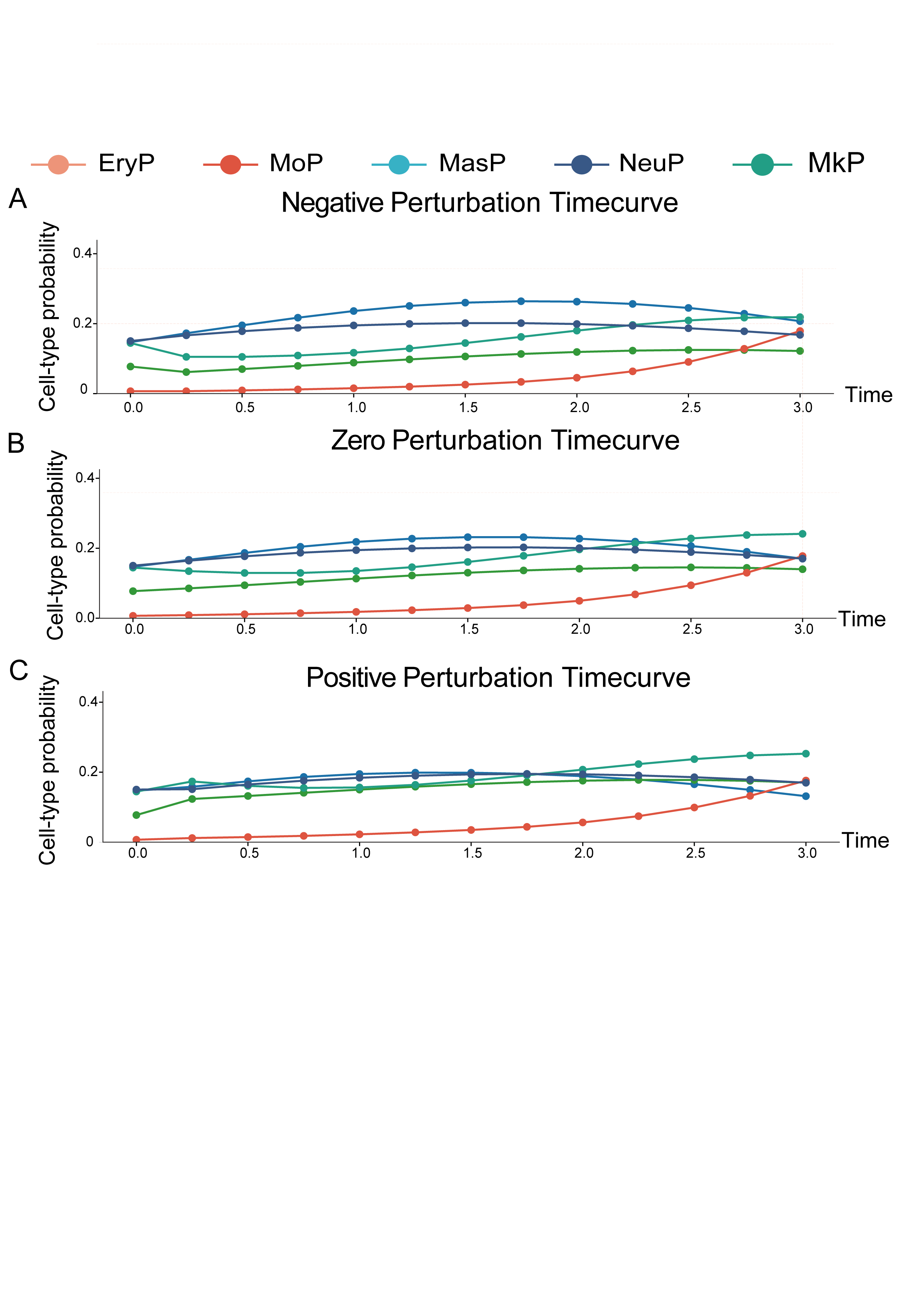

**Supplementary Fig. 9. Temporal fate probabilities under Mas_union perturbation.** A–C, Cell type probability trajectories under Mas_union perturbation (TPSAB1, TPSB2, CPA3, MS4A2, FCER1A and IL3RA) at z = -8 (A), 0 (B) and 8 (C), showing the temporal development of fate redistribution that depends on perturbation.

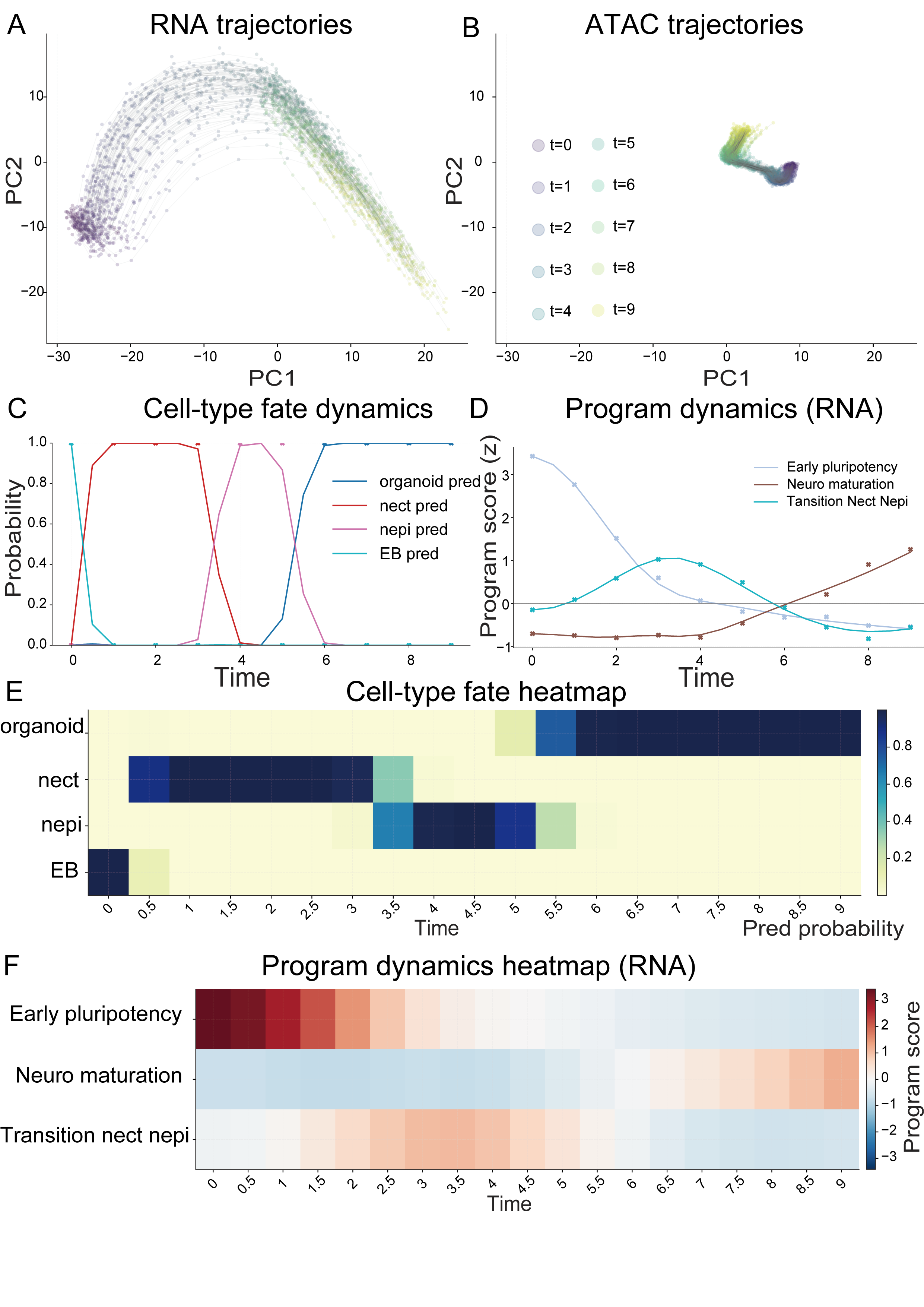

**Supplementary Fig. 10. Human brain organoid trajectory reconstruction and program dynamics.** A,B, continuous trajectories generated by MOFI projected into RNA space (A) and ATAC space (B), colored by normalized model time. C, Continuous cell type fate probability curves across organoid maturation. D, RNA program dynamics for early pluripotency, transition neuroectoderm/neuroepithelium and neural maturation programs. E, Cell type fate probability heatmap across model time. F, Heatmap summary of RNA marker program dynamics.

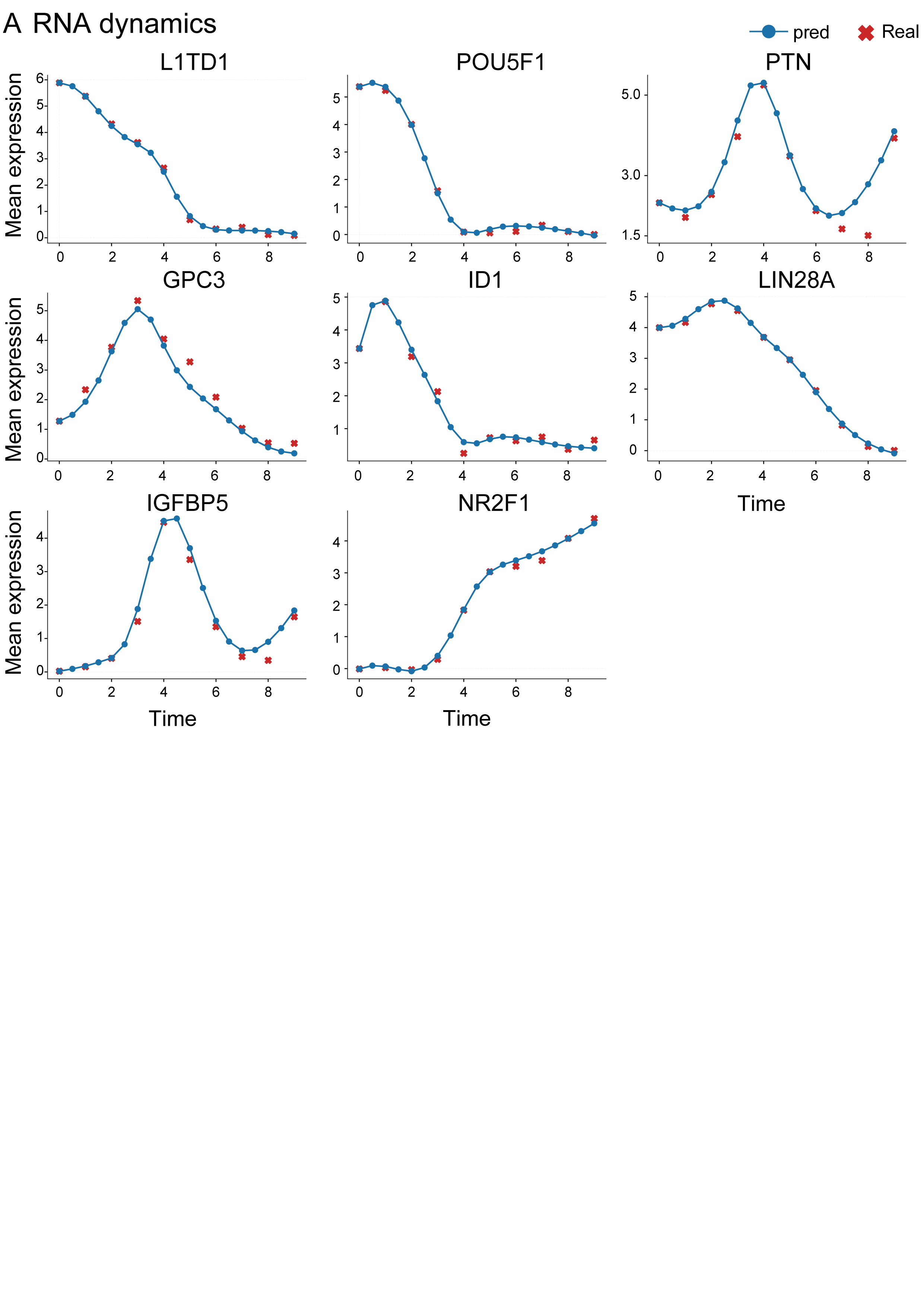

**Supplementary Fig. 11. Human brain organoid RNA marker dynamics.** A, Feature summary along the developmental path for organoid developmental programs.

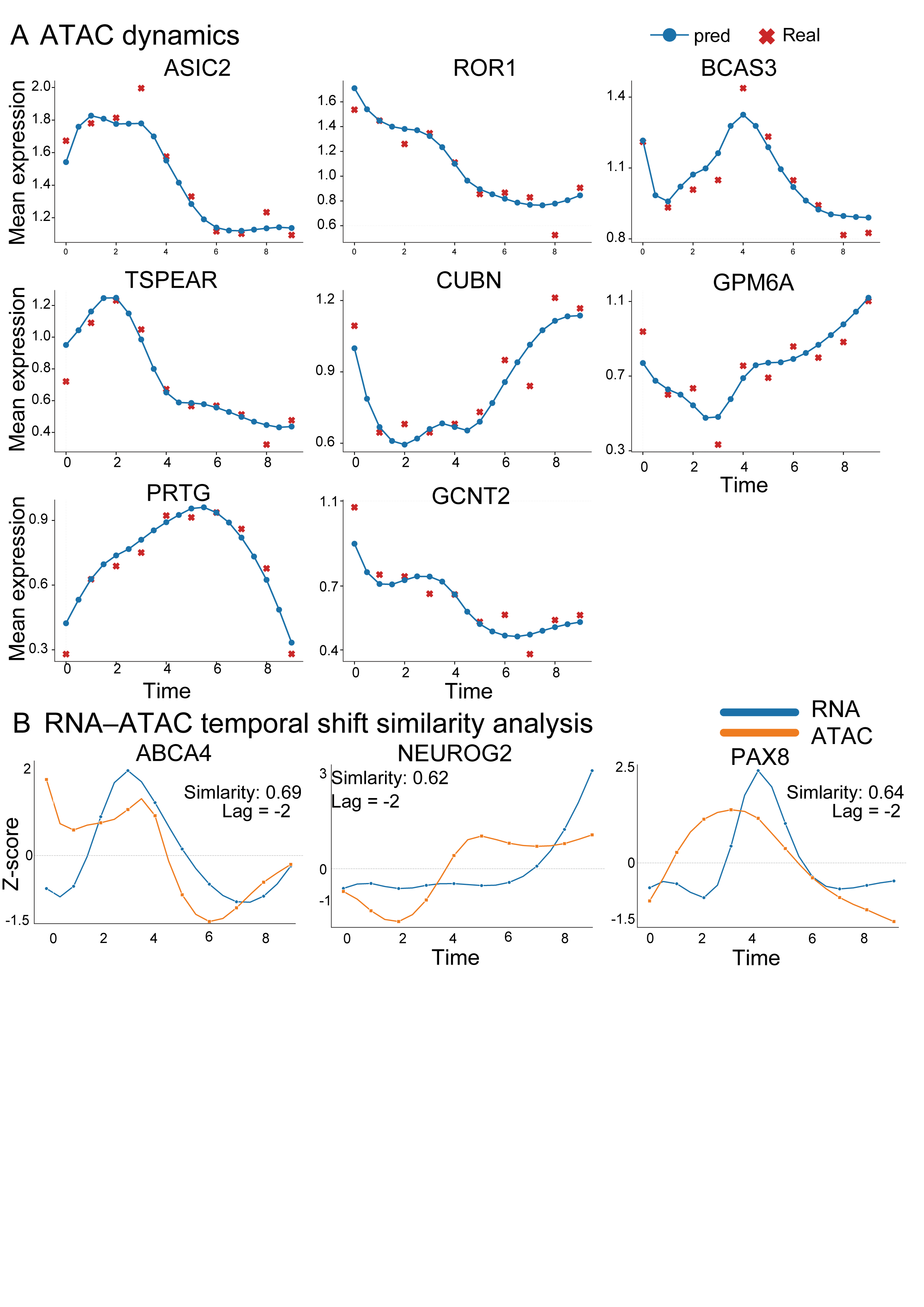

**Supplementary Fig. 12. Human brain organoid ATAC gene activity dynamics and RNA–ATAC temporal shift analysis.** A, ATAC gene activity trajectories reconstructed by MOFI for representative features across continuous model time, with observed measurements shown as anchors. B, RNA–ATAC temporal shift similarity analysis using Spearman correlation over shifts of up to two grid steps, with at least four overlapping time points and 1,000 permutations for each pair.

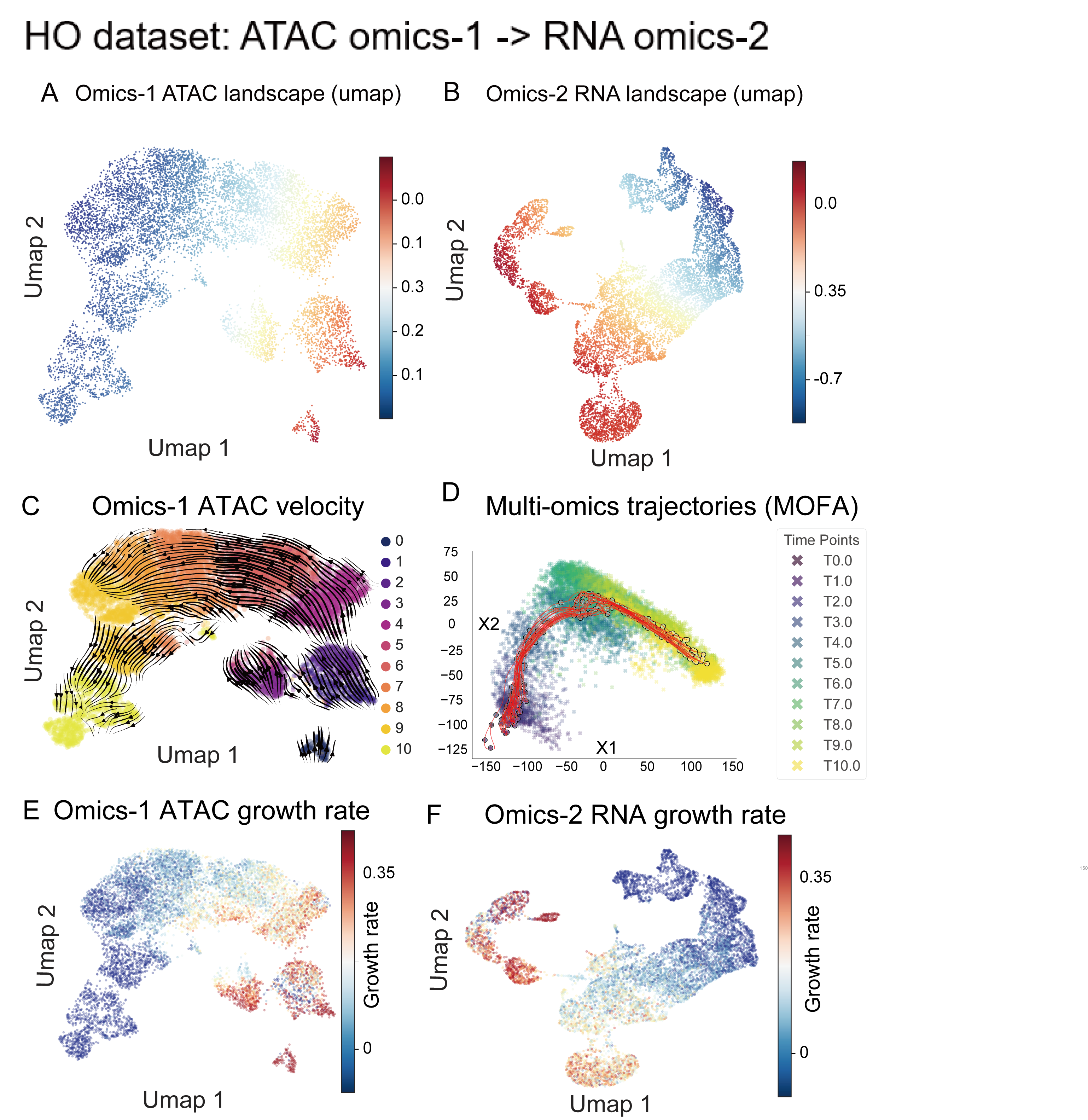

**Supplementary Fig. 13. Reverse analysis of the human brain organoid dataset with ATAC as the primary omics.** A, B, Potential landscapes inferred in the primary ATAC space (A) and the mapped secondary RNA space (B). C, ATAC velocity stream plot inferred by MOFI. D, Joint multi-omics projection showing generated trajectories overlaid on the observed data manifold by MOFA[7]. E, F, growth rate predicted by MOFI in ATAC space (E) and RNA space (F) under reverse direction training.

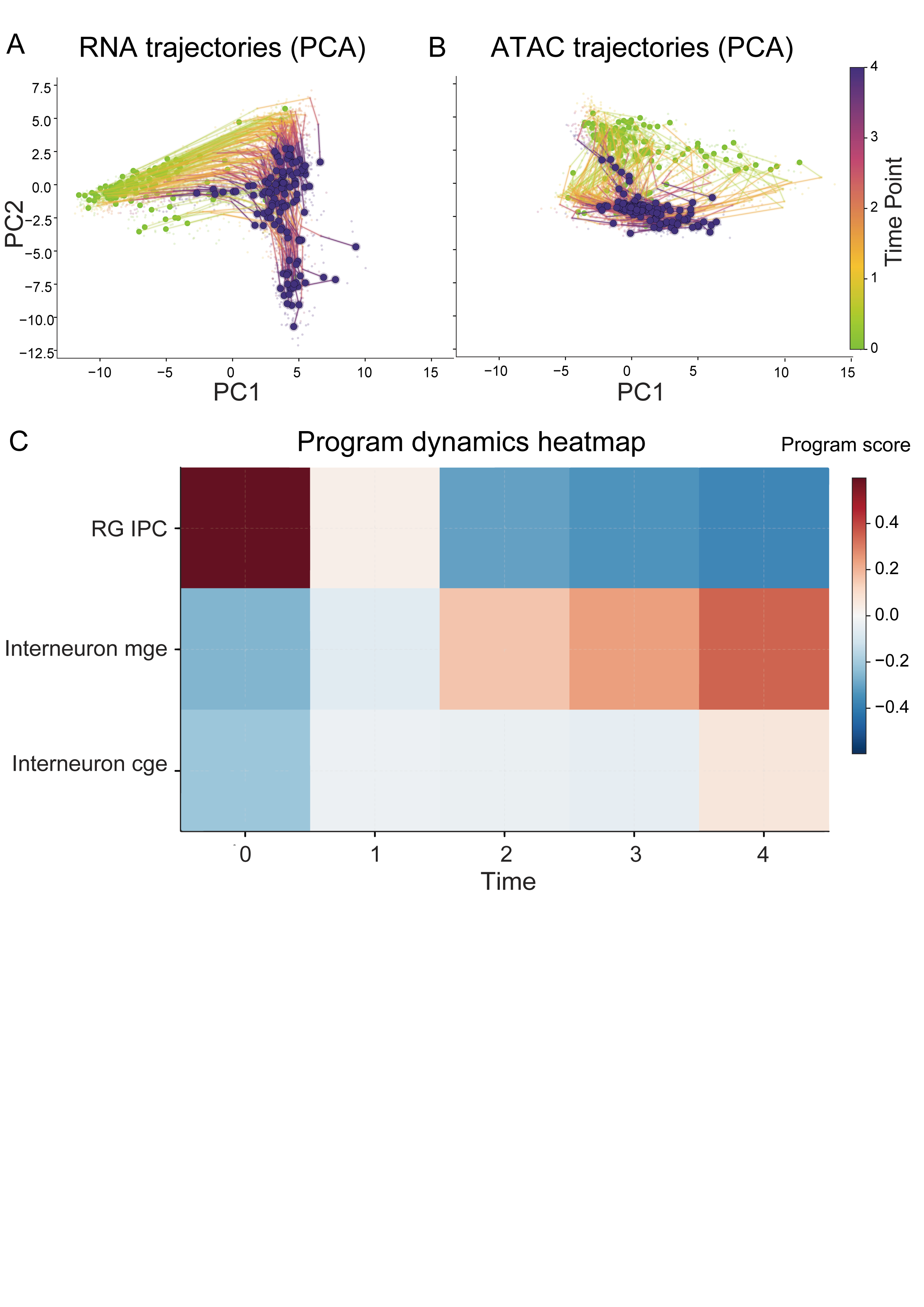
**Supplementary Fig. 14. Developing human cortex RNA to ATAC trajectory reconstruction and program analysis.** A, B, continuous trajectories generated by MOFI for the developing human cortex dataset using RNA as the primary omics (A) and ATAC as the secondary omics (B). C, Program heatmap showing early radial glial/intermediate progenitor activity and later interneuron programs.

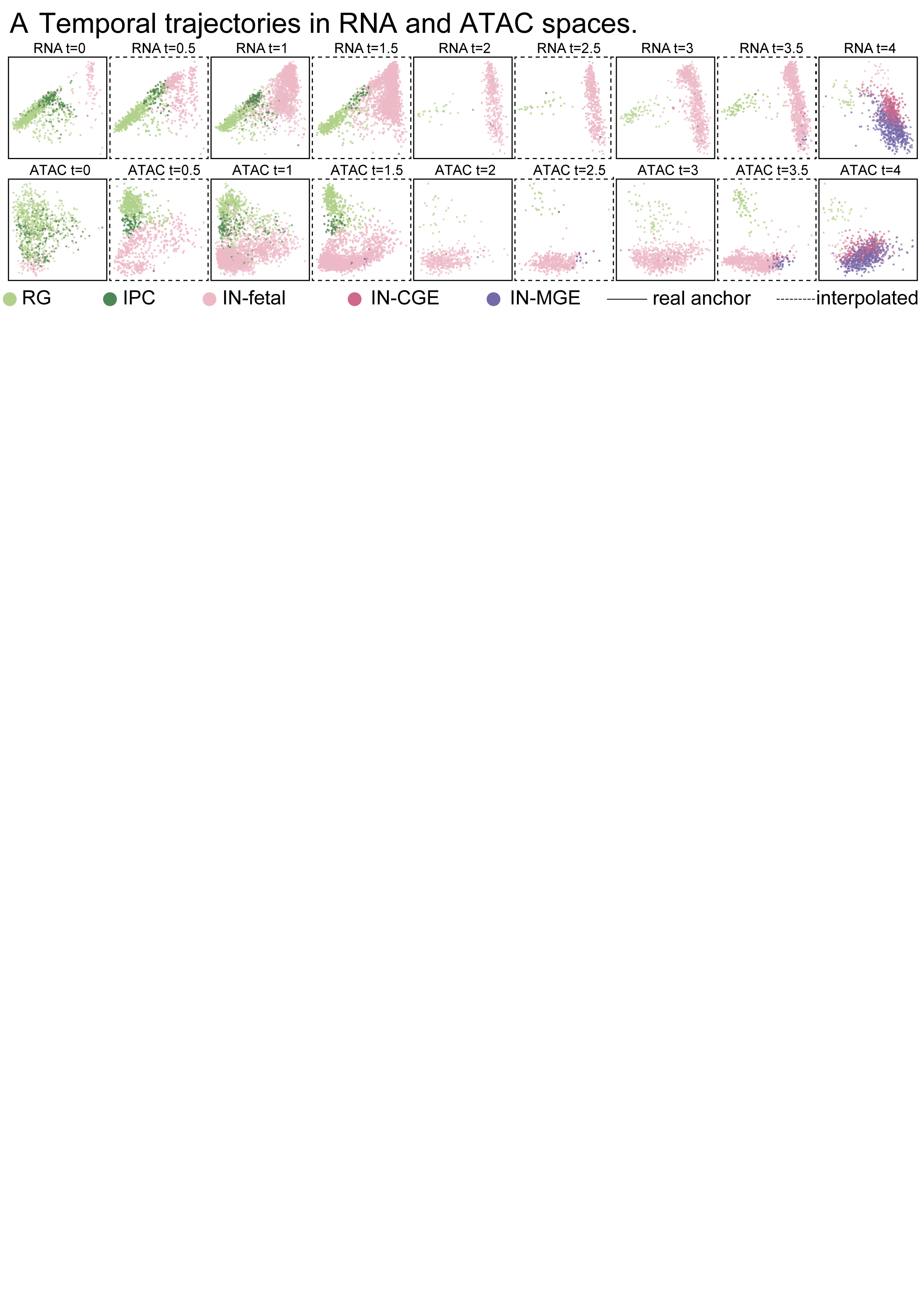

**Supplementary Fig. 15. Temporal cortical trajectories in RNA and ATAC spaces.** A, Temporal PCA panels for RNA and ATAC showing the spatial distribution of observed cells and cells generated by MOFI across developmental time, from early regions enriched for progenitors toward later interneuron states.

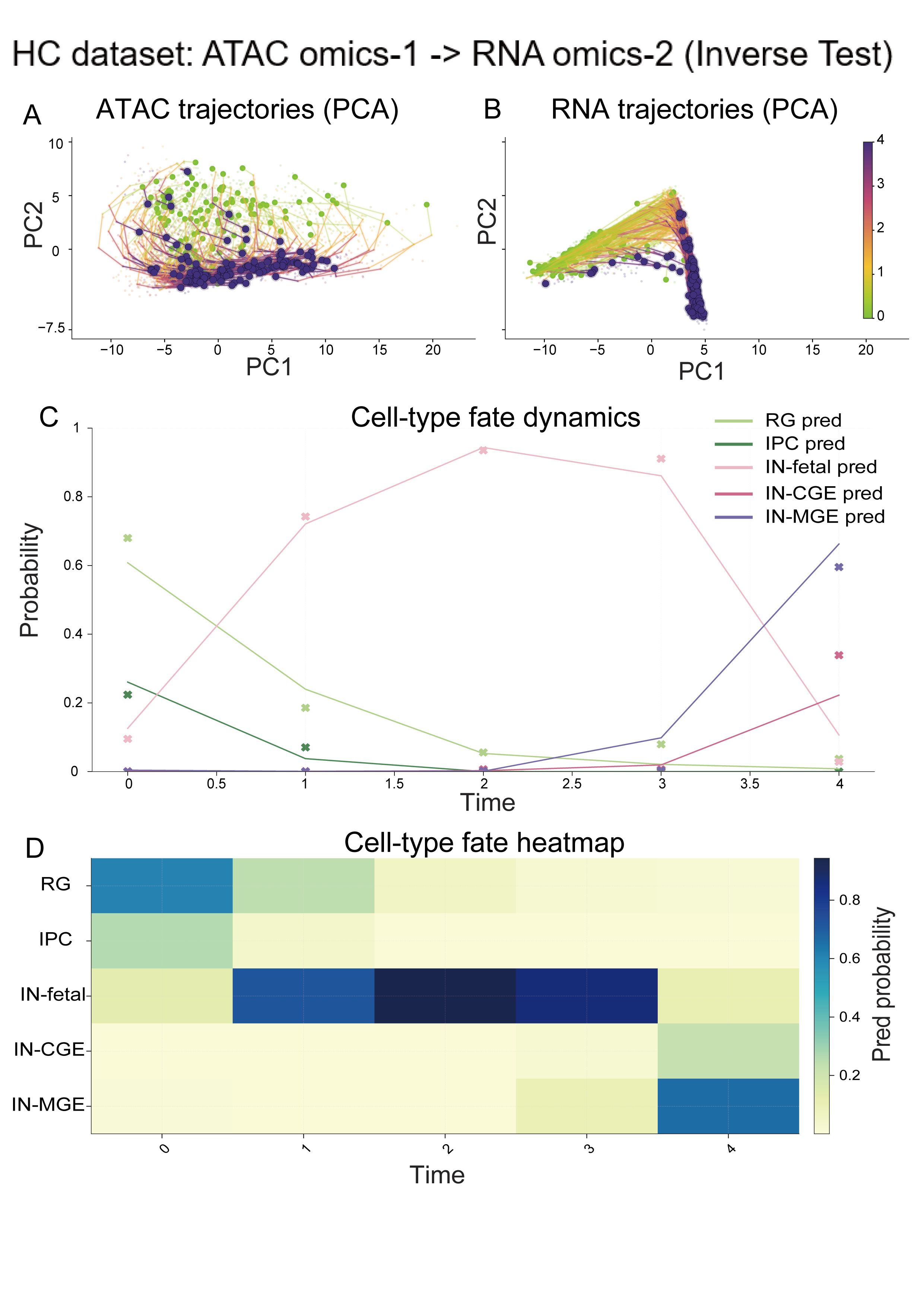

**Supplementary Fig. 16. Reverse developing human cortex analysis with ATAC as the primary omics.** A, B, trajectories generated by MOFI from reverse direction training using ATAC as the primary omics (A) and RNA as secondary omics (B). C, Continuous cell type fate probability curves. D, Cell type fate probability heatmap showing depletion of early progenitors and emergence of later interneuron populations.

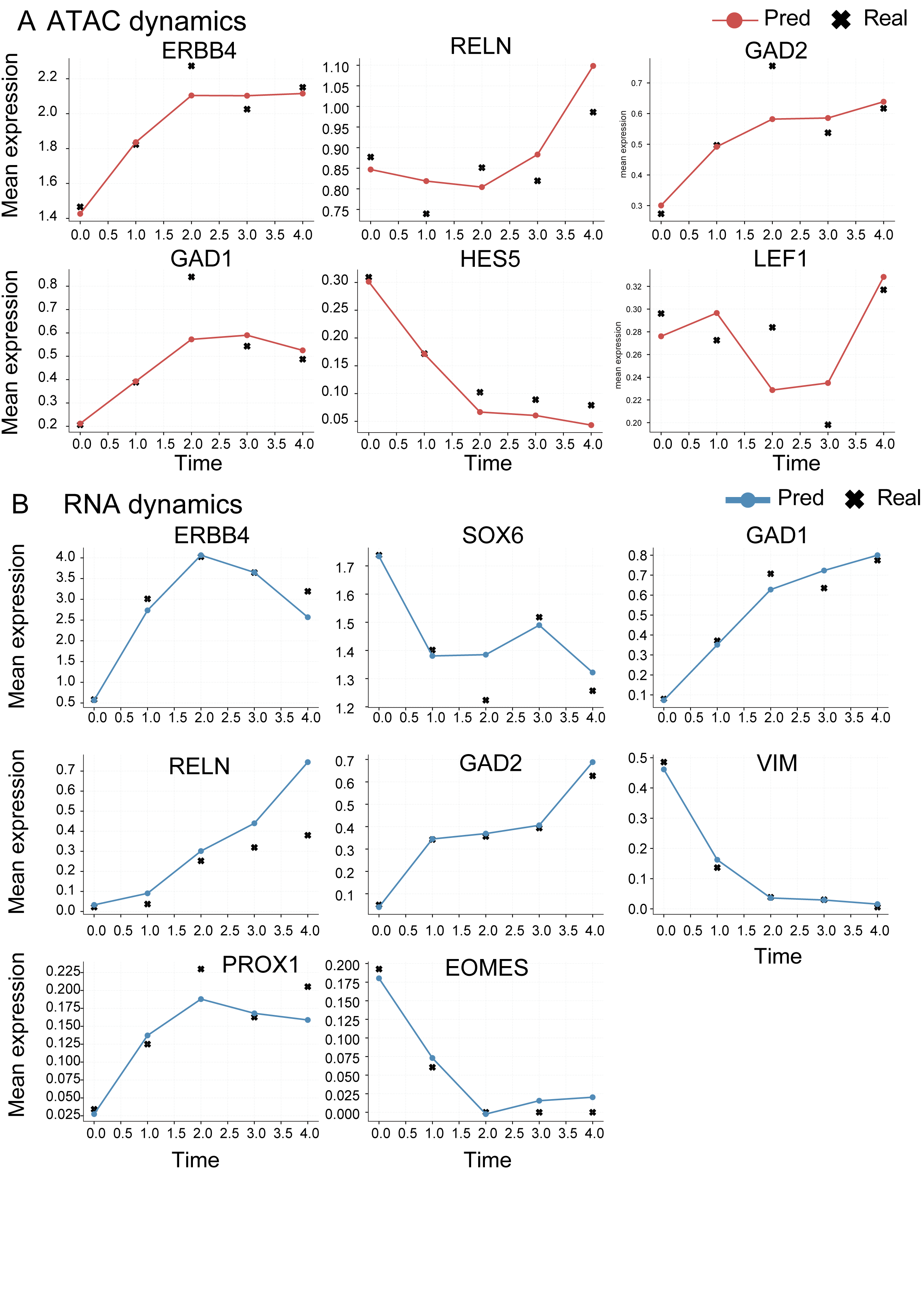

**Supplementary Fig. 17. Developing human cortex marker gene reconstruction.** A, B, marker dynamics reconstructed by MOFI in ATAC gene activity (A) and RNA expression (B). Observed measurements are shown as anchor points. The marker panels include cortical progenitor and interneuron genes such as ERBB4, RELN, GAD1, GAD2, PROX1, LEF1, HES5, SOX6, VIM and EOMES.

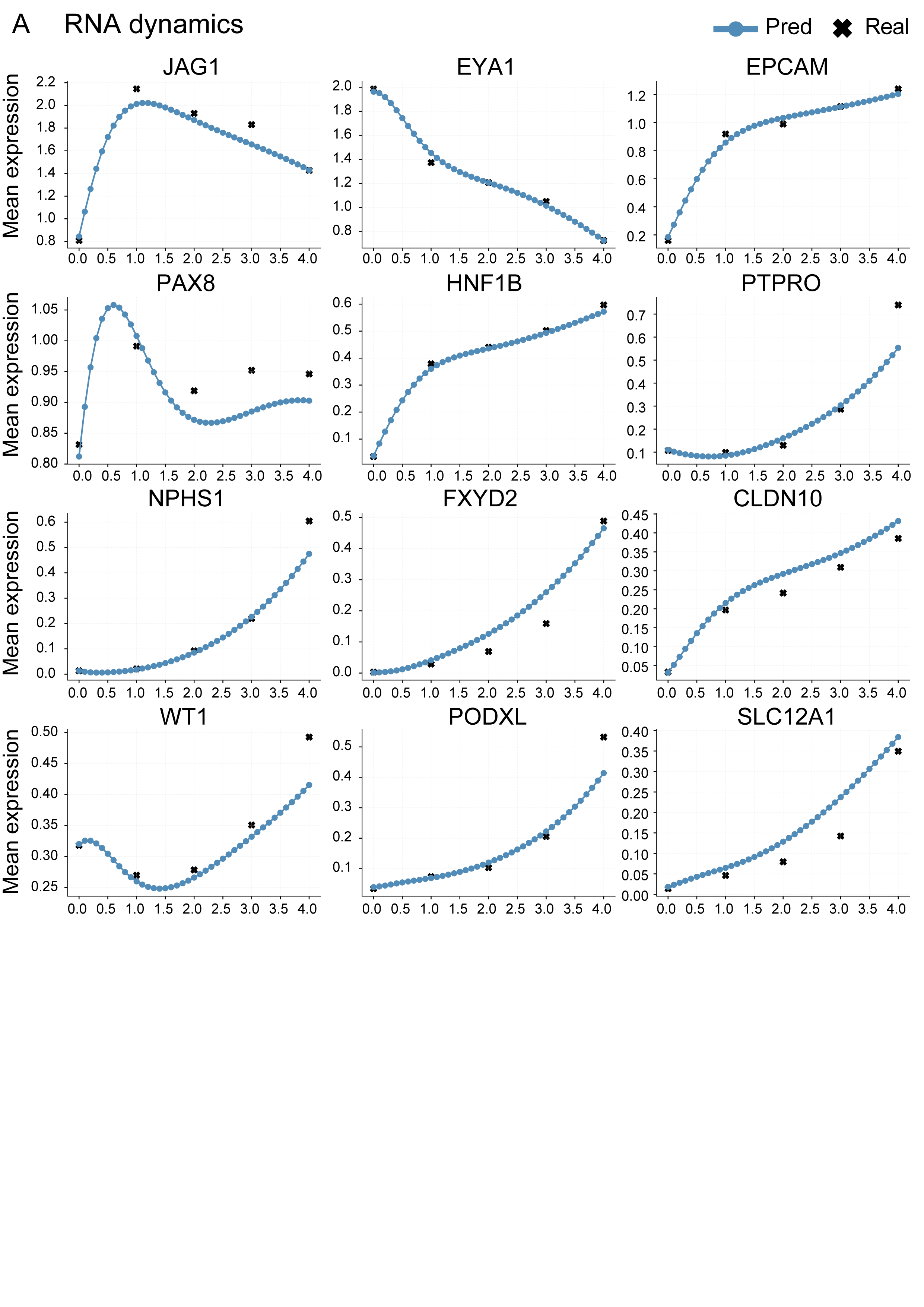

**Supplementary Fig. 18. RNA space trajectory reconstruction for the GSE213152 kidney nephron RNA–ATAC dataset.** A, RNA space molecular trajectories across the ordered temporal observations, with observed measurements shown at sampled time points.

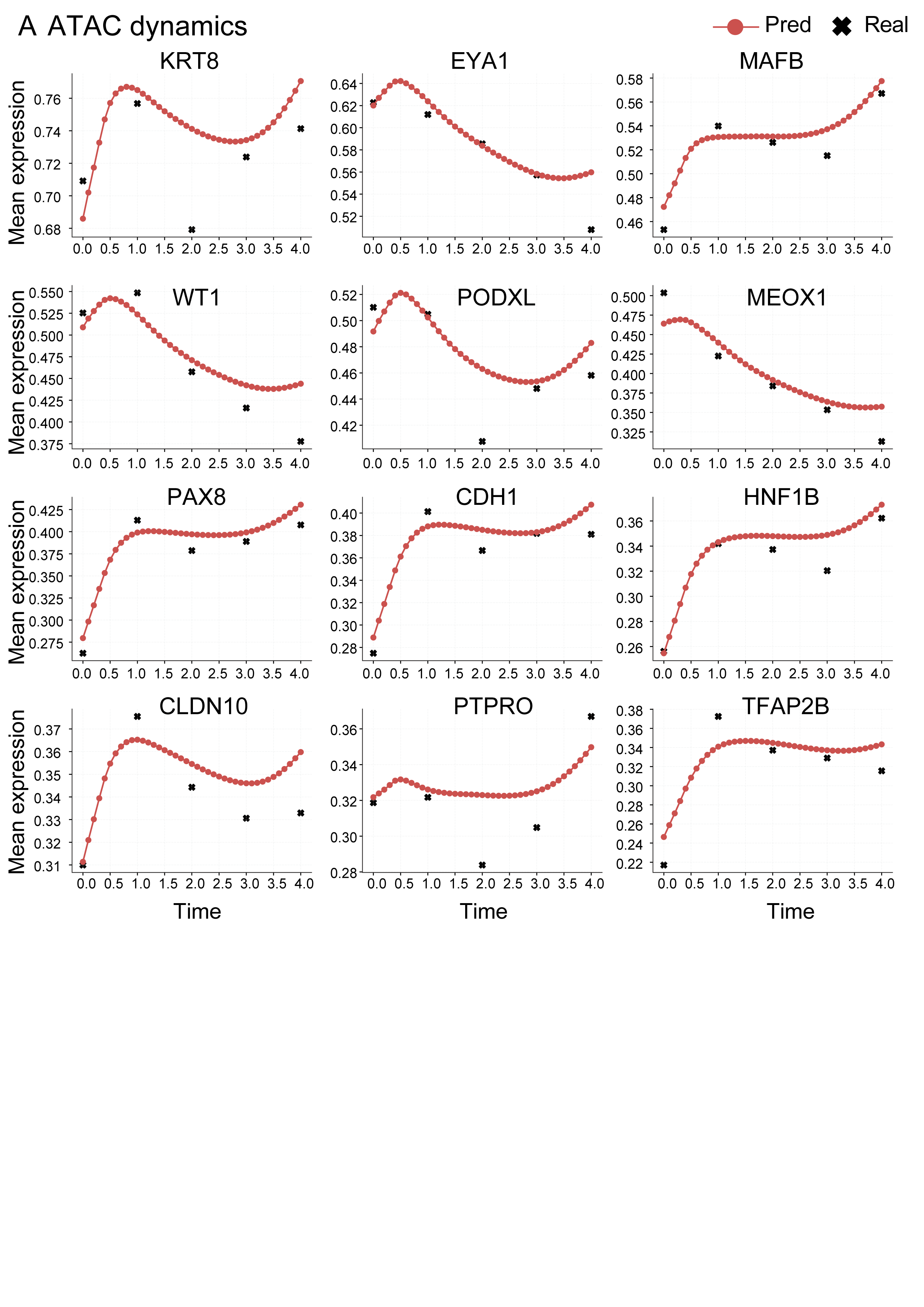

**Supplementary Fig. 19. ATAC space trajectory reconstruction for the GSE213152 kidney nephron RNA–ATAC dataset.** A, ATAC space molecular trajectories across the ordered temporal observations, with observed measurements shown at sampled time points.

**Supplementary Table 1. Reconstruction results for withheld simulated observations**

| **Evaluation setting** | **Withheld during fitting** | **Evaluated output** | **Metric** | **MOFI (mean ± s.d.)** | **Direct concatenation** | **MOSCOT** |
| --- | --- | --- | --- | --- | --- | --- |
| Paired holdout, t = 1 | Omics-1: 0,2,3,4;  omics-2: 0,2,4 | primary t = 1 | *W*_1_^2D^ | 0.1286 ± 0.0046 | 0.1276 ± 0.0097 | 0.1583 |
| Paired holdout, t = 2 | Omics-1: 0,1,3,4;  omics-2: 0,4 | primary t = 2 | *W*_1_^2D^ | 0.1154 ± 0.0182 | 0.1475 ± 0.0343 | 0.2459 |
| Paired holdout, t = 3 | Omics-1: 0,1,2,4;  omics-2: 0,2,4 | primary t = 3 | *W*_1_^2D^ | 0.1282 ± 0.0184 | 0.1608 ± 0.0525 | 0.2457 |
| Paired holdout, t = 1 | Omics-1: 0,2,3,4;  omics-2: 0,2,4 | secondary t = 1 | *W*_1_^3D^ | 0.1405 ± 0.0070 | 0.1304 ± 0.0103 | 0.1612 |
| Paired holdout, t = 2 | Omics-1: 0,1,3,4;  omics-2: 0,4 | secondary t = 2 | *W*_1_^3D^ | 0.1154 ± 0.0182 | 0.1525 ± 0.0330 | 0.5871 |
| Paired holdout, t = 3 | Omics-1: 0,1,2,4;  omics-2: 0,2,4 | secondary t = 3 | *W*_1_^3D^ | 0.1282 ± 0.0184 | 0.1617 ± 0.0521 | 0.2456 |
| primary omics holdout, primary t = 2 | Omics-1: 0,1,3,4;  omics-2: 0,2,4 | primary t = 2 | *W*_1_^2D^ | 0.0882 ± 0.0304 | 0.1557 ± 0.0172 | 0.2459 |
| Secondary omics holdout, secondary t = 1 | Omics-1: 0,1,2,3,4;  omics-2: 0,2,4 | secondary t = 1 | *W*_1_^3D^ | 0.0678 ± 0.0084 | 0.1152 ± 0.0149 | 0.3820 |
| Secondary omics holdout, secondary t = 2 | Omics-1: 0,1,2,3,4;  omics-2: 0,4 | secondary t = 2 | *W*_1_^3D^ | 0.0912 ± 0.0230 | 0.1043 ± 0.0117 | 0.5871 |
| Secondary omics holdout, secondary t = 3 | Omics-1: 0,1,2,3,4;  omics-2: 0,2,4 | secondary t = 3 | *W*_1_^3D^ | 0.1493 ± 0.0244 | 0.1202 ± 0.0113 | 0.4506 |

*MOFI and direct concatenation values are mean ± s.d. over five independent training seeds; MOSCOT is deterministic and is reported as a single value.

**Supplementary Table 2. Real paired multi-omics datasets used for benchmarking and supplementary application**

| **Dataset** | **Omics pair** | **Direction(s)** | **Cell types** | **Time points** |
| --- | --- | --- | --- | --- |
| HSPCs | RNA–protein (CITE-seq) | RNA to protein; protein to RNA | 6 | 4 |
| Human brain organoid (HO) | RNA–ATAC | RNA to ATAC; ATAC to RNA | 4 | 10 |
| Developing human cerebral cortex (HC) | RNA–ATAC | RNA to ATAC; ATAC to RNA | 5 | 5 |
| GSE213152 kidney nephron | RNA–ATAC | RNA to ATAC | 6 | 5 |

**Supplementary Table 3. Mean W1 distances over time for reconstruction across four paired single-cell multi-omics datasets. (It summarizes the results of Supplementary Table 5-12)**

| **Dataset** | **Omics** | **MOFI** | **scMultiNODE** | **Direct concatenation** |
| --- | --- | --- | --- | --- |
| HSPCs | RNA | 6.0725 | 6.7880 | 7.0792 |
| HSPCs | Protein | 12.9678 | 15.0378 | 14.9410 |
| Human brain organoid | RNA | 13.3377 | 13.8435 | 20.0336 |
| Human brain organoid | ATAC | 4.9749 | 5.4237 | 6.3692 |
| Developing human cerebral cortex (HC) | RNA | 6.1566 | 6.0429 | 6.9205 |
| Developing human cerebral cortex (HC) | ATAC | 6.1360 | 6.2195 | 7.0486 |
| GSE213152 kidney nephron | RNA | 6.3490 | 6.8378 | 7.9906 |
| GSE213152 kidney nephron | ATAC | 4.7522 | 4.9904 | 4.7192 |

**Supplementary Table 4. Descriptive summary of pairwise W1 comparisons across fitted real data benchmark points**

| **Comparison set** | **Number of evaluated points** | **MOFI lower than direct concatenation** | **MOFI lower than scMultiNODE** |
| --- | --- | --- | --- |
| Time points after initialization | 40 | 37 / 40 (92.5%) | 33 / 40 (82.5%) |

**Supplementary Table 5. HSPCs RNA W1 distances**

| **Time** | **MOFI W1** | **scMultiNODE W1** | **Direct concatenation W1** |
| --- | --- | --- | --- |
| 1 | 5.3180 | 5.5010 | 6.0687 |
| 2 | 5.5012 | 6.3740 | 6.6642 |
| 3 | 7.3982 | 8.4891 | 8.5048 |

**Supplementary Table 6. HSPCs protein W1 distances**

| **Time** | **MOFI W1** | **scMultiNODE W1** | **Direct concatenation W1** |
| --- | --- | --- | --- |
| 1 | 10.8565 | 12.6638 | 13.0030 |
| 2 | 12.4475 | 13.7547 | 14.6089 |
| 3 | 15.5994 | 18.6950 | 17.2112 |

**Supplementary Table 7. Human brain organoid RNA W1 distances**

| **Time** | **MOFI W1** | **scMultiNODE W1** | **Direct concatenation W1** |
| --- | --- | --- | --- |
| 1 | 8.7833 | 9.9336 | 13.3453 |
| 2 | 10.3656 | 11.4087 | 14.2561 |
| 3 | 10.9525 | 13.1516 | 14.8446 |
| 4 | 13.0758 | 13.2613 | 20.4223 |
| 5 | 13.4699 | 14.6228 | 18.1547 |
| 6 | 14.1469 | 15.0347 | 19.5861 |
| 7 | 14.4463 | 15.5390 | 21.1120 |
| 8 | 16.8535 | 14.9586 | 26.6948 |
| 9 | 17.9457 | 16.6808 | 31.8863 |

**Supplementary Table 8. Human brain organoid ATAC W1 distances**

| **Time** | **MOFI W1** | **scMultiNODE W1** | **Direct concatenation W1** |
| --- | --- | --- | --- |
| 1 | 4.3318 | 4.6913 | 5.6731 |
| 2 | 4.3083 | 4.8482 | 5.4846 |
| 3 | 4.4675 | 5.2016 | 5.5571 |
| 4 | 6.2504 | 6.6314 | 7.7595 |
| 5 | 5.5381 | 5.6882 | 6.9197 |
| 6 | 5.1139 | 5.6019 | 6.2290 |
| 7 | 4.7717 | 5.2523 | 6.0422 |
| 8 | 4.8695 | 5.3046 | 6.3621 |
| 9 | 5.1232 | 5.5934 | 7.2954 |

**Supplementary Table 9. Developing human cerebral cortex (HC) RNA W1 distances**

| **Time** | **MOFI W1** | **scMultiNODE W1** | **Direct concatenation W1** |
| --- | --- | --- | --- |
| 1 | 5.2384 | 5.3748 | 6.4211 |
| 2 | 5.7626 | 5.4778 | 6.3168 |
| 3 | 5.8210 | 5.6417 | 6.5882 |
| 4 | 7.8042 | 7.6774 | 8.3560 |

**Supplementary Table 10. Developing human cerebral cortex (HC) ATAC W1 distances**

| **Time** | **MOFI W1** | **scMultiNODE W1** | **Direct concatenation W1** |
| --- | --- | --- | --- |
| 1 | 6.0004 | 6.2683 | 7.7731 |
| 2 | 5.8790 | 5.6974 | 6.6321 |
| 3 | 6.1146 | 6.0060 | 6.5716 |
| 4 | 6.5498 | 6.9064 | 7.2175 |

**Supplementary Table 11. GSE213152 kidney nephron RNA W1 distances**

| **Time** | **MOFI W1** | **scMultiNODE W1** | **Direct concatenation W1** |
| --- | --- | --- | --- |
| 1 | 6.2544 | 6.8046 | 7.8235 |
| 2 | 6.0012 | 6.4222 | 7.6122 |
| 3 | 6.3813 | 6.6430 | 7.9415 |
| 4 | 6.7589 | 7.4814 | 8.5853 |

**Supplementary Table 12. GSE213152 kidney nephron ATAC W1 distances**

| **Time** | **MOFI W1** | **scMultiNODE W1** | **Direct concatenation W1** |
| --- | --- | --- | --- |
| 1 | 4.3581 | 4.8718 | 4.4657 |
| 2 | 4.9211 | 5.0063 | 4.9033 |
| 3 | 4.9313 | 5.0549 | 4.7900 |
| 4 | 4.7983 | 5.0284 | 4.7178 |

**Supplementary Table 13. Intermediate time point holdout reconstruction results across four real datasets**

| **Dataset** | **Holdout design** | **t** | **Evaluated omics** | **MOFI** | **scMultiNODE** | **Direct concatenation** |
| --- | --- | --- | --- | --- | --- | --- |
| GSE213152 | Paired | 1 | RNA | **7.4816** | 9.6145 | 9.2251 |
| GSE213152 | Paired | 1 | ATAC | **4.5125** | 5.5081 | 4.6724 |
| GSE213152 | Secondary only | 1 | ATAC | **4.3534** | 5.2310 | 4.7050 |
| GSE213152 | Paired | 2 | RNA | **6.3055** | 8.2871 | 8.4170 |
| GSE213152 | Paired | 2 | ATAC | **4.9499** | 5.2841 | 5.0959 |
| GSE213152 | Secondary only | 2 | ATAC | **4.9346** | 5.1110 | 5.1949 |
| GSE213152 | Paired | 3 | RNA | **6.5880** | 8.8614 | 8.8297 |
| GSE213152 | Paired | 3 | ATAC | **4.8849** | 5.4782 | 4.9956 |
| GSE213152 | Secondary only | 3 | ATAC | **4.8927** | 5.7679 | 5.0279 |
| HC | Paired | 1 | RNA | 7.3454 | 7.8692 | **6.5859** |
| HC | Paired | 1 | ATAC | **7.5780** | 8.2597 | 7.8989 |
| HC | Secondary only | 1 | ATAC | **6.1725** | 8.3963 | 7.9848 |
| HC | Paired | 2 | RNA | 7.5502 | 8.3407 | **7.0647** |
| HC | Paired | 2 | ATAC | **6.5807** | 7.7754 | 7.5094 |
| HC | Secondary only | 2 | ATAC | **5.9391** | 7.4717 | 7.7060 |
| HC | Paired | 3 | RNA | **7.5231** | 7.9394 | 8.3038 |
| HC | Paired | 3 | ATAC | **6.7358** | 7.1502 | 7.8909 |
| HC | Secondary only | 3 | ATAC | **6.2077** | 7.0095 | 7.9930 |
| Human brain organoid | Paired | 1 | RNA | **13.3229** | 20.1435 | 17.5141 |
| Human brain organoid | Paired | 1 | ATAC | **5.0052** | 6.8533 | 6.0677 |
| Human brain organoid | Secondary only | 1 | ATAC | **4.3157** | 7.1794 | 6.1367 |
| Human brain organoid | Paired | 2 | RNA | **15.4366** | 20.0699 | 20.7438 |
| Human brain organoid | Paired | 2 | ATAC | **5.2217** | 7.4975 | 7.6377 |
| Human brain organoid | Secondary only | 2 | ATAC | **4.2822** | 7.7513 | 7.3969 |
| Human brain organoid | Paired | 3 | RNA | **18.5194** | 22.8010 | 19.7890 |
| Human brain organoid | Paired | 3 | ATAC | **6.4032** | 8.0968 | 7.7406 |
| Human brain organoid | Secondary only | 3 | ATAC | **4.3877** | 8.5328 | 7.7673 |
| Human brain organoid | Paired | 4 | RNA | **21.9107** | 22.4640 | 22.8584 |
| Human brain organoid | Paired | 4 | ATAC | **7.4626** | 8.3965 | 8.1245 |
| Human brain organoid | Secondary only | 4 | ATAC | **6.2138** | 8.8196 | 8.9542 |
| Human brain organoid | Paired | 5 | RNA | **16.6915** | 22.3061 | 20.1841 |
| Human brain organoid | Paired | 5 | ATAC | **6.2239** | 7.1655 | 7.4897 |
| Human brain organoid | Secondary only | 5 | ATAC | **5.4607** | 7.0213 | 8.0951 |
| Human brain organoid | Paired | 6 | RNA | **14.6014** | 21.5885 | 20.1895 |
| Human brain organoid | Paired | 6 | ATAC | **5.1970** | 6.4725 | 6.7798 |
| Human brain organoid | Secondary only | 6 | ATAC | **5.1359** | 6.6049 | 6.7853 |
| Human brain organoid | Paired | 7 | RNA | **15.0807** | 20.3981 | 20.4407 |
| Human brain organoid | Paired | 7 | ATAC | **4.8849** | 6.4078 | 6.9924 |
| Human brain organoid | Secondary only | 7 | ATAC | **4.7686** | 6.7774 | 6.6114 |
| Human brain organoid | Paired | 8 | RNA | **17.1950** | 21.4900 | 24.2116 |
| Human brain organoid | Paired | 8 | ATAC | **5.1307** | 6.1528 | 6.9667 |
| Human brain organoid | Secondary only | 8 | ATAC | **4.7122** | 6.8075 | 7.1765 |
| HSPCs | Paired | 1 | RNA | **5.5433** | 8.3059 | 6.5300 |
| HSPCs | Paired | 1 | Protein | **11.1621** | 19.0841 | 13.4818 |
| HSPCs | Secondary only | 1 | Protein | **10.8793** | 19.4627 | 13.3109 |
| HSPCs | Paired | 2 | RNA | **6.0981** | 9.7914 | 7.4876 |
| HSPCs | Paired | 2 | Protein | **13.4588** | 20.0352 | 15.7718 |
| HSPCs | Secondary only | 2 | Protein | **12.8255** | 20.1023 | 16.4656 |

Note: Each row reports W1 at the omitted intermediate time point for seed 42. “Paired” removes both modalities at the target time and evaluates each withheld modality separately. “Secondary only” removes only the secondary modality while retaining its paired primary observation. Lower W1 indicates closer reconstruction. Bold values mark the lowest W1 among the three models in each row.

**Supplementary Table 14. Recommended starting configuration for MOFI dynamics training**

| **Setting** | **Recommended starting value** |
| --- | --- |
| Model components | velocity, growth |
| PCA dimensions | 50 |
| Map between omics | Spliced autoencoder mapper; reported starting hidden dimension 32; mapper fixed during standard dynamics fitting unless update_transport is enabled |
| Velocity network | hidden dimension 400; 2 layers; residual = false |
| Growth network | hidden dimension 400; 2 layers; residual = false |
| Optimizer learning rate | 1 × 10⁻⁴ |
| Batch size | 400 |
| Stage 1: primary space initialization | 100 epochs; $\lambda_{\text{OT},1}$ = 1; $\lambda_{\text{mass}}$ = 0.1; $\lambda_{Engery}$ = 0 |
| Stage 2: multi-omics refinement | 200 epochs; $\lambda_{\text{OT},1}$ = 1; $\lambda_{\text{mass}}$ = 1.0; $\lambda_{Engery}$ = 0.01; $\lambda_{\text{OT},2}$ = 1 |
| Random seed used for reported MOFI real data runs | 42 |

**Supplementary Table 15. Settings used for each trained MOFI dynamics model**

| **Dataset** | **Paired cells** | **Map setting** | **Actual training schedule** |
| --- | --- | --- | --- |
| Simulation | 2452 | T is set as  $\left[ X,Y,\exp(-\frac{\left( X-2 \right)^{2}}{2\times0.05}-\frac{\left( Y-1.5 \right)^{2}}{2\times0.05} \right]$ | Stage 1: 200 epochs  $\lambda_{\text{OT},1}:$1, $\lambda_{\text{mass}}:$ 0.1, $\lambda_{\text{energy}}:$ 0.05, $\lambda_{\text{OT},2}: 0;$  Stage 2: 400 epochs  $\lambda_{\text{OT},1}:$1, $\lambda_{\text{mass}}:$ 0.02, $\lambda_{\text{energy}}:$ 0.5, $\lambda_{\text{OT},2}: 0;$  (Sync-energy considers omics-2) |
| HSPCs | 28,087 | $\alpha_{\mathrm{same}}:1 ,\alpha_{\mathrm{cross}}:40$  $\alpha_{c\mathrm{onsistency}}:45$ | Stage 1: 180 epochs  $\lambda_{\text{OT},1}:$1.2, $\lambda_{\text{mass}}:$ 1, $\lambda_{\text{energy}}:$ 0, $\lambda_{\text{OT},2}: 0;$  Stage 2: 40 epochs  $\lambda_{\text{OT},1}:$ 0.2, $\lambda_{\text{mass}}:$ 8, $\lambda_{\text{energy}}:$ 5, $\lambda_{\text{OT},2}: 1;$ |
| Human brain organoid | 9,894 | $\alpha_{\mathrm{same}}:1 ,\alpha_{\mathrm{cross}}:20$  $\alpha_{c\mathrm{onsistency}}:100$ | Stage 1: 150 epochs  $\lambda_{\text{OT},1}:$1, $\lambda_{\text{mass}}:$ 0.1, $\lambda_{\text{energy}}:$ 0, $\lambda_{\text{OT},2}: 0;$  Stage 2: 200 epochs  $\lambda_{\text{OT},1}:$ 1, $\lambda_{\text{mass}}:$ 10, $\lambda_{\text{energy}}:$ 0.01, $\lambda_{\text{OT},2}: 1;$ |
| Developing human cerebral cortex (HC) | 6,163 | $\alpha_{\mathrm{same}}:1 ,\alpha_{\mathrm{cross}}:20$  $\alpha_{c\mathrm{onsistency}}:1$ | Stage 1:100 epochs  $\lambda_{\text{OT},1}:$1, $\lambda_{\text{mass}}:$ 4 $\lambda_{\text{energy}}:$ 0, $\lambda_{\text{OT},2}: 0;$  Stage 2a:100 epochs  $\lambda_{\text{OT},1}:$1, $\lambda_{\text{mass}}:$ 2 $\lambda_{\text{energy}}:$ 1, $\lambda_{\text{OT},2}: 1;$  Stage 2b:200 epochs  $\lambda_{\text{OT},1}:$ 1, $\lambda_{\text{mass}}:$ 1, $\lambda_{\text{energy}}:$ 0.01, $\lambda_{\text{OT},2}: 1;$ |
| GSE213152 kidney nephron | 60,845 | $\alpha_{\mathrm{same}}:1 ,\alpha_{\mathrm{cross}}:20$  $\alpha_{c\mathrm{onsistency}}:100$ | Stage 1: 100 epochs  $\lambda_{\text{OT},1}:$1, $\lambda_{\text{mass}}:$ 0.1 $\lambda_{\text{energy}}:$ 0, $\lambda_{\text{OT},2}: 0;$  Stage 2: 200 epochs  $\lambda_{\text{OT},1}:$ 1, $\lambda_{\text{mass}}:$ 1, $\lambda_{\text{energy}}:$ 0.01, $\lambda_{\text{OT},2}: 1;$ |

Note: $\alpha_{\mathrm{multi}}$ is set as 0.1.

**Supplementary Table 16. Predefined marker sets used for HSPCs perturbation analyses**

| **Target group** | **Marker set annotation** | **Genes** |
| --- | --- | --- |
| Ery_core_1 | Erythroid marker set 1 | KLF1, GATA1, TFRC |
| Mk_core_1 | Megakaryocyte marker set 1 | ITGA2B, GP1BA |
| Mk_core_2 | Megakaryocyte marker set 2 | PF4, PPBP |
| Mk_union | Combined megakaryocyte marker set | ITGA2B, GP1BA, PF4, PPBP |
| Mas_core_1 | Mast cell marker set 1 | TPSAB1, TPSB2, CPA3 |
| Mas_core_2 | Mast cell marker set 2 | MS4A2, FCER1A, IL3RA |
| Mas_union | Combined Mas marker set | MS4A2, FCER1A, IL3RA, TPSAB1, TPSB2, CPA3 |

Supplementary References

[1] Peng, Q., Zhou, P. & Li, T. stVCR: spatiotemporal dynamics of single cells. Nat. Methods 23, 542–553 (2026).
[2] Klein, D. et al. Mapping cells through time and space with moscot. Nature 638, 1065–1075 (2025).
[3] Zhang, J., Chakravarthy, M. & Singh, R. scMultiNODE: Integrative and Scalable Framework for Multi-Modal Temporal Single-Cell Data. In Proc. 16th ACM International Conference on Bioinformatics, Computational Biology and Health Informatics, Article 69 (ACM, 2025).
[4] Cang, Z., Wang, J., Wei, X. & Zhao, Y. Synchronization of unbalanced dynamical optimal transport across multiple spaces. Preprint at arXiv:2602.18725 (2026).
[5] Chen, R. T. Q., Rubanova, Y., Bettencourt, J. & Duvenaud, D. Neural ordinary differential equations. In Advances in Neural Information Processing Systems 31, 6572–6583 (NeurIPS, 2018).
[6] Fleck, J. S. et al. Inferring and perturbing cell fate regulomes in human brain organoids. Nature 621, 365–372 (2023).
[7] Argelaguet, R. et al. Multi-Omics Factor Analysis—a framework for unsupervised integration of multi-omics data sets. Mol. Syst. Biol. 14, e8124 (2018).
